# Identification of eukaryotes as a bycatch in global microbiome data

**DOI:** 10.64898/2026.11.21.765608

**Authors:** Oleksandr M. Maistrenko, Chan Yeong Kim, Suguru Nishijima, Anthony Fullam, Pamela Ferretti, Christian Schudoma, Marisa Isabell Keller, Shahriyar Mahdi Robbani, Ece Kartal, Wasiu Akanni, Renato Alves, Thea Van Rossum, Supriya Khedkar, Daniel Podlesny, Yasushi Kojima, Takashi Hamabata, Naomi Uemura, Naoyoshi Nagata, Anja Spang, Anna Heintz-Buschart, Thomas Sebastian B. Schmidt, Michael Kuhn, Peer Bork

## Abstract

Eukaryotes shape microbiomes as macroscopic hosts, microscopic community members, and food sources, yet the diversity and ecological significance of microbial eukaryotes remain poorly resolved. Here, we present metaEukaryome, an organelle genome-centered framework for detection of eukaryotes in metagenomic data that leverages mitochondrial and plastid sequences as robust signatures of presence. Applying metaEukaryome to 75,727 metagenomic samples collected across the globe reveals strong environment-specific patterns in the prevalence of detected eukaryotes. Integrating eukaryotic and prokaryotic profiles further reveals recurrent cross-domain co-occurrence and symbiosis in the human gut. In particular, *Blastocystis* is associated with metabolically reduced, uncultured bacterial clades, including *Candidatus* Borkfalkia and CAG-345, whose DNA and RNA are detected in a *Blastocystis* culture previously reported as axenic. Their complementary metabolic repertoires further support a candidate symbiotic relationship. These findings imply that novel (endo-)symbionts of gut protists can be discovered by reanalysing existing data.

## Introduction

Eukaryotes (plants, fungi, animals, and protists) comprise essential community-organizing species across all ecosystems^1–5^. They occupy every trophic level of food webs, as primary producers, decomposers of dead organic matter, parasites or other symbiont types, and predators up to the apex at both microscopic and macroscopic scales. While many studies perform surveillance of the composition of ecosystems^6–10^, there is a limited understanding of the cross-domain interactions. In microbiome research, amplicon sequencing has been adapted to estimate prevalences and abundances of eukaryotes alongside prokaryotes, which enables holistic characterization of ecosystems’ functioning. Various marker genes have been targeted to assess diet and eukaryotic community composition, including protist/protozoan parasites, fungi and dietary plants/animals in human and animal-associated microbiomes, as well as directly in environmental samples^11–17^. However, these approaches are limited by the primers used and thus restrict the range of species that can be detected. In contrast, untargeted whole-metagenome shotgun (WMS) sequencing in principle allows for the simultaneous detection of all groups of organisms (prokaryotes, eukaryotes and viruses) present in a sample. The vast majority of metagenomic data analysis focuses on prokaryotes while eukaryotes are understudied due to computational challenges and other reasons^1,18–26^. The main reasons for exclusion of eukaryotes from analyses are often their low abundance, lack of reference genomes or bacterial contamination of existing ones, large size and high complexity of nuclear genomes. Recently, multiple studies have started to tackle these technical issues and gaps in knowledge by developing approaches and pipelines for detecting eukaryotic components in metagenomic samples^27–49^ (**Table S1**). Previous studies have shown that, for example, there is an enrichment of prokaryotic taxa in *Blastocystis*-positive human gut samples, and that carriage of this protist is associated with better cardiovascular health^39,50,51^. Indeed, even dietary signatures can be inferred from WMS data in fecal paleo-samples^52,53^ using gene-based profiling or in contemporary samples and potentially used to model metabolic contribution of diet^48^. In environmental WMS samples eukaryotes are also abundant and diverse, and it is possible to recover eukaryotic bins via assembly^54^. Taken together, this strongly suggests that prokaryotic and eukaryotic communities need to be considered in conjunction, given their mutual ecological impact^1^. Yet, only a few studies systematically include eukaryotes in the interpretation of metagenomic data^50,52,53,55–58^. Moreover, many studies have focused on a specific group of eukaryotes, for example, parasites/commensals^50,51^, food species^48^, or specific taxonomic groups (e.g., fungi^59^) or analyzed a limited set of samples. Thus, we currently have a very limited understanding of eukaryotic diversity across the global microbiome, including their role in human and other animal microbiomes, which hampers efforts to resolve the ecological role of eukaryotes and their connection to and impact on the prokaryotic fraction of the community.

Here we introduce metaEukaryome, which targets 11,932 mitochondrial and/or plastid species clusters and incorporates the Protist Ribosomal Reference (PR2) database of 18S rRNA sequences to detect additional protists lacking organelle genomes. The advantage of this approach lies in the high copy number and conserved structure of eukaryotic organelle genomes, which makes it possible to assess the diversity of eukaryotes in metagenomic sequencing data. Moreover, mitochondrial and plastid genome sequences are available for many taxa for which nuclear reference genomes remain unavailable, thereby expanding the taxonomic breadth of eukaryotes that can be detected and profiled. We comprehensively profiled eukaryotic life across the SPIRE metagenomic database (n = 75,727)^60^ with curated metadata in Metalog^61^. In ocean and soil samples, we observed known patterns of cosmopolitan distribution of multiple algae and fungal species, respectively. In human and animal gut microbiomes, metaEukaryome robustly identified parasitic worms, fungi, protists, and diet species. In addition to eukaryotic patterns related to age groups, income levels and geography in the human gut microbiome, we found that many symbiotic (*sensu* de Bary^62^) eukaryotes, as hypothesized^21^, were associated with increased alpha diversity of the prokaryotic community (with several exceptions in which eukaryotes were associated with decreased alpha diversity). We also discovered and functionally characterized novel associations between individual (diet or symbiotic) eukaryotic species and prokaryotic species in the human gut microbiome, including multiple putative cases of (endo-)symbioses between gut protists (e.g., *Blastocystis*, *Entamoeba*, *Dientamoeba*, and *Endolimax*) and uncultured small-genome prokaryotes.

## Results

### metaEukaryome accurately detects eukaryotes in metagenomic data

We developed the metaEukaryome pipeline to detect eukaryotes across Earth’s microbiomes. metaEukaryome identifies eukaryotic organelle DNA, specifically mitochondrial DNA (mtDNA) and plastid DNA (ptDNA), from WMS sequencing data. In addition, to capture protists that lack mtDNA and ptDNA (due to an anaerobic lifestyle and/or lack of sequenced organelle genomes), such as *Monocercomonoides exilis*^63^ and *Entamoeba histolytica*^64^, we also incorporated 18S rRNA sequences from the PR2 database^65^. In total, metaEukaryome enables the detection of 11,932 eukaryotic species clusters based on plastid and mitochondrial genomes (corresponding to the largest set of profileable species among similar tools - 16,737 species/strains with names on NCBI Taxonomy) and an additional 346 species clusters based on the PR2 database, which are absent from mitochondrial and plastid genome databases (**Table S2**, **Figure 1A**, **Methods**). To benchmark the accuracy of metaEukaryome, we applied it to metagenomic samples for which eukaryote presence had been independently assessed by non-metagenomic approaches, including qPCR (98 samples for helminths and 68 samples for *Blastocystis*)^66,67^, microscopic examination (351 samples for helminths)^68^, and ITS sequencing (122 samples for fungi)^69^, and compared the performance with existing metagenomic tools for eukaryote detection (i.e., taxaTarget, CORRAL, EukDetect, and MetaPhlAn4)^37–41^. To enable direct comparison with the reference data, profiler outputs were binarized into presence/absence calls and evaluated using precision, recall, and F1 score (**Methods**). metaEukaryome outperformed the other publicly available methods (F1 score increase ≥ 0.3 in all cases) in detecting parasitic nematodes, including *Ascaris*, *Necator*, and *Ancylostoma* (**Figure S1A–S1C**). This may reflect distinctive features of these helminths, which produce large numbers of eggs^70,71^, each containing particularly high mitochondrial content and therefore a higher mtDNA-to-chromosomal DNA ratio^72^. Beyond parasitic worms, metaEukaryome achieved performance comparable to the best-performing publicly available tools (F1 score difference ≤ 0.01 in all cases) for detecting *Blastocystis* and fungi (*Candida* and *Saccharomyces*) (**Figure S1D–S1F**). Together, these results show that metaEukaryome detects diverse eukaryotes in metagenomes and expands the scope of metagenomic profiling across a broader range of eukaryotic organisms.

**Figure 1.**
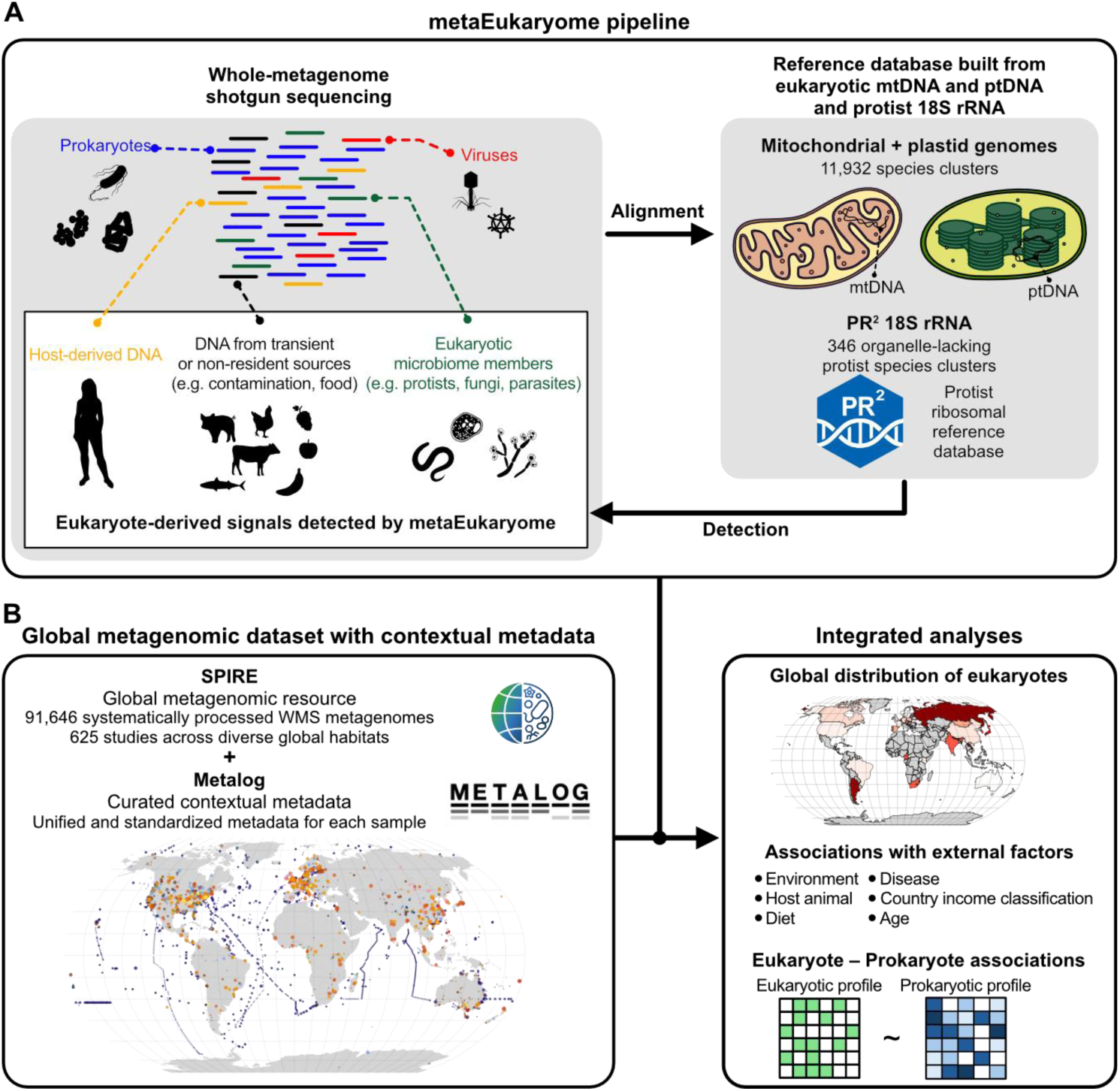
Overview of the metaEukaryome pipeline and global applications. (**A**) Whole-metagenome shotgun (WMS) sequencing captures DNA derived from multiple sources. Using human fecal WMS data as an example, most reads originate from prokaryotes, which make up the majority of the gut microbiome, but reads can also derive from viruses, host DNA, dietary material, contamination introduced during sampling, and eukaryotic microorganisms inhabiting the gut. metaEukaryome detects mitochondrial DNA (mtDNA), plastid DNA (ptDNA), and protist 18S rRNA from these datasets, thereby enabling detection of host-derived DNA, diet-derived DNA, and eukaryotic symbionts. (**B**) In this study, we applied metaEukaryome to 75,727 systematically processed WMS metagenomes from 578 studies in SPIRE, spanning diverse environments worldwide, and integrated the corresponding contextual metadata curated in Metalog. These combined data were used to characterize the global distribution of eukaryotes and to examine associations with environmental and host-related factors, including diet, disease, and country income classification, as well as associations between eukaryotic and prokaryotic profiles in the human gut.

metaEukaryome is also designed to detect food-derived eukaryotic species in the fecal samples. We analyzed data from the Japanese 4D cohort to verify that the detected diet-derived eukaryotic signals reflect actual food intake. For this cohort, both gut metagenomes and dietary questionnaire data are available (n = 4,198, **Methods**)^73^. We found concordance between detected eukaryotic organisms and multiple entries in the dietary questionnaire (**Figure S1G**). For example, the detection of *Gallus* (i.e., chicken) in metagenomes was positively associated with both egg and meat consumption, and *Sus* (i.e., pork) with meat intake (0.05 ≤ *P* < 0.1). In addition, the genus *Saccharomyces*, which includes baker’s yeast in bread, beer, and wine, was positively associated with bread consumption, while *Kazachstania*, a yeast frequently detected in fermented foods^74^, showed positive associations with the consumption of fermented products such as kimchi and Japanese pickles (*P* < 0.05). These associations support the validity of our framework and suggest that dietary intake can be partially reconstructed from eukaryotic signatures in metagenomic data. However, not all expected associations were observed (**Figure S1G**), possibly due to the low amount of remaining dietary DNA in fecal samples and the limited resolution of dietary questionnaires in capturing the frequency, quantity, and preparation methods of food intake.

### Global distributions of eukaryotes across environmental microbiomes

We applied metaEukaryome to 75,727 whole-metagenome shotgun datasets from the SPIRE database^60^, spanning diverse environments worldwide, and integrated the corresponding curated contextual data retrieved from Metalog^61^ (**Figure 1B**). We detected most high-level taxonomic groups of eukaryotes and, on average, 55 ± 38% (sd) of the reference species clusters were detected at least once (**Figure 2A, Figure S2, Table S2, Table S3**(available on zenodo 10.5281/zenodo.24104693)). The highest total number of detections was observed for Streptophyta (plants), Ecdysozoa (includes crustaceans, insects, roundworms), Chordata, Spiralia (flatworms, annelids, mollusks), Dikarya (fungi), Stramenopiles, and Chlorophyta (green algae). Only three groups, Chordata, Dikarya, and Streptophyta, were seen in all major habitats (human: gut, mouth, skin, vagina; animal: gut; aquatic: ocean and marine, freshwater; air; soil; other environments including anthropogenic habitats such as buildings). This finding may be due to the widespread nature of fungal spores and plant pollen or could reflect the ease with which these materials can contaminate samples during laboratory processing. Various species such as the human skin fungus *Malassezia*, the fungal plant parasite *Alternaria*, pine, and willow, as well as various mushrooms were detected in samples across the majority of habitats, possibly due to contamination with spores or pollen grains. We also detected multiple model organisms such as mice, *Drosophila* and *Caenorhabditis elegans*; and organisms of high research interest such as *Plasmodium* in various environments. This type of contamination may occur due to consecutive sequencing of model organisms and metagenomic samples on the same machines as well as during sample preparation, e.g., sample collection, DNA extraction or library preparation. We manually curated an overview of putative contaminants (**Table S4).** We provide eukaryotic profiles across samples in the supplementary materials and the SPIRE website (https://spire.embl.de/eire/).

**Figure 2.**
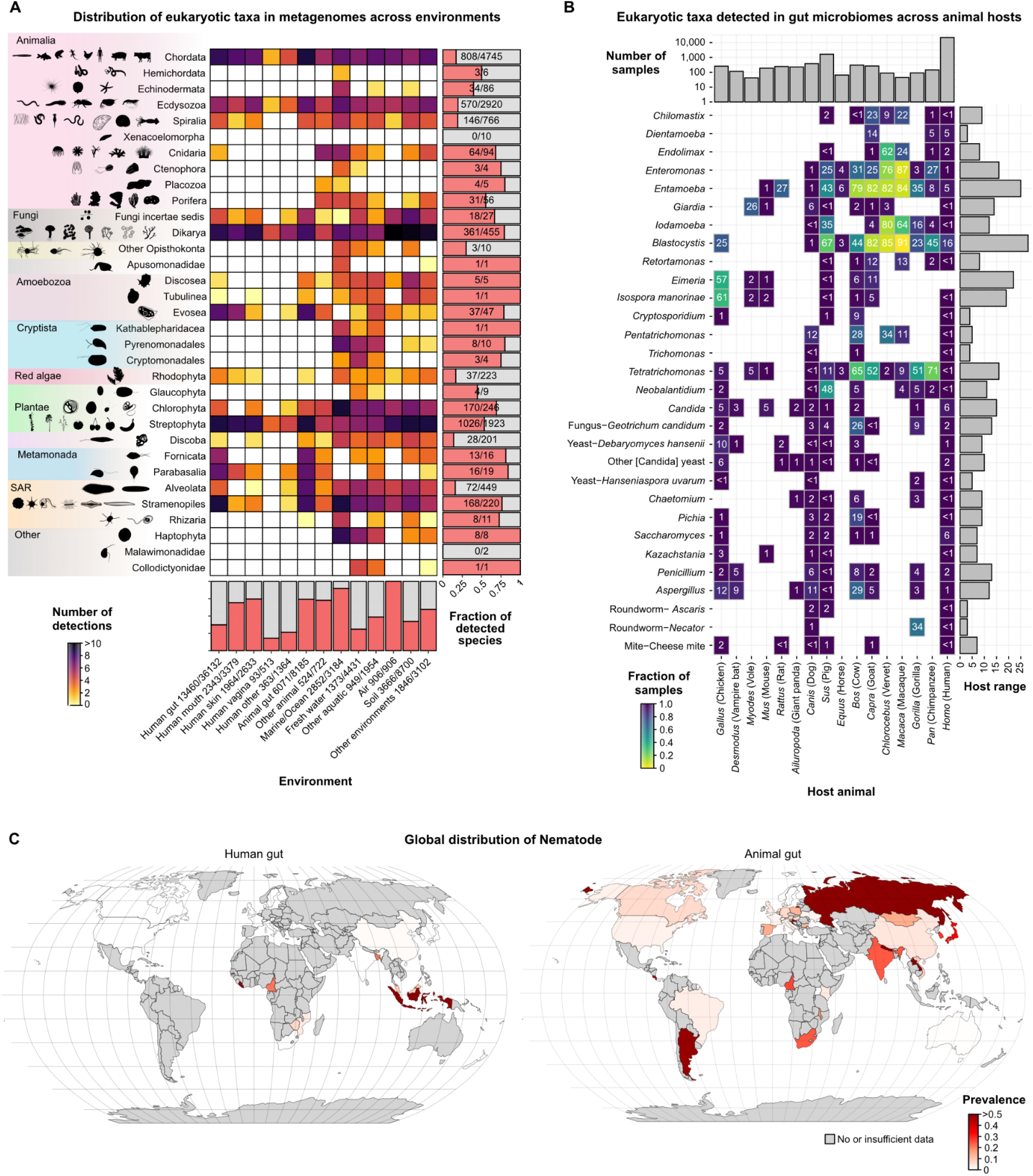
Eukaryotes are detectable across all habitats in WMS data. **(A)** Overview of detectable eukaryotes in WMS data across all major habitats (schematic figures of eukaryotes were downloaded from PhyloPic^183^). Heatmap color represents the total count of species/genus clusters across each taxonomic group (y-axis) and habitat group (x-axis). Multicellular eukaryotes capable of producing spores and pollen grains are prevalent in the majority of environments due to efficient dispersal which can also result in contamination of samples. Detections of human DNA are excluded from the figure for samples obtained from human subjects. **(B)** Overview of sharing of the symbiotic eukaryotes in the gut between human and the most sequenced animals (≥50 samples). Bars on the right side show the number of animal host species in which each symbiont was present (host range). Full overview available in **Figure S6**. Top bars show the total number of samples for each host species. **(C)** Global distribution of nematodes in the human and animal gut. In the human gut, nematodes show a distribution consistent with neglected tropical diseases, with the highest prevalence in tropical low- and middle-income countries. In contrast, in the animal gut, nematodes are broadly distributed worldwide without a clear dependence on country income status.

In ocean and freshwater samples, we observed the highest diversity of eukaryotes - all except two groups of eukaryotes (Xenacoelomorpha and Malawimonadidae) (**Figure 2A**, **Figure S2**) were detectable across aquatic habitats. We were able to confirm the previously observed cosmopolitan distribution of many algae (for more detail see https://spire.embl.de/eire/). For example, the most prevalent species in saline aquatic environments were microscopic algae, such as Haptophyta: *Emiliania huxleyi* (which is part of *Gephyrocapsa huxleyi* species^75^) and *Phaeocystis*; Chlorophyta (green algae): *Chloropicon*, *Chloroparvula*, *Bathycoccus*, *Pycnococcus*; Cryptophyta: *Teleaulax*; Stramenopiles: *Pseudo-nitzschia*, *Aureococcus anophagefferens*, and *Florenciella*. All of these algae have previously been suggested to play a large role in ecosystem functioning^76,77^. Multiple fungi were also observed in marine water, e.g., *Rhodotorula* and *Malassezia*. While some of them have been reported as pelagic species^78^, it is also possible that they are contaminants^79^ or passively dispersed as spores. We observed known ecological niche separation in the genus *Phaeocystis*^80^, which is responsible for foamy blooms in the open ocean and across the shoreline^81^. At the species level, we found that *Phaeocystis globosa* was more prevalent closer to equatorial regions and *Phaeocystis antarctica* in colder areas of the ocean (**Figure S3**). We also detected multiple groups of marine macro-organisms belonging to all major groups of Opisthokonta such as Placozoa (*Trichoplax*), mollusks (e.g., *Mytilus*, *Aplysia*, and *Sthenoteuthis*), fish (e.g., *Diaphus*, *Triphoturus*, *Lampadena*, *Auxis,* and *Cubiceps*), Cnidaria (*Nanozoanthus*, *Haloclava,* and *Aurelia*), Ctenophora (e.g., *Beroe*), Porifera (*Halisarca*, *Hippospongia*, *Geodia*, *Poecillastra*, and *Aplysina*). In freshwater habitats, we observed predominantly microscopic algae Chlorophyta (*Choricystis*, *Mychonastes, Micractinium, Ankyra, Chlamydomonas, Tetradesmus, Micromonas, Botryococcus, Scherffelia, Bathycoccus, Chlorella*); Cryptomonadales (*Cryptomonas curvata*); Pyrenomonadales (*Teleaulax amphioxeia*); Stramenopiles (*Cyclotella*, *Thalassiosira*, *Nannochloropsis*, *Asterionella*, *Ulnaria*); Alveolata - *Paramecium*; and multiple groups of amoebas (Discosea - *Acanthamoeba*; Tubulinea - *Vermamoeba*). Multicellular animals were represented in freshwater habitats: for example, we detected a crustacean water “flea” - *Daphnia* (Ecdysozoa). We also observed multiple plant species (Streptophyta) likely due to dispersal of pollen grains, seeds, and deposition of leaves or wood debris in the water. We also detected eukaryotes in extreme environments, for example, a green algae (*Dunaliella salina*) and brine shrimp (*Artemia*) in hypersaline lakes and salterns.

In soil the most common detectable taxonomic groups were fungi, plants and some metazoa. In particular, we detected plant parasitic fungi such as *Alternaria, Fusarium, Ilyonectria*; multiple mycorrhiza-forming symbiotic fungi - *Glomus*, *Russula*, *Cairneyella*, *Lactarius*; avirulent plant symbionts - *Trichoderma*; parasites of Nematoda - *Pochonia*, common saprotrophs for example *Podila* (also known as *Mortierella*), *Pseudogymnoascus* and *Penicillium*. Metazoa were represented by termites (e.g., *Heterotermes*, *Compositermes*), soil nematodes (*Xiphinema*), root-knot nematodes (*Meloidogyne*), oligochaete earthworms (*Aporrectodea*). Among plants, the most common detectable genera were *Salix/Populus* (e.g., willow and poplar trees), *Triticum* (wheat and its relatives), *Sphagnum* (moss), *Carex* (many species of monocot grasses/sedges), and *Solanum* (potatoes, tomatoes and others). It is established that many species of plants can form symbioses with multiple species of fungi^82^. We observed several positive associations (Pearson correlation) between plants and fungal species which are known to form symbiotic relationships (**Figure S4**). For example, mushroom-forming fungi from Basidiomycota: *Lactifluus*, *Russula*, *Lactarius*, *Paxillus* were co-occurring with various plant species such as *Camellia*, *Ilex*, *Olea*, *Fagus*, *Betula*, *Prunus* and many others^83^. Among arbuscular mycorrhizal fungi (Glomeromycotina), we also observed several positive associations with various monocot and dicot plants, for example, *Glomus* was associated with *Chionochloa*, *Milla*, and *Spondias*. Multiple fungal parasites were associated with agricultural species, for example, *Zea* (corn) was associated with *Ustilago*, *Fusarium*, and *Alternaria*. *Ustilago* was also associated with a cosmopolitan genus of monocot grasses - *Chrysopogon*, implying that it can serve as a vast reservoir of fungal opportunistic pathogens in natural habitats^84^.

In the air, the highest detection counts were seen for plant-parasitic fungi that produce high numbers of spores: *Alternaria*, *Pyrrhoderma*, *Ganoderma*, *Trametes*, *Fomitopsis*, and *Schizopora*, in line with previous studies^85^. The most frequently detected plant species were *Styphnolobium japonicum* (Fabaceae), *Nephrolepis biserrata* (fern), and multiple monocot grasses, which are known to strongly rely on wind pollination.

### Global distributions of eukaryotes across animal and human microbiomes

In the human gut, we detected 234 symbiotic and 438 food species of eukaryotes (**Figure 2A-C**, **Figure S5**, **Figure S6**). The most prevalent was the gut commensal *Blastocystis* sp. (observed in 16% of samples). Twelve eukaryotic symbiotic (including mutualistic, pathogenic, and parasitic) species clusters had prevalences >1% in the human gut: *Dientamoeba*, *Endolimax*, *Enteromonas*, *Entamoeba*, *Blastocystis*, *Candida*, *Geotrichum*, *Debaryomyces*, other “Candida” (*Nakaseomyces*), *Saccharomyces*, *Penicillium*, and *Aspergillus*. The human samples were also found to harbor a large number of species representing rare symbionts which are often pathogens (observed in <1% of the gut samples): *Giardia*, *Cryptosporidium*, *Neobalantidium*, *Trichomonas*, *Tetratrichomonas* and *Pentatrichomonas*. We observed a high prevalence of the genus *Entamoeba* (5%) in the human gut samples. However, only 166 samples (<0.5%) contained at least one virulence gene from *Entamoeba histolytica*, suggesting low overall prevalence of pathogenic *Entamoeba*. We also detected *Entamoeba gingivalis* in 18% of oral samples, which is a known commensal symbiont in the oral cavity in the human population and might be relevant for development of periodontal diseases^86^. Furthermore, in both the mouth and gut, we detected various symbionts, e. g. *Candida*, *Malassezia*, *Saccharomyces*, Torula yeast (*Cyberlindnera*), *Trichomonas* as well as the most common dietary genera such as *Bos* (cow), *Sus* (pig) and *Gallus* (chicken), wheat/barley, apples, cherries, and strawberries. In skin samples, we observed *Demodex* mites (*Demodex folliculorum* - 14%, *Demodex brevis* - 7%), *Malassezia* fungus (*Malassezia restricta* - 65%, *Malassezia globosa* - 58%, *Malassezia sympodialis* - 21%, and *Malassezia furfur* - 0.8%) and multiple food species (e. g., banana, chicken) as well as pets or farm animals (cats, dogs, and chicken). In vaginal samples, the most frequently detected eukaryotic species were *Candida albicans* (8.6%) (known commensal and opportunistic pathogen in the reproductive tract^87,88^) and *Camellia* (2.1%) (tea plant, potentially a component of hygienic or medicinal products^89,90^, or might be a contaminant). In some human gut and skin samples, we also observed putative traces of application of medicinal plants. For example, in an ankylosing spondylitis study^91^, in samples from a fraction of subjects (14/134 with any detectable eukaryotes) with diseases, we observed DNA from the genera *Humulus*/*Cannabis*. *Cannabis* can be part of the pain relief medication during ankylosing spondylitis^92,93^. An additional explanation can be the consumption of beer since the genus *Humulus* includes hops, which is an ingredient in beer or various other products^94^.

In the animal gut samples (e.g., pig, cow, and multiple others), we detected 31 eukaryotic gut symbionts that were also detected in human gut samples (**Figure 2B, Figure S6**) and many traces of diet-derived species (e.g., wheat, soybean, sorghum and other related species). While symbiont detection rates in the human population were often low for many symbionts (<1%), in animal samples, detection frequencies were high and in some cases reaching >80%, for example, *Entamoeba*, *Enteromonas*, and *Blastocystis* in macaques. A notably diverse group of parasites, *Eimeria*/*Cyclospora*/*Isospora* (includes >26 species), which is common in animals (especially in birds^95^ and goats, **Figure 2B**), was nearly absent in human samples (i.e., < 1%). Multiple potentially food-derived organisms such as *Hanseniaspora*, *Chaetomium*, *Pichia*, *Kazachstania*, *Saccharomyces* and cheese mites were prevalent at low rates in humans and multiple animal species. *Candida* and *Saccharomyces* yeasts were the only symbiotic organisms in the subset that showed a higher prevalence (6%) in humans compared to other animals. Species with highly specialized diets such as panda (bamboo plants) and vampire bats (blood) carried only fungal symbionts (e.g., *Candida*, other fermentation yeasts, *Chaetomium*, *Aspergillus* for pandas and *Candida*, *Debaryomyces*, *Penicillium*, *Aspergillus* for vampire bats) and none of the above-mentioned protists. Surprisingly, rodents (including wild populations and animals in captivity) also had a low prevalence of protist symbionts which are common in primates or bovidae and did not have any *Blastocystis.* This result may imply lineage-specific eukaryote-host symbiotic associations.

### Estimation of eukaryotic pathogen burden in the human population

metaEukaryome is suitable for estimating and monitoring pathogen burden in the human and animal populations as well as in environmental WMS sequencing data. This is important for understanding how pathogenic and commensal species contribute to microbiome structuring, to the health of human, animal and plant populations. The pipeline can detect >200 species clusters of parasitic and pathogenic organisms of humans, animals and plants. Across the analyzed metagenomic data from the human gut, we observed more than 200 fungi, protists and multicellular animals which can be opportunistic pathogens and parasites (**Figure S5, Figure S6**, **Table S5**, **Figure 2B-C**, **Figure 3A-C**). The most common opportunistic pathogen in the human population was *Candida*, detected in almost all countries. We detected members of the genus *Candida* in 6% of gut, 5.4% skin, 11.9% mouth, and 8.8% vaginal metagenomes. *Candida* consists of multiple species of known opportunistic pathogens, such as *Candida albicans*, *C. tropicalis*, *C. parapsilosis* and emerging ones such as *Candida sojae*^96^. The majority of pathogens and parasites were present in the human population at low prevalence, however, some countries had a higher prevalence of parasitic nematodes and flatworms (**Figure 2C, Figure S7A**). In addition to common pathogens and parasites we detected rare ones, for example, yeast-like green microalga *Prototheca*^97^ in Guinea-Bissau (15 detections) and in Brazil (one detection). Among other rare pathogens in gut WMS data we detected kinetoplastid pathogen *Leishmania*^98^ (two detections in Canada), *Kudoa septempunctata* a cnidarian parasite^99^ (one detection in South Korea), Alveolate fish parasite - *Uronema marinum* (two detections in China), and an apicomplexan parasite - *Cryptosporidium* (one detection each in Tanzania and the United Kingdom, and three detections in Bangladesh)(**Figure S7B**).

**Figure 3.**
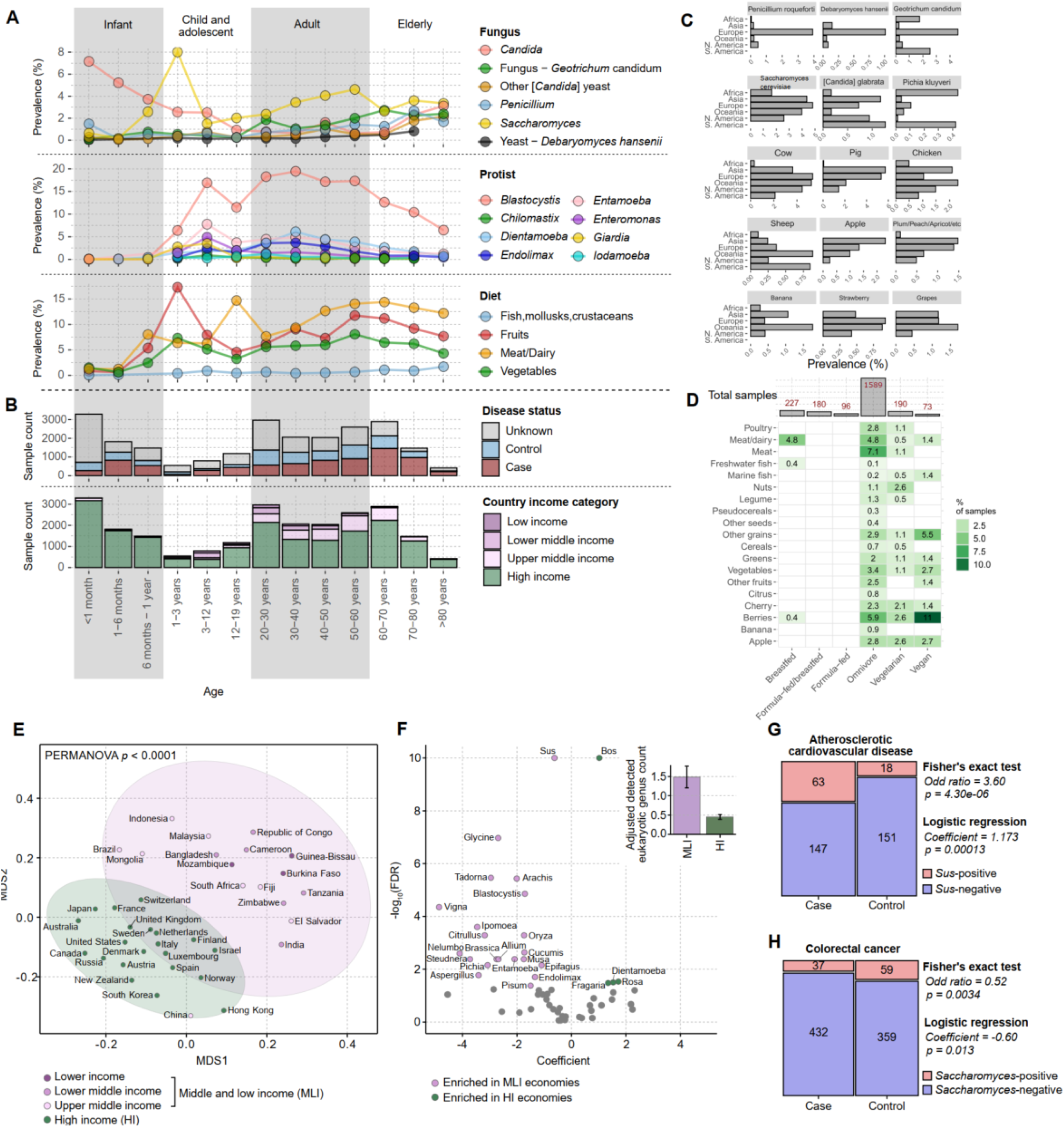
Eukaryote association with income level and diseases. **(A)** Distribution of most prevalent symbiotic and food eukaryotes across the age groups in the human gut. **(B)** Health and income status of subjects in the age-distribution in figure **(A)**. (**C)** Common diet-derived eukaryotes in human gut WMS samples from different continents. **(D)** Type of diets and detectable eukaryotes. **(E)** Multidimensional scaling (MDS) of country- or region-level average gut eukaryote profiles. Countries/regions are colored by the World Bank income category, with low-, lower-middle-, and upper-middle-income economies grouped as middle- and low-income (MLI), and compared with high-income (HI) economies. Ellipses indicate group-level clustering. Eukaryote composition was significantly associated with economic status (PERMANOVA, p < 0.0001). **(F)** Differential prevalence analysis of eukaryotic genera between MLI and HI economies. Each point represents a genus, plotted by effect size (coefficient) and significance, shown as −log10(FDR-adjusted p value). Purple indicates genera enriched in MLI economies, green indicates genera enriched in HI economies, and selected taxa are labeled. The inset shows the adjusted number of detected eukaryotic genera per sample in MLI and HI economies. **(G,H)** Associations between gut eukaryote detection and host disease status. **(G)** Detection of *Sus* in atherosclerotic cardiovascular disease and **(H)** detection of *Saccharomyces* in colorectal cancer are shown as the numbers of positive and negative samples in case and control groups. Fisher’s exact test and logistic regression statistics are indicated.

Many human and animal commensals, mutualists, parasites and pathogens were not detected in environmental samples at high rates except in sewage, wastewater and other anthropogenic environments, e.g., hospital buildings. For example, in sewage we detected *Blastocystis* (15% of samples), *Entamoeba* (12%), *Dientamoeba* (5%), *Enteromonas* (3%), *Endolimax* (1%), *Iodamoeba* (3%), *Neobalantidium* (16%), *Candida* (5%), *Ascaris* (7%) and multiple other potentially pathogenic and commensal species. We also detected multiple mutualistic and food-fermentation fungi (which sometimes also can be opportunistic pathogens), for example, *Geotrichum* (25%), *Debaryomyces* (6%), and *Saccharomyces cerevisiae* (36%) in those samples. Thus, monitoring organisms in sewage/wastewater can be instrumental not only to estimate eukaryotic pathogen burden in the human population^100–108^ but also to evaluate frequencies of mutualistic, food-derived, commensal eukaryotic organisms, which in turn expands our understanding of symbiont circulation across ecosystems and impact on populations from One-Health perspective^109,110^.

We analyzed distribution of soil-transmitted helminth (STH) (predominantly nematodes) across human intestines, animal intestines, and waste handling facilities motivated by their high contribution to pathogen burden and the superior performance of metaEukaryome in parasitic worm detection (**Figure S1A–S1C**). We found that carriage of nematodes in humans, classified as Neglected Tropical Diseases (NTDs), predominantly occurred in countries such as Malaysia, Indonesia, Bangladesh, Mozambique, Zimbabwe, Cameroon, Guinea-Bissau, and Liberia, but was rarely observed in countries outside of the tropical regions which are classified by the World Bank^111,112^ as high income (**Figure 2C, Figure S7A**). In contrast, parasite prevalences within the animal intestine and wastewater-handling microbiomes remained high in all countries (**Figure 2C**). This suggests that helminth infection in animal hosts remains widespread across countries, highlighting its continued relevance for veterinary health and parasite surveillance^113,114^. However, these country-level prevalence patterns should be interpreted with caution because of substantial sampling bias across countries. For example, the particularly high detection rates in Malaysia, Indonesia, and Liberia likely reflect the fact that many public metagenomic samples from these regions were generated in studies specifically focused on helminth infection^66,68^. Likewise, animal gut samples may also differ systematically in host species across countries.

### Human gut eukaryote profiles vary with country income status, diet, age and disease

While for prokaryotes it is well established that there is an age-related development/succession of the microbiome, for eukaryotes such knowledge is limited^115–121^. We observed changes in numbers and types of detectable eukaryotes in the gut throughout lifetime. In particular, as expected, newborns and infants before the age of 6 months exhibited limited prevalence of the most common eukaryotic symbionts and diet species (**Figure 3A**). *Candida* prevalence in the gut was highest during the first year of life (6.6%), potentially reflecting colonization from the vaginal, skin, or other microbiomes^122^, and declined with age thereafter. With the introduction of solid food in infants after 6 months, multiple eukaryotic species appeared in the samples. Rates of detectable eukaryotes stayed stable across age groups, with Blastocystis tending to increase in early life and decline later, and Saccharomyces and meat species tending to increase with age. Similar age-associated patterns for *Candida*, *Saccharomyces*, and *Blastocystis* have been reported previously^38^. However, it is important to note that the underlying data in each age group are affected by other confounding factors such as health status, geographic location, income status and sampling size, which are likely to contribute to any age-related signal (**Figure 3B**). While we observed interesting trends in age-related dynamics of prevalence of eukaryotes, it is important to mention that at the moment there is an uneven sampling of different age groups in the human population. In particular, there are more samples from the first year of life than from the next two decades of life. Increasing sampling size of undersampled age groups, geographic regions with in depth metadata on dietary preferences will enable interpretation of eukaryome development in early life and its effect on future health outcomes and ageing. Mass-produced meat and fruit species were observed on all continents, although detectability varied (**Figure 3C)**. We observed a tendency towards a higher diversity of dietary species in Africa and Asia (**Figure S8**, **Table S6**) compared to Europe and Americas. China had the highest diversity of consumed seafood, amphibian, reptilian and mammalian species(**Table S6**, **Figure S8, Figure S9A**). We detected evidence of honey consumption in multiple countries, including the Hadza population from Tanzania, as previously shown (**Figure S9B**)^123^. Yet, many species likely remained undetected because of limited numbers of reference genomes. Eukaryotic detectability depended on reported diet type, for example, detections of meat, poultry and meat/dairy species were 6.4, 9, and 2.6 times higher in omnivore than in vegetarian subjects (**Figure 3D**).

To examine in more detail how human gut eukaryote profiles vary by geographic location or lifestyle, we projected country- or region-level average eukaryote profiles into a multidimensional scaling (MDS) space (**Methods**). We observed that eukaryote composition was more clearly separated by economic status than by geography or ethnicity, with the most pronounced distinction between High-income (HI) and Middle- and low-income (MLI) economies (**Figure 3E**, PERMANOVA p < 0.0001). This pattern is consistent with prior prokaryote-focused gut microbiome studies in which microbiome composition also differed by host country income status^124^. To identify specific eukaryotes driving these differences, we compared taxon prevalence between HI and MLI groups using mixed-effects logistic regression, adjusting for sex and sequencing depth and accounting for study as a random effect (**Methods**). Overall, a greater number of eukaryotes were detected in MLI samples (adjusted mean detected genera: 1.48 in MLI vs. 0.45 in HI), and more taxa were enriched in MLI economies (**Figure 3F**). Notably, diet-related eukaryotes showed strong differences. Legumes including *Glycine* (soybean), *Arachis* (peanut), and *Vigna* (some of the legumes, for example, beans) were highly enriched in MLI economies (mixed-effects logistic regression, FDR-corrected p-value (q) < 1×10⁻⁴), and with the exception of *Fragaria* (strawberry) and *Rosa* (rose and many other related species), most plant-based foods were also enriched in MLI economies (q < 0.05). Among animal-derived foods, *Sus* (pork) and *Tadorna/Anas* (duck) were enriched in MLI economies, whereas *Bos* (beef) was significantly enriched in HI economies (q < 1×10⁻⁵). These findings suggest that dietary differences between income categories underlie much of the variation in eukaryote profiles. Beyond diet-derived signals, symbiotic protists including *Blastocystis*, *Entamoeba*, and *Endolimax* were enriched in MLI economies (q < 1×10⁻⁵), whereas *Dientamoeba* was more frequently detected in HI economies (**Figure 3F**). This is consistent with previous studies reporting that dientamoebiasis, unlike other parasitic infections, is also common in high-income countries^125,126^.

We further investigated associations between eukaryotes and non-transmissible host diseases using diseases with sufficient case–control sample sizes, adjusting for sex, sequencing depth, and study (**Methods**). The strongest signal was observed in atherosclerotic cardiovascular disease, where *Sus* (pork) was more frequently detected in the patient group (**Figure 3G**, logistic regression *p* = 0.00013). This may relate to previous reports implicating red or processed meat consumption as a cardiovascular disease risk factor^127^. However, as *Sus* detection does not exclusively reflect these dietary sources and because conflicting findings have also been reported^128^, the results should be interpreted cautiously. In colorectal cancer (CRC) cohorts, *Saccharomyces* was more frequently observed in controls than in CRC patients (**Figure 3H**, logistic regression *p* = 0.013). This is consistent with prior studies showing that *Saccharomyces cerevisiae* and *Saccharomyces boulardii* can promote apoptosis of cancer cells and attenuate colon cancer progression^129–132^.

### Gut protists and fungi show contrasting associations with prokaryotic community structure

The human gut microbiome harbors a large diversity of prokaryotes that may engage in interactions with eukaryotic microorganisms. To examine these interactions more comprehensively, we correlated the metaEukaryome-derived eukaryotic profiles with mOTUs-based prokaryotic profiles from SPIRE and Metalog. We first tested whether the presence of individual eukaryotic genera was associated with prokaryotic alpha diversity, controlling for potential confounders including host sex and sequencing depth, with study-country modeled as a random effect (**Methods**). *Blastocystis* and *Dientamoeba* were associated with increased prokaryotic alpha diversity, whereas *Candida* and *Nakaseomyces* were associated with reduced alpha diversity. Detection of several food-derived taxa, including *Linum* (flax), *Fragaria* (strawberry), *Vitis* (grape), *Musa* (banana), and *Bos* (beef), was also associated with higher prokaryotic alpha diversity (linear mixed-effects models, q < 0.05, **Figure 4A**). We then examined associations between eukaryotes and the relative abundance of individual prokaryotic taxa using zero-inflated beta regression^133^, adjusting for sex, sequencing depth, and study–country (**Methods**). Overall, symbiotic eukaryotes, particularly some fungi and protists, showed substantially stronger associations with both alpha diversity and prokaryotic relative abundance than did food-derived taxa. In particular, *Blastocystis*, *Dientamoeba*, *Candida*, and *Nakaseomyces* exhibited the largest numbers of significant associations (zero-inflated beta regression, q < 0.05), indicating that these eukaryotes are linked to broad shifts in gut prokaryotic community structure (**Figure 4A**).

**Figure 4.**
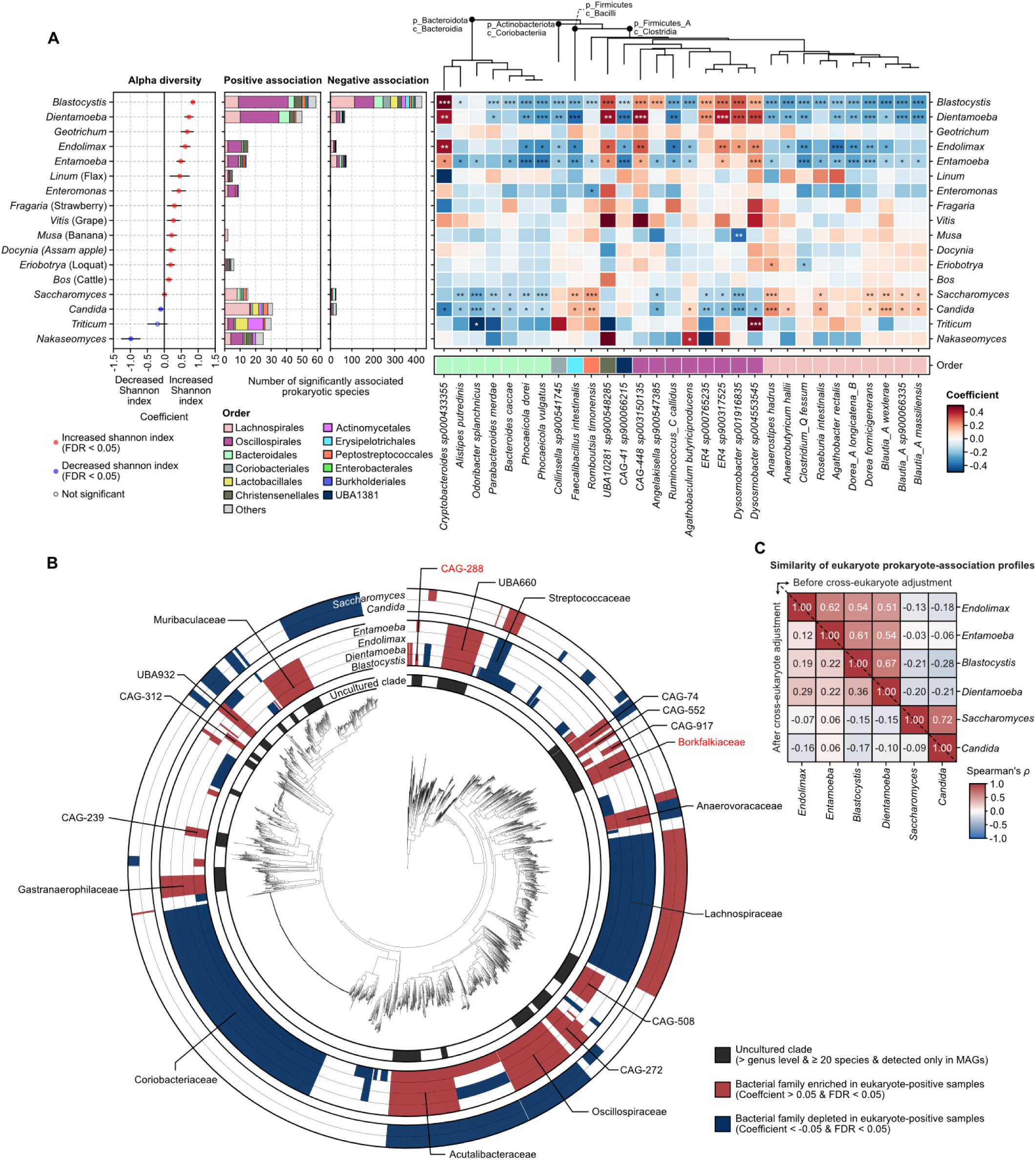
Association between eukaryotes and prokaryotes in the human gut. **(A)** Associations between gut eukaryotes and prokaryotic community features across human gut metagenomes. Left, effect sizes for associations between eukaryote detection and prokaryotic alpha diversity (Shannon index), estimated using linear mixed-effects models adjusted for sex and sequencing depth, with study-country included as a random intercept. Red- and blue-filled points indicate significant positive and negative associations, respectively (q < 0.05). Middle, numbers of significantly associated prokaryotic species showing positive or negative associations with each eukaryote, stratified by prokaryotic order. Right, regression coefficients from zero-inflated beta regression models for representative eukaryote-prokaryote species associations, with prokaryotic species ordered by phylogeny. Red cells indicate prokaryotic species enriched in samples in which the corresponding eukaryote was detected, whereas blue cells indicate depletion. (*q<0.1, **q<0.01, ***q<0.001) **(B)** Distribution of eukaryote-associated prokaryotic species across the human gut bacterial phylogeny provided by HRGM v2. Concentric rings indicate significant positive (red) or negative (blue) associations for *Blastocystis*, *Dientamoeba*, *Endolimax*, *Entamoeba*, *Candida*, and *Saccharomyces*. Black segments denote uncultured clades, defined as family-level or higher clades containing at least 20 species and detected exclusively in metagenome-assembled genomes (MAGs). The clades highlighted in red, CAG-288 and Borkfalkiaceae, correspond to the lineages containing species detected in the axenic *Blastocystis* culture. CAG-288 contains CAG-345 sp000433315, whereas Borkfalkiaceae contains *Ca.* Borkfalkia ceftriaxoniphila. **(C)** Similarity of eukaryote-prokaryote association profiles among major gut protists and fungi, measured as Spearman’s correlation of regression coefficients across prokaryotic species. The upper triangle shows correlations before cross-eukaryote adjustment, and lower triangle shows correlations after adjustment for co-detected eukaryotes.

We found that four protist genera, *Blastocystis*, *Dientamoeba*, *Endolimax*, and *Entamoeba*, showed highly similar association patterns across prokaryotic species. Samples carrying these protists were enriched for Oscillospirales and depleted for Lachnospirales. In contrast, samples carrying the fungi *Candida* and *Saccharomyces* tended to harbor more Lachnospirales and fewer Oscillospirales (**Figure 4A**). A similar contrast between protists and fungi was also evident when the results were projected onto the full gut bacterial phylogeny^134^ (**Figure 4B**). Within the protist and fungal groups, eukaryotic genera showed similar patterns of positive and negative associations across prokaryotic species, with pairwise Spearman’s ρ values generally exceeding 0.5, whereas protist-fungal comparisons showed weak negative correlations (**Figure 4C**). Because these similarities could simply reflect co-occurrence of multiple eukaryotes in the same samples, we repeated the analysis with cross-eukaryote adjustment (**Methods**). Under this model, the correlation between *Saccharomyces* and *Candida* disappeared, suggesting that their similar association profiles were primarily driven by co-detection, whereas positive correlations among protists were retained to a substantial extent (**Figure 4C**).

### *Blastocystis* is linked to metabolically reduced uncultured bacteria

Several prokaryotes enriched in samples positive for four protist genera, *Endolimax*, *Entamoeba*, *Blastocystis*, and *Dientamoeba*, were assigned GTDB r207 placeholder names such as UBA660, CAG-552, and CAG-508, indicating that they remain uncultured and poorly characterized. We therefore tested whether protist-associated bacteria have typical genomic features of symbiotic prokaryotes and preferentially belonged to exclusively uncultured gut bacterial clades, defined here as clades above the genus level comprising at least 20 species that have been recovered exclusively as metagenome-assembled genomes (MAGs). This analysis revealed that multiple bacteria enriched in protist-positive samples indeed showed strong overlap with these exclusively uncultured clades (**Figure 4B**).

Among protists, we specifically focused on *Blastocystis*, the most frequently detected protist in the gut and the one showing the strongest associations with prokaryotic taxa (**Figure 4A**). Prokaryotes enriched in *Blastocystis*-positive samples exhibited reduced metabolic independence^134,135^ (Pearson’s r = -0.277, p < 1×10⁻^13^, **Figure 5A**), as reflected by extensive loss of core metabolic KEGG modules^136^. Consistent with this, trait annotations from metaTraits^137^ showed that these species tended to have reduced genome sizes (Pearson’s r = -0.207, p < 1×10⁻^7^), higher pangenome openness (Pearson’s r = 0.188, p < 1×10⁻^6^, **Figure 5B**) and limited capacity to grow on disaccharides such as trehalose and sucrose (Logistic regression p < 1×10⁻^6^, **Figure 5C**). These features may explain why these taxa have remained difficult to culture. We further compared the genome contents of prokaryotes enriched versus depleted in *Blastocystis*-positive samples at the level of KEGG modules. Prokaryotes enriched in *Blastocystis*-positive samples tended to lack a broad range of modules involved in metabolism of cofactors and vitamins, lipid metabolism, carbohydrate metabolism, nucleotide metabolism, and amino acid metabolism (Fisher’s exact test q < 0.01, **Figure 5D**), suggesting dependence on other organisms for core metabolic functions.

**Figure 5.**
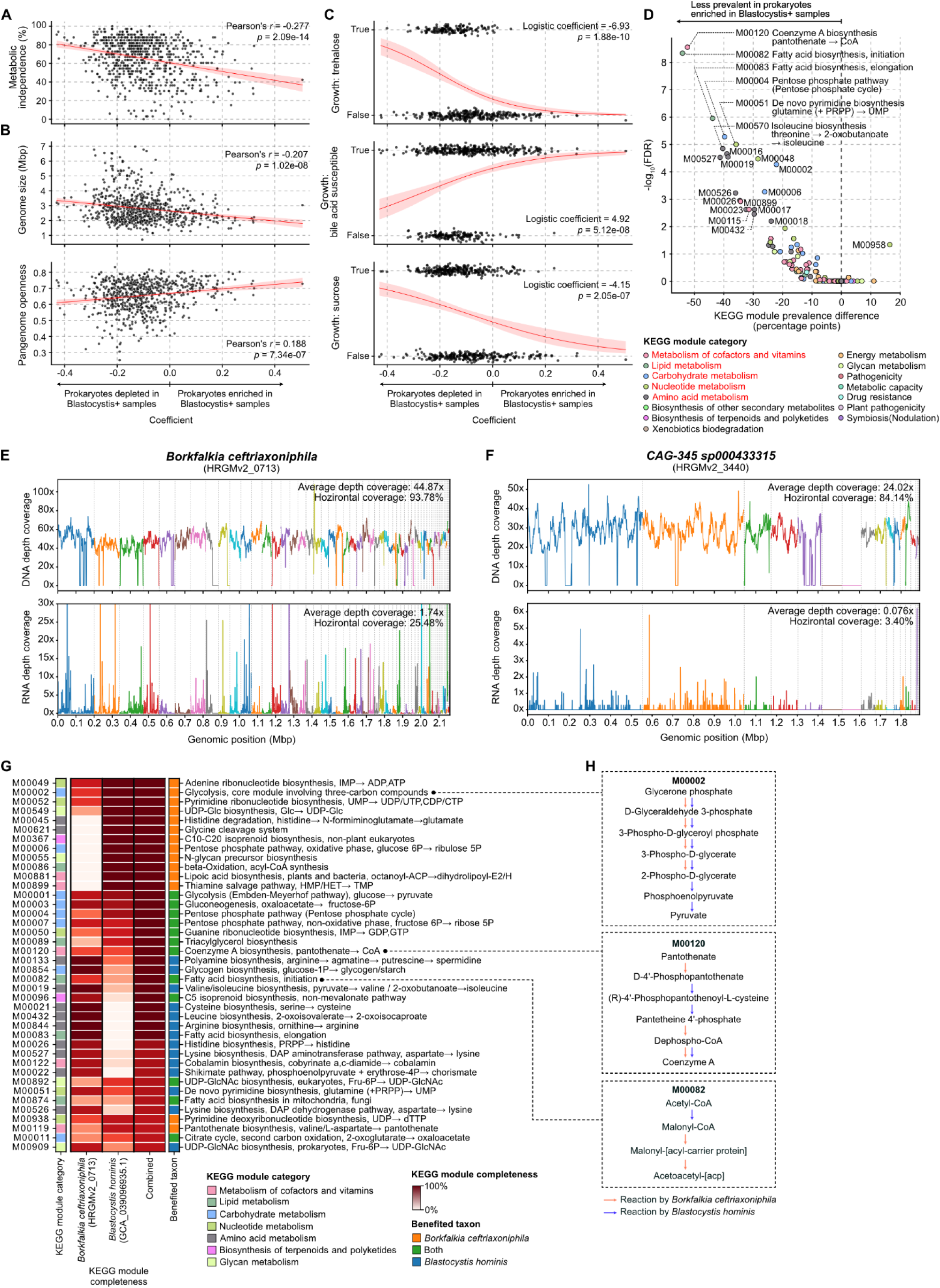
Putative symbiotic relationship between Blastocystis and uncultured bacteria. (**A–C**) Genomic and phenotypic traits of prokaryotes whose abundances are associated with the presence of *Blastocystis* in the human gut. Positive x-axis values indicate prokaryotic species enriched in *Blastocystis*-positive samples. (A) Metabolic independence, estimated from the number of core KEGG metabolic modules. (B) Genome size and pangenome openness. Red lines and shaded areas indicate linear regression fits and 95% confidence intervals (CIs). (C) Three growth-related traits from metaTraits showing strong associations with *Blastocystis* enrichment. Traits are shown as binary values (true or false), with jitter applied for visualization. Red lines and shaded areas indicate logistic regression fits and 95% CIs. (**D**) Comparison of KEGG modules encoded in the genomes of prokaryotes enriched in *Blastocystis*-positive samples (coefficient > 0.1, FDR < 0.05) and depleted in *Blastocystis*-positive samples (coefficient < -0.1, FDR < 0.05). Positive x-axis values indicate KEGG modules that are more prevalent in prokaryotes enriched in *Blastocystis*-positive samples. The five KEGG module categories highlighted in red contain modules showing significant differences at FDR < 0.01. (**E–F**) Prokaryotes detected in long-read DNA-seq and short-read RNA-seq data from an axenic culture of *Blastocystis hominis* (Nand II), together with genome coverage profiles. (**E**) *Ca.* Borkfalkia ceftriaxoniphila and (**F**) CAG-345 sp000433315 were detected in both DNA-seq and RNA-seq data, with coverage distributed across multiple contigs spanning their genomes. Vertical dashed lines indicate contig boundaries, and different colors denote distinct contigs in the corresponding MAGs. (**G**) Complementarity of KEGG modules encoded by the genomes of *Blastocystis hominis* (Nand II) and *Ca.* Borkfalkia ceftriaxoniphila. The combined column indicates KEGG module completeness after pooling enzymes encoded by both genomes. Shown are the 40 complementary KEGG modules with the highest combined completeness, in which modules incomplete in one genome can be complemented by enzymes encoded in the other. (**H**) Detailed reaction-level complementarity for selected KEGG modules among the complementary modules shown in (G). M00002, M00120, and M00082 were among the modules commonly missing in prokaryotes enriched in *Blastocystis*-positive samples in (D), and this panel shows how specific reactions within these modules can be completed by enzymes encoded in the partner genome.

We next asked whether some of these bacteria might live in symbiosis with *Blastocystis*. To identify specific candidates, we analyzed long-read DNA sequencing and RNA sequencing data generated from *Blastocystis* cultures^138^ and searched for co-detected prokaryotes (**Methods**). Although xenic cultures contained diverse gut commensal prokaryotes (**Figure S10A, Table S7**), this was difficult to interpret as evidence of symbiosis given the nature of xenic culture. Strikingly, however, DNA sequencing of a *Blastocystis hominis* culture previously reported as axenic (subtype 1, Nand II) detected *Candidatus* Borkfalkia ceftriaxoniphila^139,140^, CAG-345 sp000433315, *Staphylococcus epidermidis*, and *Staphylococcus warneri* (**Figure S10A**). Because *S. epidermidis* and *S. warneri* are common human skin commensals and were not detected in the RNA-seq data, they most likely reflect DNA sequencing-specific contamination. In contrast, *Ca.* B. ceftriaxoniphila and CAG-345 sp000433315, both of which remain uncultured, were also detected in the RNA-seq data, with transcription broadly distributed across their genomes (**Figure 5E–F**). Because both taxa belong to deeply branching lineages with no cultured representatives at the family level, their presence is less readily explained as conventional culturable carryover, although persistence through axenization cannot be excluded. Both taxa belong to clades that were enriched in *Blastocystis*-positive samples in the global metagenomic analysis (**Figure 4B**). To our knowledge, this represents the first evidence consistent with the presence of active *Ca. B.* ceftriaxoniphila and CAG-345 sp000433315 in a *Blastocystis* culture.

Finally, motivated by the extensive metabolic deficiencies observed in prokaryotes enriched in *Blastocystis*-positive samples, we examined whether enzymes encoded by the genomes of *Blastocystis* and *Ca.* B. ceftriaxoniphila could complement one another metabolically (**Methods**). We found that many KEGG modules were complemented across the two genomes, in some cases by enzymes encoded by *Ca.* B. ceftriaxoniphila, in others by *Blastocystis*, and in some cases by both partners contributing distinct missing steps to complete the same module (**Figure 5G**). Notably, M00120 (coenzyme A biosynthesis) and M00082 (fatty acid biosynthesis, initiation), which were among the KEGG modules most frequently missing in prokaryotes enriched in *Blastocystis*-positive samples (**Figure 5D**), were completed jointly by *Blastocystis* and *Ca.* B. ceftriaxoniphila (**Figure 5H**). Similar complementary modules, again including M00120 and M00082, were also observed for CAG-345 sp000433315 (**Figure S10B**). Together, these results support a candidate symbiotic relationship between *Blastocystis* and these uncultured bacteria, potentially involving metabolic complementarity.

## Discussion

Eukaryotes play a key role in ecosystem functioning but are often low-abundant in WMS samples. In this study, we took advantage of accumulated large-scale WMS data^60^ and curated metadata^61^ to investigate associations between eukaryotes and prokaryotes at comprehensive phylogenetic and ecological scales. We developed a specialized pipeline that addresses key technical issues by taking advantage of biological properties of organisms such as high ploidy of mitochondrial and plastid genomes, enabling high qualitative sensitivity across many lineages of eukaryotic organisms across Earth’s habitats.

Across environmental WMS samples we observed many common and rare eukaryotic species. In particular in marine samples we observed many major cosmopolitan species of algae, and protists as well as various multicellular organisms, consistent with previous studies using metatranscriptomic data from the ocean^141^. As expected, in soil samples we detected predominantly fungal and plant species because of high relative biomass from these taxonomic groups and available reference genomes. Furthermore, we were able to confirm several known associations between plants and fungal mutualists and pathogens (**Figure S4**). These results together suggest that eukaryotes can be systematically characterized using metaEukaryome across any types of WMS data and also can be used for discovery of novel eukaryote-environmental properties, eukaryote-eukaryote and eukaryote-prokaryote and other combinations of associations.

We found that the presence of eukaryotes was strongly associated with the prokaryotic part of the community in the human gut (**Figure 4A,B**). Similar to the loss of bacterial diversity in high-income countries^142,143^, we observed a decline in the diversity of detectable eukaryotes. Loss of eukaryotes such as *Entamoeba histolytica* or helminths can be viewed as a positive development for health of the human population since these species can be parasitic and pathogenic, however, loss of other commensals (e.g., *Blastocystis*) may be a negative outcome for health and requires further investigation, considering that eukaryotes play a role in the development and modulation of the immune response and have the potential to protect from autoimmune diseases^144,145^.

We observed a higher number of positive associations of different dietary species with MLI countries (**Figure 3F**), suggesting higher diversity of foods. In HI countries, by contrast, diversity is likely to be lower due to consumption of highly-processed foods with less traceable DNA and fewer different food species in general. In the GI tract, food is digested, so detection of DNA is expected to be challenging. Digestion of DNA is dependent on the efficiency of the enzymes of the host and symbiotic community. High rates of detection of the DNA might be a result of chronic or situational disruption in digestion. For example, digestion efficiency declines with age, or during GI tract diseases, which could lead to more detections of diet-derived eukaryotes^146^. Low rates of detections can be a result of low food intake before sampling and/or high digestive capacity of the host enzymes and the microbiome and possibly the property of a healthy state of the organism. Additionally, the type of the product and cooking methods can have a high impact on detection efficiency due to variable quantities of high-quality DNA yield. For example, in dairy products the amount of DNA is lower compared to meat products from the same species^147,148^. Nevertheless, it is possible to expand knowledge of subjects’ dietary intake even if diet metadata was not reported. A recent study developed a specialized pipeline (MEDI) to evaluate the metabolic contribution of ingested food based on detections of diet species in the gut samples^48^. In the next steps, the combination of digestion rates information and metabolic contribution of food, combined with symbiont profiles will be insightful for: (1) evaluating the health of the GI tract and broad health outcomes of patients and healthy subjects; (2) developing personalized evidence-based dietary interventions to maintain health. It is important to note that such pipelines (e.g., MEDI and metaEukaryome) provide an insight into the dietary context of subjects. However these tools do not fully replace, but rather complement a dietary questionnaire because of variable digestion rates in populations and many foods or supplements do not leave any DNA trace.

In contrast to dietary species, gut symbionts are expected to be active within their microbiome and thus to be detectable even at relatively low sequencing depth. Amplicon/metabarcoding studies have detected many groups of symbiotic protists, fungi and multicellular organisms in humans^14^ and animals^149^. Eukaryotes have repeatedly been associated with various health phenotypes of the host^22,150^, microbiome structure^151,152^ and individual prokaryotic species^153^. Yet, many questions remain unanswered, for example, on the nature of interactions between protists, fungi, prokaryotes and the host. Such gaps in knowledge can be addressed using WMS data, considering that it allows species-level resolution for prokaryotes as well as being free from amplification-biases and includes sequence data for both eukaryotes and prokaryotes simultaneously. Free-living protists are known to harbor many uncultured prokaryotic (endo-)symbionts^154,155^. Likewise, we found that many uncultured bacterial clades were consistently associated with several protists in the human gut, and because these bacteria were distributed broadly across the bacterial phylogeny rather than confined to specific lineages (**Figure 4B**), this pattern suggests a general ecological link between protists and uncultured bacteria. The observation that these protist-associated bacteria tended to have reduced genome sizes and lack multiple core metabolic pathways is consistent with metabolic dependence on partner organisms and therefore with a symbiotic lifestyle^156,157^. This interpretation is further supported by culture-based evidence that DNA and RNA from *Ca.* Borkfalkia (Bacillota_A) and *CAG-345* (Bacillota_I), which were enriched in *Blastocystis*-positive samples, were detected in an independently validated axenic *Blastocystis* culture. Notably, earlier electron microscopy studies had suggested intracellular bacteria in *Blastocystis*^158–160^, and a recent review has revisited the possibility of bacterial endosymbiosis within *Blastocystis*^161^. In this context, our detection of *Ca.* Borkfalkia and *CAG-345* in axenic *Blastocystis* renews this hypothesis and motivates direct experimental validation, for example, by fluorescence *in situ* hybridization (FISH). Such symbiotic interaction can have relevance for health since it has been reported before that protists can drive evolvability of antibiotic resistance among prokaryotes in general^162^. More broadly, if a substantial fraction of gut uncultured bacteria depend on eukaryotic symbionts for metabolic complementation, this could help explain why many of these taxa have remained refractory to isolation and would provide a new framework for understanding uncultured members of the gut microbiome. Furthermore, this study suggests a revised view of community structuring in the gut. We hypothesize that in addition to the core gut microbial community^163^ and food-derived transient prokaryotes^164^, there is a community of reduced-genome prokaryotic symbionts of gut protists which are likely to perform different ecological functions compared to other prokaryotic species from the core community in the gut.

### Limitations and caveats of the study

#### Contamination

Multiple groups of organisms have highly dispersible life-stages, such as spores in fungi, pollen grains in plants, cysts and eggs in many groups of protists and animals. For example, *Malassezia*, a common human skin symbiont, is frequently considered a contaminant^79^ because shed epidermal material can enter samples. Human DNA is likewise commonly detected in environmental samples. This suggests that for accuracy and fine-grade tracing of rare organisms and pathogens, sampling and laboratory processing has to adhere to increased standards. *Low coverage and underestimation of prevalences and abundances due to prokaryotic targeting:* DNA extraction protocols can preferentially target the prokaryotic part of the community, which hinders efficiency and consistency of the estimates of the eukaryotic part of the community. *Coverage of reference genome database:* Although we recovered many eukaryotic species (the largest number of species among current tools), the current reference genome database accounts only for ∼1% of total eukaryotic species diversity when compared to the number of species in the Barcode of Life (accessed in August 2026)^165^. Consequently, more high-quality genomes from other species would substantially improve the results of the metaEukaryome and similar pipelines. This is an especially large limitation for geographic regions with many understudied species. *Limitations for interpretation of the relative abundance of eukaryotes:* Limitations in estimating relative abundance have to be considered when interpreting relative abundance of prokaryotes and eukaryotes together. Prokaryotic and eukaryotic cells can have varying ploidy^166,167^, which limits the estimates of absolute abundance or a conceptually comparable relative abundance of prokaryotic and eukaryotic organisms within any given sample. Additionally, in many habitats eukaryotes have low abundances, which is not always reflective of their high significance in the ecosystem. Finally, the metaEukaryome pipeline relies on nuclear 18S rRNA genes for detection of eukaryotes that have lost their mitochondrial genome or lack a sequenced one. This may impose constraints on abundance comparability between species detected with mitochondrial/plastid genomes and 18S rRNA genes. *Causal inference:* DNA-based detection alone does not establish whether detected eukaryotes are viable or metabolically active, nor does co-occurrence demonstrate direct interaction between eukaryotic and prokaryotic taxa. Directionality and cause-effect relationships have to be interpreted with caution. It is possible for prokaryotes to affect eukaryotes and eukaryotes to affect prokaryotes during community assembly or symbiotic co-evolution. Causality and directionality in these relationships need to be validated experimentally.

### Future directions

This work is an example of a comprehensive approach in studying ecosystems from a One Health perspective. The pipeline enables detection of hundreds of commensals, parasites and opportunistic pathogen species, as well as species with unknown ecological roles within a metaorganism or holobiont. It allows detecting putative novel interactions and complexities within the microbiome and with the host (e.g., putative symbiosis between *Blastocystis* or other amoebas with small-genome prokaryotes such as *Ca.* B. ceftriaxoniphila), which can lead to discovery and re-evaluation of the role of commensals/parasites in health and diseases in human, animals and the environment. As a result, understanding of cross-domain interactions can improve the capacity to isolate and cultivate the vast uncharacterized diversity of prokaryotes^168^. Developments in the detection of eukaryotes in WMS data tremendously expand capacity for understanding community structure even at strain level without amplification biases which are introduced in amplicon metagenomics/metabarcoding. This in turn enables detection of diet, parasites and leads to a holistic view of microbiomes. Future studies will focus on experimental validations and integration of WMS data, metabarcoding, eukaryotic meta-pangenomics and other omics approaches to investigate in depth the genetic diversity of eukaryotes and its impact on various phenotypes and structures of the ecosystems.

## Methods

### Description of metaEukaryome pipeline

The metaEukaryome pipeline contains a database of mitochondrial and plastid genomes, 18S rRNA genes and known virulence genes from eukaryotes, prokaryotes and viruses. Metagenomic reads were mapped against each of the reference databases using bwa v0.7^169^. NGless v0.11 and samtools v1.16 were used to remove reads that have <70 bp match to the reference and average nucleotide identity <96%, <98%, <97% and <95% for mitochondrial, plastid, PR2 (Protist Ribosomal Reference database) and virulence genes respectively (**Table S8**). Mapping quality criteria were set to mapq=1 for mitochondrial and plastid genome references and mapq=20 for PR2 and virulence genes. Average nucleotide identity criteria approximately correspond to operational approximation of species (for mitochondria) or genus (for plastids) boundary thresholds (**Table S8, Figure S11**). Unless otherwise specified, we used ≥5% of horizontal coverage (corresponding to at least ∼1Kbp for mitochondria and ∼5Kbp for plastids) as a threshold to consider species present for plastids and mitochondria reference genomes and ≥20% for PR2 and virulence genes. metaEukaryome can profile metagenomes of 10 Gb size in 3.9 hours on 24 cores. Performance can also be boosted by focusing only on the reference databases of interest.

### Genomes download and database building for metaEukaryome

Mitochondrial and plastid genomes were downloaded from GenBank from NCBI on 25.01.2021 which included >18,000 genomes, corresponding to 16,737 species/strains with names on NCBI Taxonomy. We ran fastANI all-against-all approach (with command fastANI --refList <list of genomes> --ql <list of genomes> -- matrix --minFraction 0.1 --fragLen 1000 -t 50 -o <output file name>). **Figure S11A-S11D** illustrates density of average nucleotide identity (ANI) and genome overlap estimates between each pair of genomes. ANI parameters were used to determine the optimal threshold for dereplication of plastid and mitochondrial genomes. As a result we obtained 11,932 species clusters across all available plastid and mitochondria genomes.

To extend the potential of the pipeline to detect eukaryotes with unavailable or without mitochondrial genomes (due to biological reasons such as organism being anaerobic), we downloaded the PR2 database (v4.14) of 16S rRNA and 18S rRNA genes^65^. We dereplicated the PR2 database using CD-HIT v4.7 with command “cd-hit-est -i <input.file> -o <output.file> -n 9 -c 0.97 -aS 0.5 -d 0 -r 1 -g 1 -M 30000 -sc 1”. Dereplication reduced the initial 197,602 sequences to 33,627. In construction of databases, we followed the approach by Arranz et al^170^. For interpretation, we only focused on major known human and animal eukaryotic symbionts (346 species) which do not have sequenced organelle genome or do not have mitochondria, as well as, *Blastocystis* which has an extensive number of 18S rRNA reference sequences in PR2 database. Target eukaryotes are listed in **Table S2.** All other 16S rRNA and 18S rRNA genes were kept in the reference database for achieving competitive mapping leaving only high quality reads in the mapping profile of target species in **Table S8**. We manually annotated highly prevalent and known species which are components of diet or pathogens/parasites to facilitate further interpretation of obtained eukaryotic profiles. We followed pathogen definitions proposed by Jochum and Stecher^171^. The list of pathogenic and commensal protozoa and fungi^172^ was curated based on CDC^173^ and WHO documentation^174^.

### Metagenomic data

The metaEukaryome pipeline was applied to 578 studies (75,727 samples) of WMS samples included in SPIRE and Metalog databases^60,61^. Samples originated from all major habitats: human microbiome (217 studies, 44,021 samples), animal (82 studies, 8,907 samples), sea/ocean (19 studies, 3,184 samples), and other environments such as freshwater, soil, air and anthropogenic environments (259 studies, 19,615 samples). For more detail see **Figure 2A, Figure S2 and Table S3**. From the downstream analysis, all detections of human DNA were ignored because of data protection reasons^175^. We also ignored all putative contaminant species from interpretation for human samples and samples originating from other habitats. The list of putative contaminant species is available in **Table S4**.

### Benchmarking the performance of metaEukaryome using public datasets

To evaluate the performance of metaEukaryome, we collected public metagenomic samples with references indicating the presence or absence of specific eukaryotic species. From the study by Lokmer et al.^67^, we obtained fecal whole-metagenome shotgun (WMS) sequencing data accompanied by qPCR references for *Blastocystis*. For the three soil-transmitted helminths (STH)—*Ascaris lumbricoides*, *Trichuris trichiura*, and hookworm—we collected qPCR references and paired fecal WMS samples from the Rosa et al.^66^ study, and Kato-Katz technique-based detections and paired fecal WMS samples from the Tee et al.^68^ study. In addition to the public datasets, we also prepared 122 samples in which we performed fungal internal transcribed spacer (ITS) sequencing and fecal WMS^69^.

For the metagenomic samples of the datasets, we profiled eukaryotes using metaEukaryome as well as publicly available eukaryote detection tools: CORRAL^41^, MetaPhlAn4 v4.0.1^37^, taxaTarget^40^, and EukDetect v1.0.1^38^. We used default parameters for all profilers. For profilers that provided abundance data, we binarized the results into presence/absence. We also binarized references that provided abundance criteria and compared these with the profilers’ results to evaluate precision, recall, and F1 score.

### Association analysis of dietary questionnaire data and eukaryotes in the Japanese cohort

To validate the dietary signals detected from human gut metagenomes, we analyzed data from the Japanese 4D cohort, in which both shotgun metagenomic sequencing data and dietary questionnaire data were available for 4,198 individuals. Details of cohort recruitment, sample collection, and data generation have been described previously^73^. Briefly, dietary information was collected from each participant using a Food Frequency Questionnaire (FFQ), which recorded the consumption frequency of 22 food items using a seven-level ordinal scale. Fecal microbial DNA was sequenced on the Illumina HiSeq platform, with a target sequencing depth of approximately 1 Gb per sample. Given the relatively shallow sequencing depth, we applied more lenient thresholds for eukaryote detection in this cohort. For the mitochondria- and plastid DNA-based approach, a eukaryote was considered present when horizontal coverage exceeded 0.03. Eukaryote detection results were then aggregated at the genus level.

We then manually curated the eukaryotic taxa detected by metaEukaryome and identified taxa that were likely derived from dietary sources. Corresponding pairs of dietary items and detected eukaryotic taxa were then defined (e.g., chicken consumption and the detection of chicken DNA, or egg consumption and the detection of chicken DNA). A total of 28 diet–eukaryote pairs were analyzed. For each pair, logistic regression analysis was performed using the glm function in R v4.4, with the presence or absence of the corresponding eukaryotic taxon as the dependent variable and the FFQ consumption frequency (treated as a numeric variable) as the independent variable.

### Preprocessing of human gut eukaryote data

From the full set of samples with eukaryote profiling, we extracted human gut samples for downstream analyses of associations with income class, disease status, and the gut prokaryome. Metadata on sex, age, country, and disease status for these samples were obtained from Metalog. Because many eukaryotes are profiled from dietary DNA, and infants have dietary patterns that differ substantially from adults, infant (age < 1 yr) samples were excluded from the association analysis. Sequencing depth strongly influenced eukaryote detection (**Figure S12**). To reduce the impact of low sequencing depth, samples with fewer than 1×10⁷ reads were excluded from downstream analyses, and sequencing depth was additionally included as a covariate in the statistical models. In total, 15,499 human gut samples sequenced worldwide were included in the analysis. Human DNA detections were ignored after profiling with metaEukaryome. Presence/absence of individual eukaryotes was determined using horizontal coverage thresholds of 0.05 for organelle DNA-based profiling (MT, PL) and 0.5 for rRNA gene-based profiling (PR2). A more stringent threshold for PR2-based profiling was chosen to increase confidence and accuracy of the detections. Results from different databases used for profiling were merged at the genus level. Putative contaminant eukaryotic species were excluded prior to downstream analyses (**Table S4**).

### Comparison of gut eukaryotes across income classifications

We aggregated genus-level eukaryote profiles by country/region and retained geographies with ≥20 samples (n=38). For each country pair, we computed Spearman’s rank correlation (ρ) of the average genus-level eukaryotic profiles and converted it to a dissimilarity (*d* = (1 - ρ)/2). We performed metric multidimensional scaling (MDS) on the precomputed dissimilarity matrix to obtain two dimensions for visualization. Each country/region was assigned to the World Bank income classification based on 2024 gross national income (GNI) per capita^111^. Because high-income economies showed clear separation from the other three classes (low, lower-middle, and upper-middle income), we collapsed the categories into two groups: High income versus Middle and low income (the union of low, lower-middle, and upper-middle income).

To identify eukaryotic taxa differing between the two economy groups, we fit a mixed-effects logistic regression (binomial GLMM) for each taxon detected in at least 10 samples. The response variable was taxon presence/absence at the sample level. Fixed effects included economy class (High vs. Middle/low), sex (binary), and sequencing depth (log-transformed read count), and a random intercept for study accounted for between-study heterogeneity. Models were fit in R (lme4::glmer). We report the fixed-effect estimate for economy class with 95% confidence interval and corresponding p-value.

To compare detected eukaryotic counts between the two economy classes, we fit a zero-inflated negative binomial GLMM, similar to the logistic regression, with fixed effects for economy class, sex, and sequencing depth, and a random intercept for study. We estimated economy class-specific average detected counts by predicting (on the response scale) at the mean sequencing depth with random effects set to zero, then averaging female/male predictions using the observed sex proportions.

### Association of gut eukaryotes with host disease status

To investigate associations between eukaryotes and host disease status, we compiled human gut samples with disease metadata from Metalog^61^. Diseases were retained if ≥100 case samples and ≥100 study-matched control samples were available, and only studies that included both case and control samples were analyzed. For each disease, we compared cases and controls within the same study set. Eukaryotic genera were included if detected in ≥10% of combined case and control samples. For each taxon–disease pair, we fit a logistic regression model with disease status as the binary outcome. Predictors included eukaryote presence/absence, sex, sequencing depth, and study indicators (dummy-coded fixed effects) to account for between-study differences. We report the regression coefficient, 95% confidence interval, and p-value for the eukaryote presence term. Fisher’s exact test was performed considering only the association between eukaryote presence and disease status, without adjusting for sex, sequencing depth, or study effects.

### Association between eukaryotes and prokaryotes in the human gut

Prokaryotic profiling of human gut metagenomic samples was performed using mOTUs3 (v3.0.1)^44^, and the resulting taxonomy was converted to the GTDB taxonomy (r207)^176^. Prokaryotic alpha diversity was quantified using the Shannon index. To identify eukaryotic genera associated with alpha diversity, we fit a linear mixed-effects model for each taxon detected in at least 10 samples. The response variable was Shannon index, and fixed effects included eukaryote presence/absence, sex, and sequencing depth. A random intercept for study– country accounted for between-study heterogeneity. We report the estimated effect size for eukaryote presence together with its FDR corrected p-value.

For associations at the level of individual prokaryotic taxa, we modeled their relative abundances using zero-inflated beta regression^133^. Each eukaryote–prokaryote pair was evaluated with eukaryote presence/absence, sex, and sequencing depth as fixed effects, and study–country as a random intercept. Analyses were conducted across all prokaryotic taxonomic ranks (from domain to species). Eukaryotic genera detected in fewer than 10 samples were excluded from the downstream analysis.

For the protists (*Endolimax*, *Entamoeba*, *Blastocystis*, and *Dientamoeba*) and fungi (*Saccharomyces* and *Candida*) that showed widespread significant associations across prokaryotic taxa, we calculated Spearman correlation coefficients between their association coefficients across all prokaryotic species (**Figure 4C**, upper right half). Because these similarities could arise from co-occurrence among eukaryotes within the same samples, we repeated the association analysis using zero-inflated beta mixed models in which other protists or fungi were included as additional fixed effects. We then recalculated Spearman correlation coefficients using the resulting adjusted association coefficients (**Figure 4C**, lower left half).

To visualize results on the prokaryotic tree of life, we used the human gut microbiome–specific catalog HRGM v2^134^. Representative genomes in HRGM v2 were taxonomically annotated with GTDB r207 using GTDB-Tk (v2.1.1)^177^ to ensure consistency with the prokaryotic profiling. To identify uncultured clades within the human gut microbiome, we examined all internal nodes of the HRGM v2 phylogenetic tree and selected the highest nodes for which all descendant leaf nodes were found exclusively in MAGs (i.e., not observed in isolates). To avoid including small clades, which may simply reflect limited culturing efforts rather than genuine unculturability, we required each clade to contain at least 20 leaf nodes, and the lowest common ancestor derived from leaf node taxonomy had to be at the family level or higher. We therefore defined uncultured clades as those containing ≥ 20 species, classified at least at the family level, that were detected only in MAGs. The metabolic independence score was obtained from HRGM v2 and calculated based on the presence of 33 metabolism-related KEGG modules^136^. For GTDB species matched to two or more HRGM representative genomes, the average score was used.

### Detection of prokaryotes from DNA and RNA sequencing data of *Blastocystis* culture

We obtained publicly available RNA-seq (PRJNA1041441) and long-read DNA sequencing (PRJNA1084796) data from a study that performed both DNA and RNA sequencing on axenic and xenic *Blastocystis* cultures^138^. To assess the presence of gut prokaryotes in sequencing reads derived from *Blastocystis* cultures, we used HRGM v2^134^, a reference catalog specific to the human gut microbiome.

For the long-read nanopore sequencing data, reads were aligned to HRGM v2 reference genomes using minimap2^178^ with the ‘-ax map-ont’ option. We then used samtools to calculate locus-level coverage and derived both horizontal coverage, defined as the fraction of the genome covered by aligned reads, and average depth coverage. A species was considered present when its genome showed an average depth coverage greater than 1 and a horizontal coverage greater than 0.5.

For the short-read RNA-seq data, reads were aligned to HRGMv2 reference genomes using bowtie2^179^ with the ‘--fast’ option. For RNA-seq, a species was considered transcriptionally detected when both horizontal coverage and average depth coverage exceeded 0.01. For the two taxa highlighted in this study, *Candidatus* Borkfalkia ceftriaxoniphila and CAG-345 sp000433315, we further manually verified that transcription was broadly distributed across the genome and across multiple contigs (**Figure 5E–F**).

### Association of *Blastocystis*-associated prokaryotes with genomic and metabolic traits

For each prokaryotic species, the association coefficient for *Blastocystis* derived from the zero-inflated beta regression was used as a continuous measure of enrichment or depletion in *Blastocystis*-positive samples. Metabolic independence scores were obtained from HRGM v2 and were calculated based on the presence of 33 metabolism-related KEGG modules^136^. For GTDB species matched to more than two HRGM representative genomes, the mean metabolic independence score was used. Genome size was calculated as the mean genome or MAG size for each species using genomes provided in SPIRE. Pangenome openness estimates were obtained from a previous study^124^, and species-level growth-related traits were obtained from metaTraits^137^. Associations between the *Blastocystis* association coefficient and continuous traits, including metabolic independence, genome size, and pangenome openness, were assessed using Pearson’s correlation. For binary traits, associations were tested using logistic regression, with the trait as the response variable and the *Blastocystis* association coefficient as the predictor.

### KEGG module prevalence comparison

The presence or absence of individual KEGG modules^136^ in each prokaryotic species was obtained from HRGM v2^134^. Prokaryotic species were classified as enriched in *Blastocystis*-positive samples when the association coefficient was >0.1 with q < 0.05, and as depleted when the coefficient was <−0.1 with q < 0.05. For each KEGG module, we compared its prevalence between the enriched and depleted groups. Statistical significance was assessed using Fisher’s exact test based on a 2 × 2 contingency table of module presence or absence and enrichment group, and p values were corrected for multiple testing using the false discovery rate (FDR).

### KEGG module complementarity analysis

To assess KEGG module complementarity between *Blastocystis hominis* and the two uncultured bacteria detected by both DNA-seq and RNA-seq in the axenic culture, *Ca.* B. ceftriaxoniphila and CAG-345 sp000433315, we compiled genome-wide protein catalogs for each organism. For *B. hominis*, we downloaded the full predicted proteome of NCBI assembly GCA_039096935.1, which was assembled from the same sample in which *Ca.* B. ceftriaxoniphila and CAG-345 sp000433315 were detected^138^. For the two bacterial genomes, we used HRGMv2_0713 (*Ca.* B. ceftriaxoniphila) and HRGMv2_3440 (CAG-345 sp000433315) from HRGM v2^134^. We annotated KEGG ortholog groups for each genome using KofamKOALA^180^ and calculated KEGG module completeness with KEMET^181^. To evaluate potential metabolic complementation, we combined the KO list of *B. hominis* with that of *Ca.* B. ceftriaxoniphila or CAG-345 sp000433315 and recalculated module completeness for each combined proteome. If the combined completeness of a given KEGG module exceeded the completeness observed in one species alone, that species was considered to benefit from complementation by the partner. If module completeness increased for both species, the module was classified as benefiting both partners.

### Ethics declarations

The Japanese 4D project was approved by the ethics committees (approval no. NCGM-S-004586) of all participating institutions, including the National Center for Global Health and Medicine and Tokyo Medical University Hospital. The study was conducted in accordance with the Declaration of Helsinki and relevant ethical guidelines. Written informed consent was obtained from all participants prior to enrollment.

## Data availability

Results of eukaryotic profiling (**Table S3**) are available on the SPIRE website (https://spire.embl.de/eire/) and at doi:10.5281/zenodo.24104693. The metaEukaryome tool is available at https://github.com/grp-bork/meta_eukaryome_detect and doi:10.5281/zenodo.24199947, reference database for the metaEukaryome pipeline available at doi:10.5281/zenodo.24198236. The long-read DNA sequencing data from the *Blastocystis* Nand II culture analyzed in this study are publicly available under BioProject PRJNA1084796. The corresponding RNA-seq data are available under BioProject PRJNA1041441.

## Code availability

Code for data processing available at https://github.com/grp-bork/metaEukaryome_analysis.

## Acknowledgements

The authors are deeply grateful to P.B. for his guidance and contributions to this work and beyond. We thank Yan Ping Yuan for computational support; Abigail Lind and Katherine Pollard for providing comments on the manuscript, raw sequencing data and information on *Blastocystis* cultures; Felix Salim for feedback on the manuscript draft; and Kyle Ardis for testing installation and usage of the metaEukaryome pipeline. This work made use of EMBL IT Services high-performance computing resources (https://zenodo.org/records/12785830)^182^. The work was supported by: the National Research Foundation of Korea (NRF) grant funded by the Korea government (MSIT) (No. RS-2023-00240807); MicrobioS (ERC-AdG-669830); Dutch Research Council (NWO-KIC) Healthy from soil to gut: the effect of the soil microbiome on human health (MicroHealth); the Ministry of Science, Research and the Arts Baden-Württemberg (MWK) within the framework of LIBIS/de.NBI; the Deutsche Forschungsgemeinschaft (DFG, German Research Foundation)—project number 460129525 (NFDI4Microbiota); the European Research Council (ERC) under the European Union’s Horizon 2020 research and innovation programme (grant agreement No. 947317, ASymbEL). Japanese 4D project was partially supported by Grants-in-Aid for Research from the National Center for Global Health and Medicine (28-2401, 29-2001, 29-2004, 19A1011, 19A1022, 19A2015, and 29-1025), the Ministry of Health, Labour, and Welfare, Japan (grant number: 19HB1003), JSPS KAKENHI Grants (JP17K09365 and JP20K08366), the Smoking Research Foundation, the DANONE Research Grant, research funding from the Japan Dairy Association (J-Milk), the Takeda Science Foundation, and COCKPIT funding. The funding bodies had no role in the study design, data collection or analysis, decision to publish, or preparation of the manuscript.

## Author contributions

Conceptualization - OMM, CYK, SN, TSBS, MK, PB.

Data curation - OMM, CYK, SN, AF, PF, CS, MIK, SMR, EK, WA, RA, TVR, SK, DP, NN, AS, AHB, TSBS, MK.

Formal analysis - OMM, CYK, SN, MK.

Funding acquisition - PB, AHB, AS, YK, NU, NN.

Investigation - OMM, CYK, SN, AF, PF, CS, MIK, SMR, EK, WA, RA, TVR, SK, DP, AS, AHB, TSBS, MK.

Methodology - AF, OMM, CYK, SN, MK.

Project administration - PB, MK, OMM, CYK, SN, AHB. Resources - AF, PB, AHB, AS.

Software - AF, OMM. Supervision - MK, PB, AHB.

Validation - NN, YK, TH, NU, SN, CYK, OMM.

Visualization - CYK, SN, OMM.

Writing – original draft - OMM, CYK, SN, MK.

Writing – review & editing - OMM, CYK, SN, AF, PF, CS, MIK, SMR, EK, WA, RA, TVR, SK, DP, NN, AS, AHB, TSBS, MK.

## Declaration of generative AI and AI-assisted technologies in the writing process

During the preparation of this work, the authors used ChatGPT (OpenAI) and Claude (Anthropic) to improve language and clarity. After using these tools, the authors reviewed and edited the contents as needed and take full responsibility for the content of the article.

## Supplementary Figures

**Figure S1.**
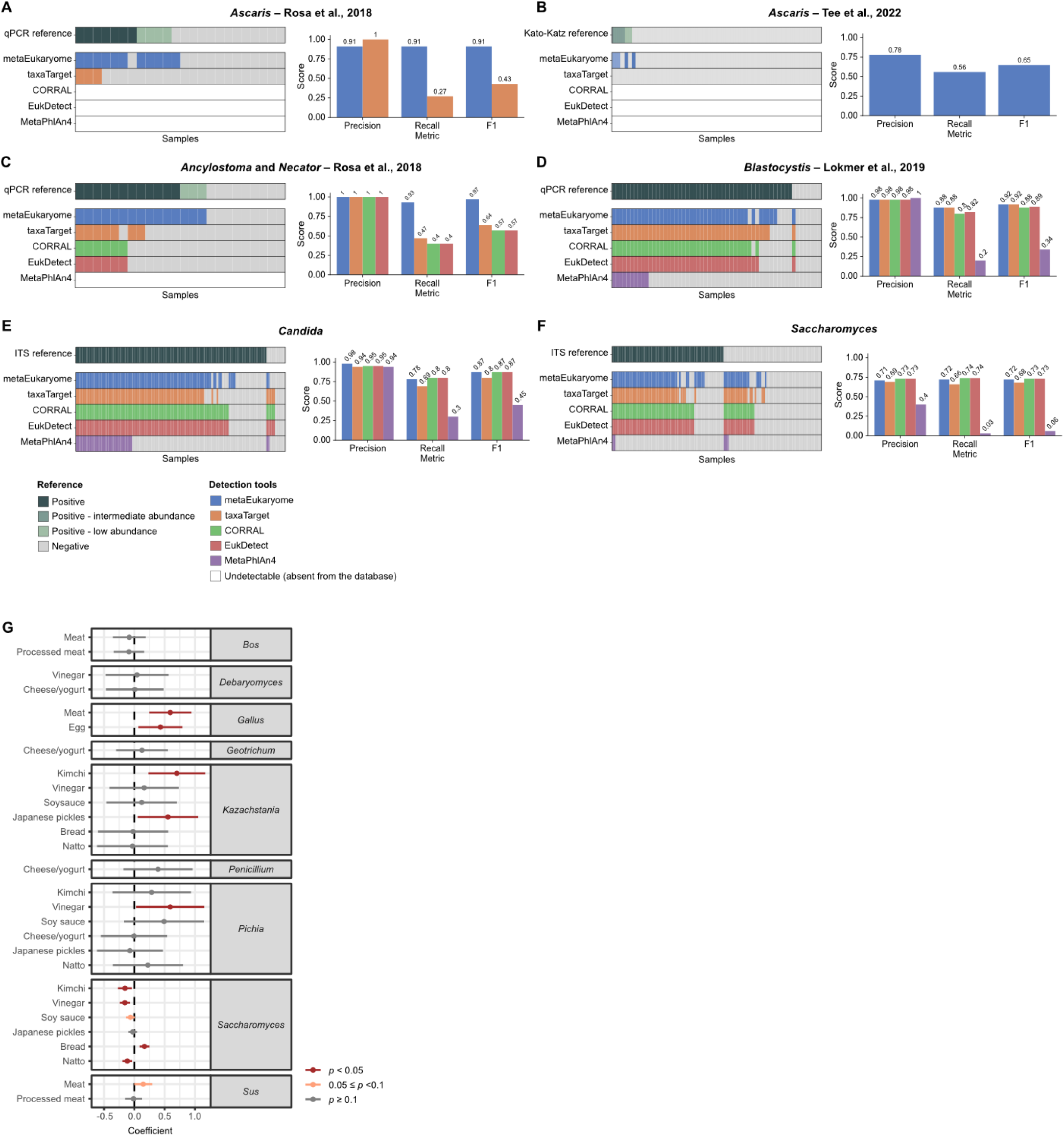
Benchmarking metaEukaryome across diverse eukaryotic lineages. Benchmarking of metaEukaryome against existing eukaryote detection tools using datasets in which the presence of target eukaryotic species had been independently established by non-metagenomic methods, including qPCR, Kato-Katz, and ITS sequencing. The benchmark spans diverse eukaryotic lineages, including (**A, B**) *Ascaris*, (**C**) hookworms, (**D**) *Blastocystis*, (**E**) *Candida*, and (**F**) *Saccharomyces*. Across these datasets, metaEukaryome covers the broadest taxonomic range and performs particularly well for worm detection. (**G**) Association between diet metadata entries and detections of corresponding eukaryotes in metagenomics data (logistic regression coefficient) in the Japanese 4D cohort (n = 4,198).

**Figure S2.**
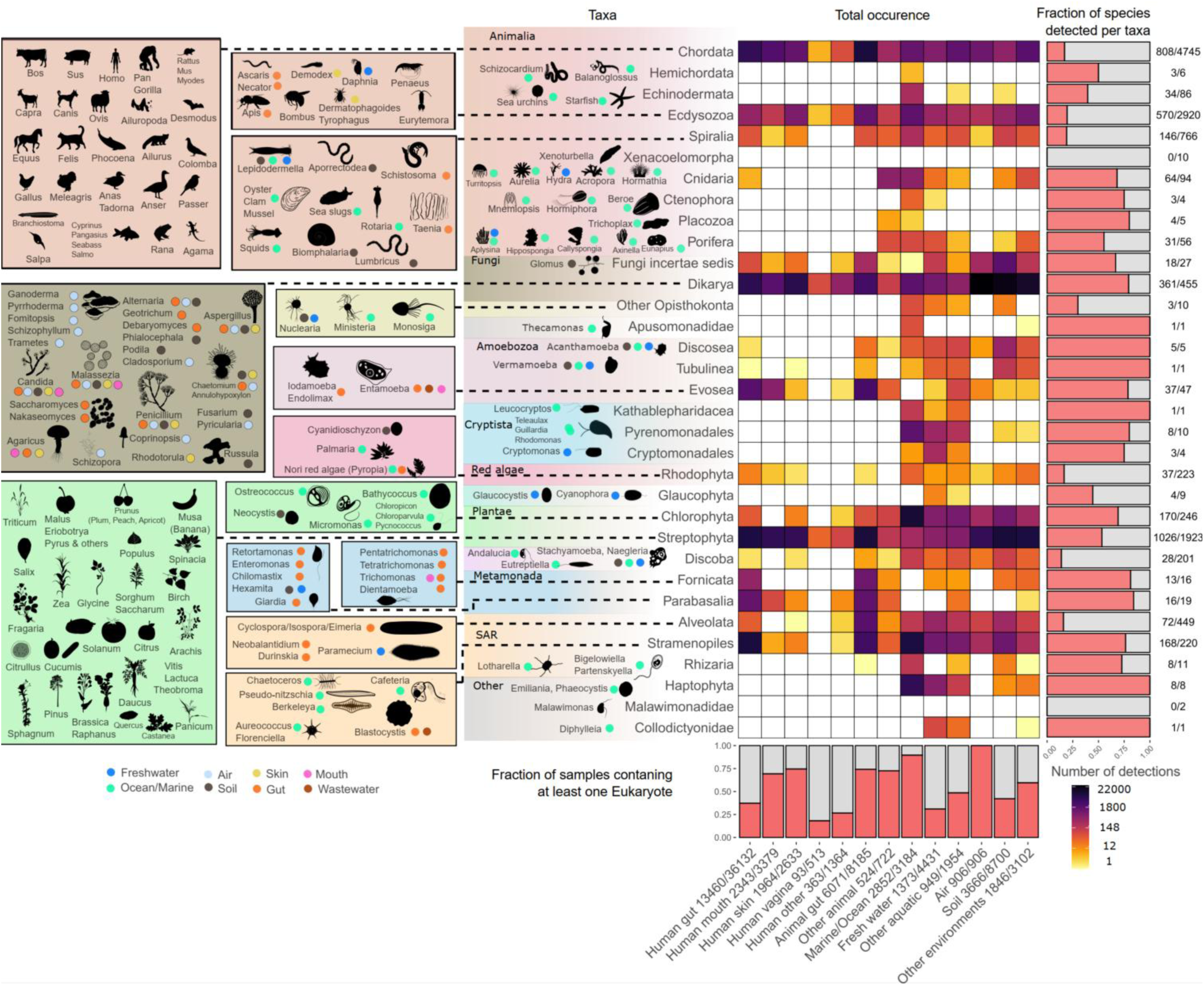
Overview of detectable eukaryotes in WMS data across all major habitats. The silhouette panel depicts examples of the most common eukaryotes that were detected in the specific habitats.

**Figure S3.**
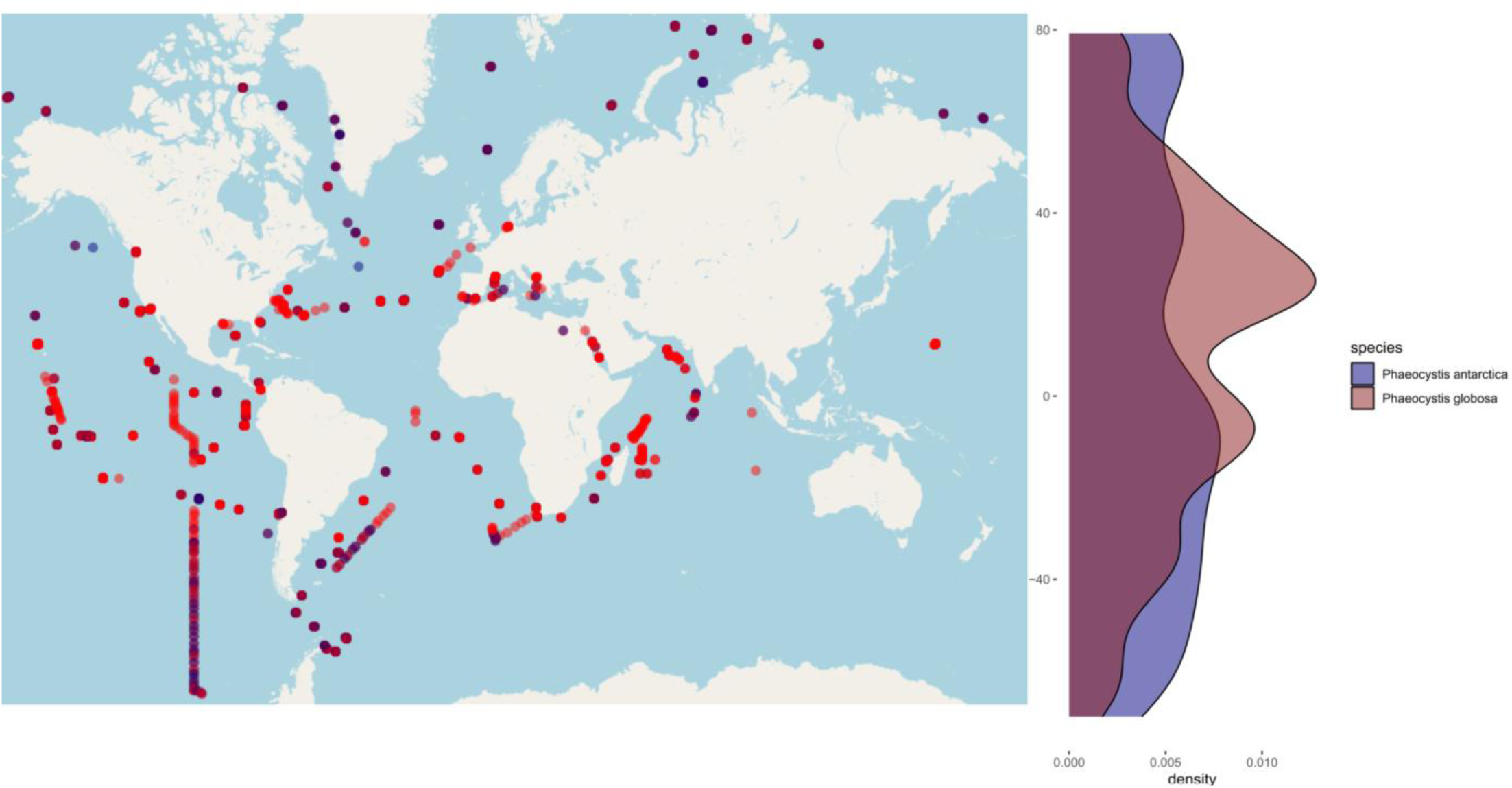
Distribution of *Phaeocystis* in the ocean. Dots represent locations of detections of *Phaeocystis antarctica* and *Phaeocystis globosa*. On the right side of the map density plot shows the distinct distribution of two species. *P. antarctica* is more prevalent in colder waters closer to polar regions.

**Figure S4.**
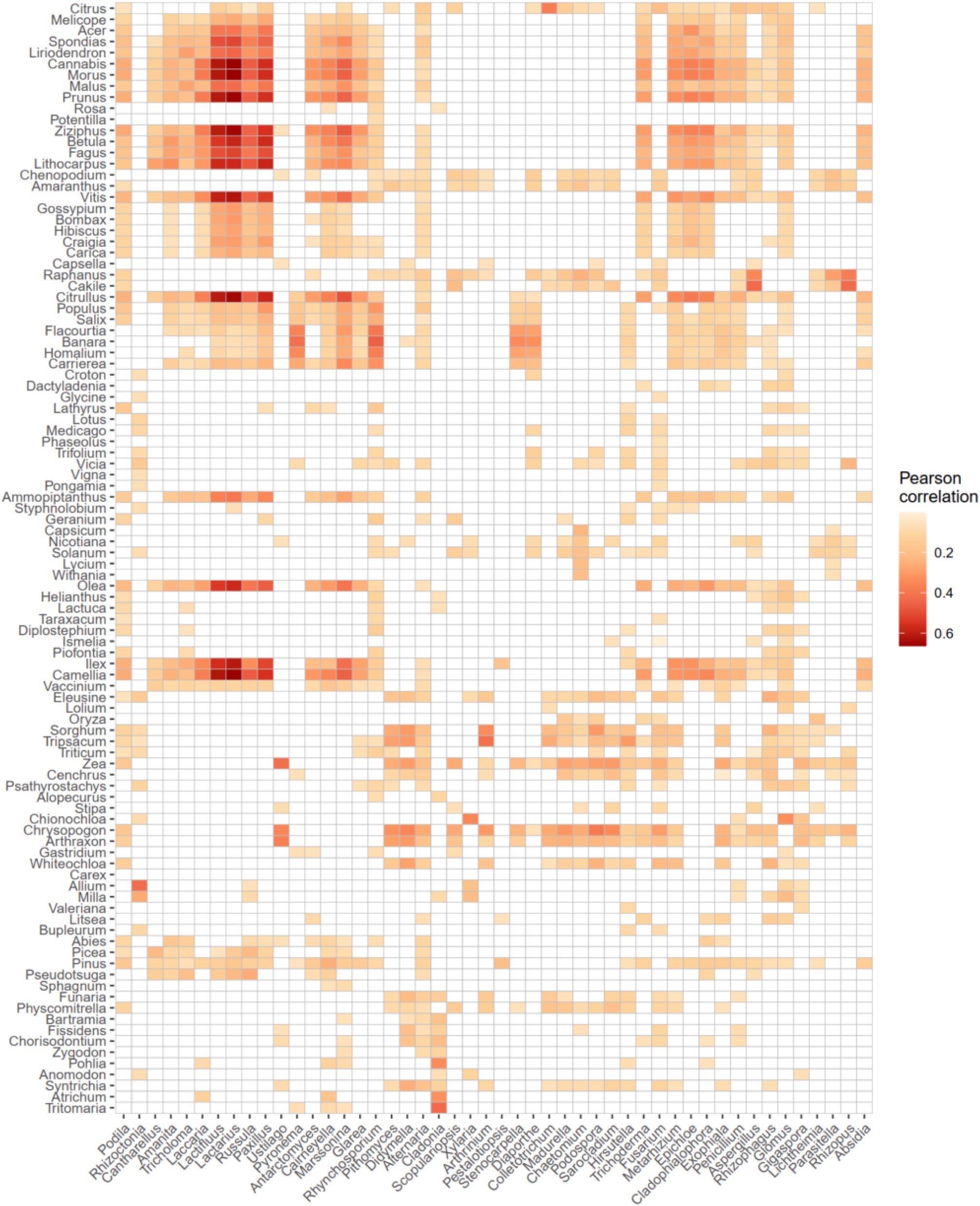
Associations between fungi and plants in soil. Strength of associations between plants and fungi across all soil samples in the dataset (average Pearson correlation at genus level across plant and fungi species clusters).

**Figure S5.**
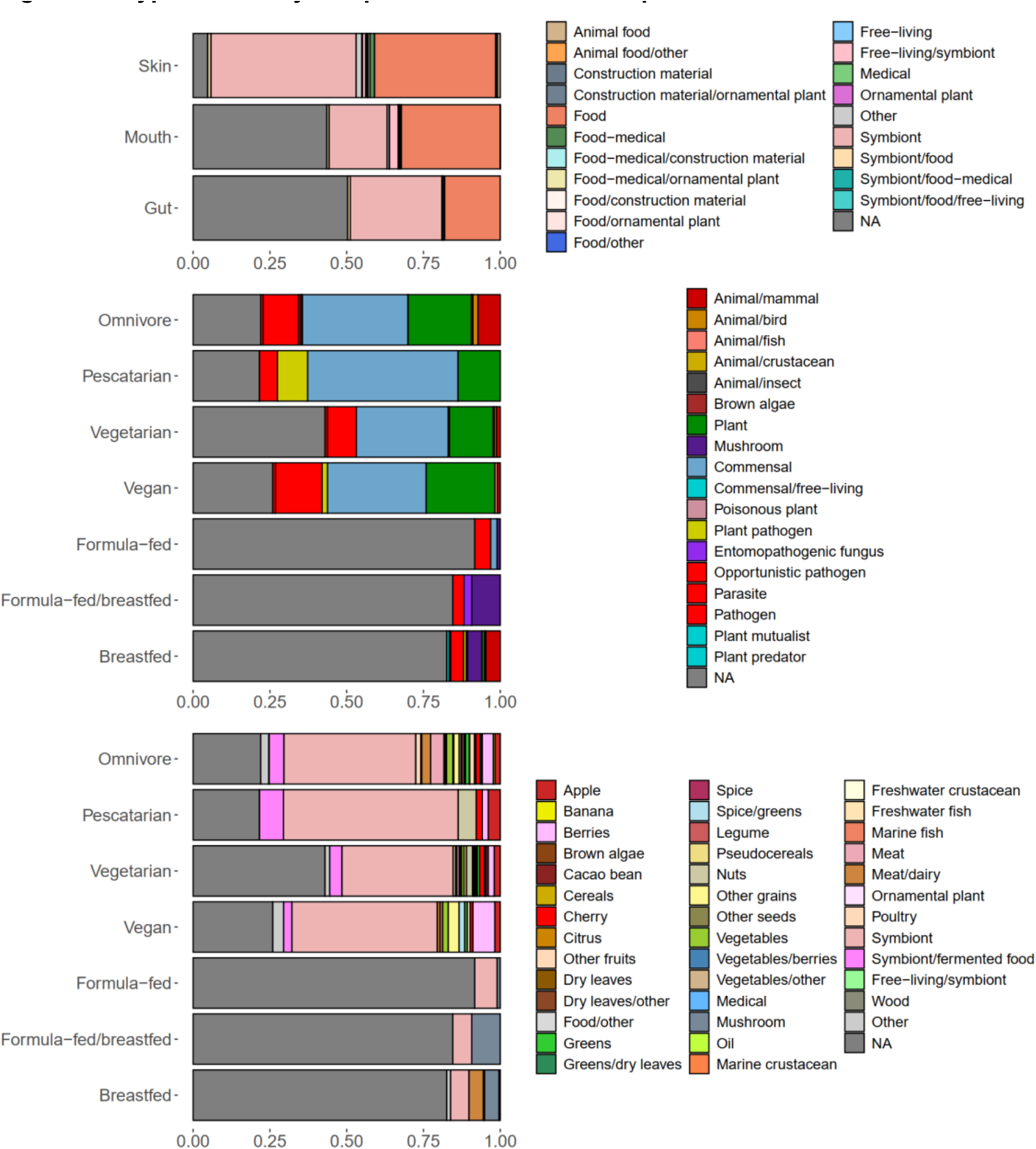
Types of eukaryotes prevalent in human samples. **(A)** Prevalence of main groups of eukaryotes in the most studied human bodysites (gut, mouth, skin). (**B–C)**. Relative prevalence of eukaryotes across types of diet reported in the metadata. NA indicates a fraction of samples without any confident detections of eukaryotes.

**Figure S6.**
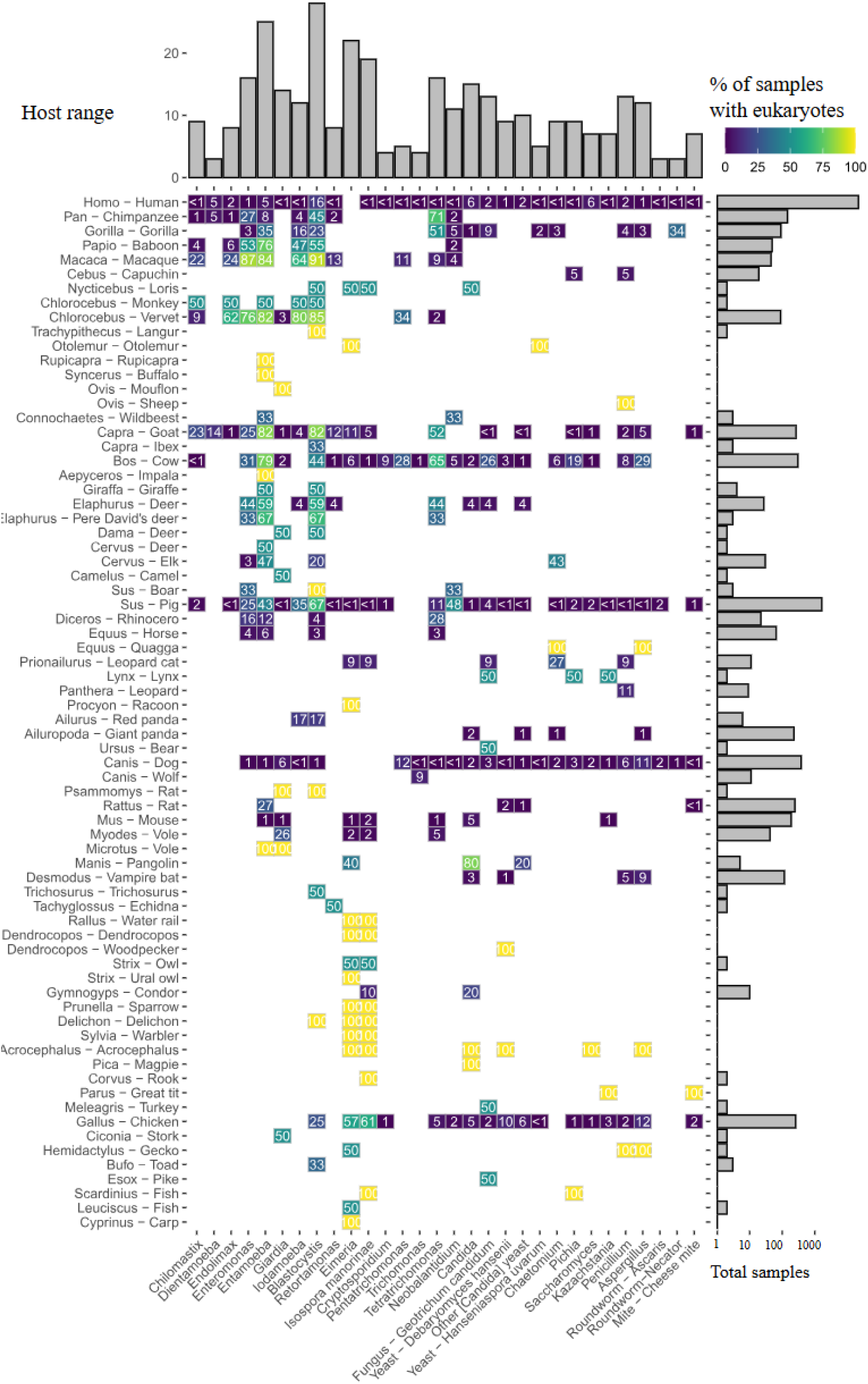
Pathogen burden and symbiont sharing between humans and all animals in the dataset (fecal samples) (extension of the Figure 2B).

**Figure S7.**
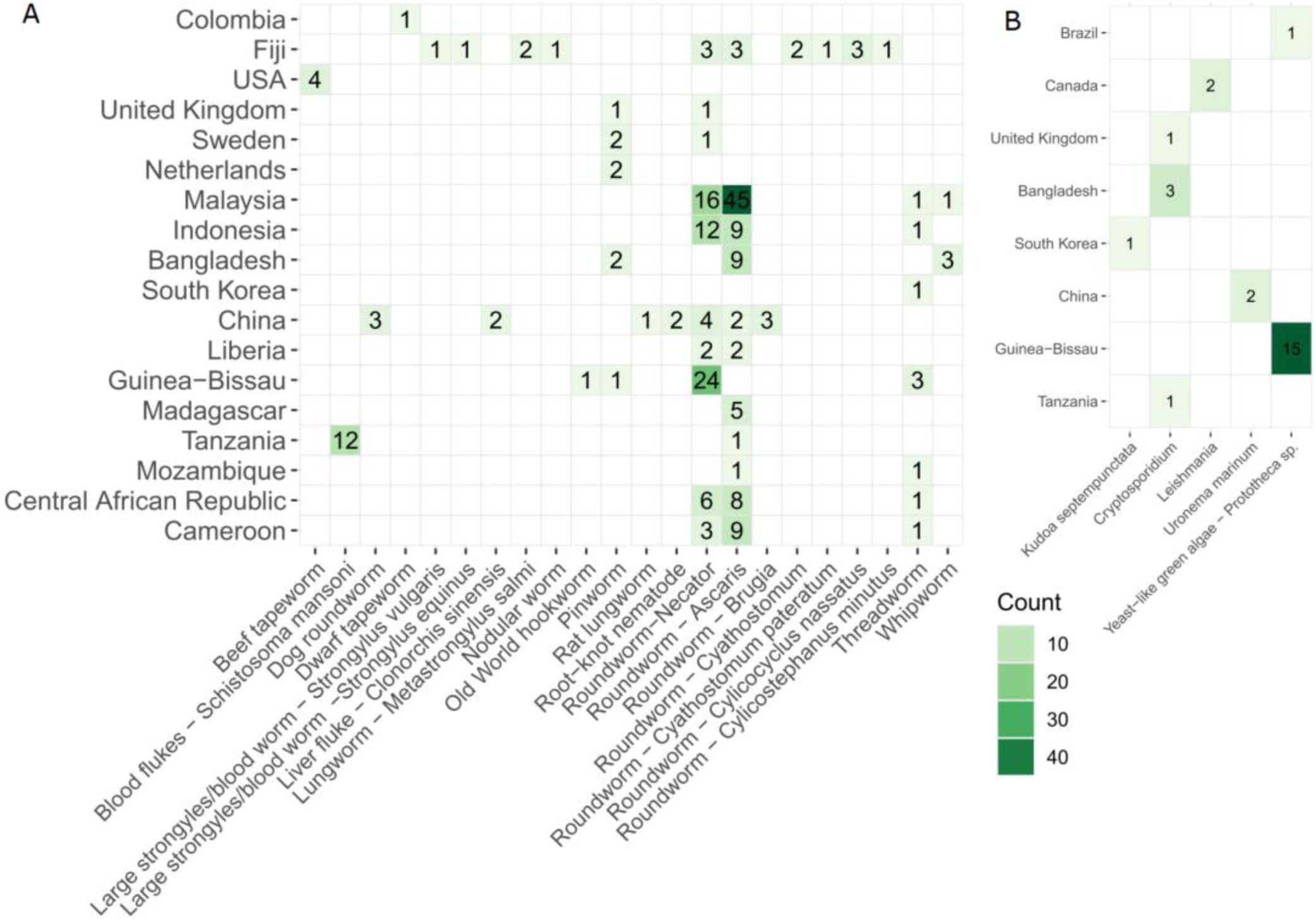
Examples of parasitic organisms in the human gut WMS samples. (**A**) Distribution of helminths in the human population. (**B**) Other rare pathogens and parasites in the human gut WMS samples.

**Figure S8.**
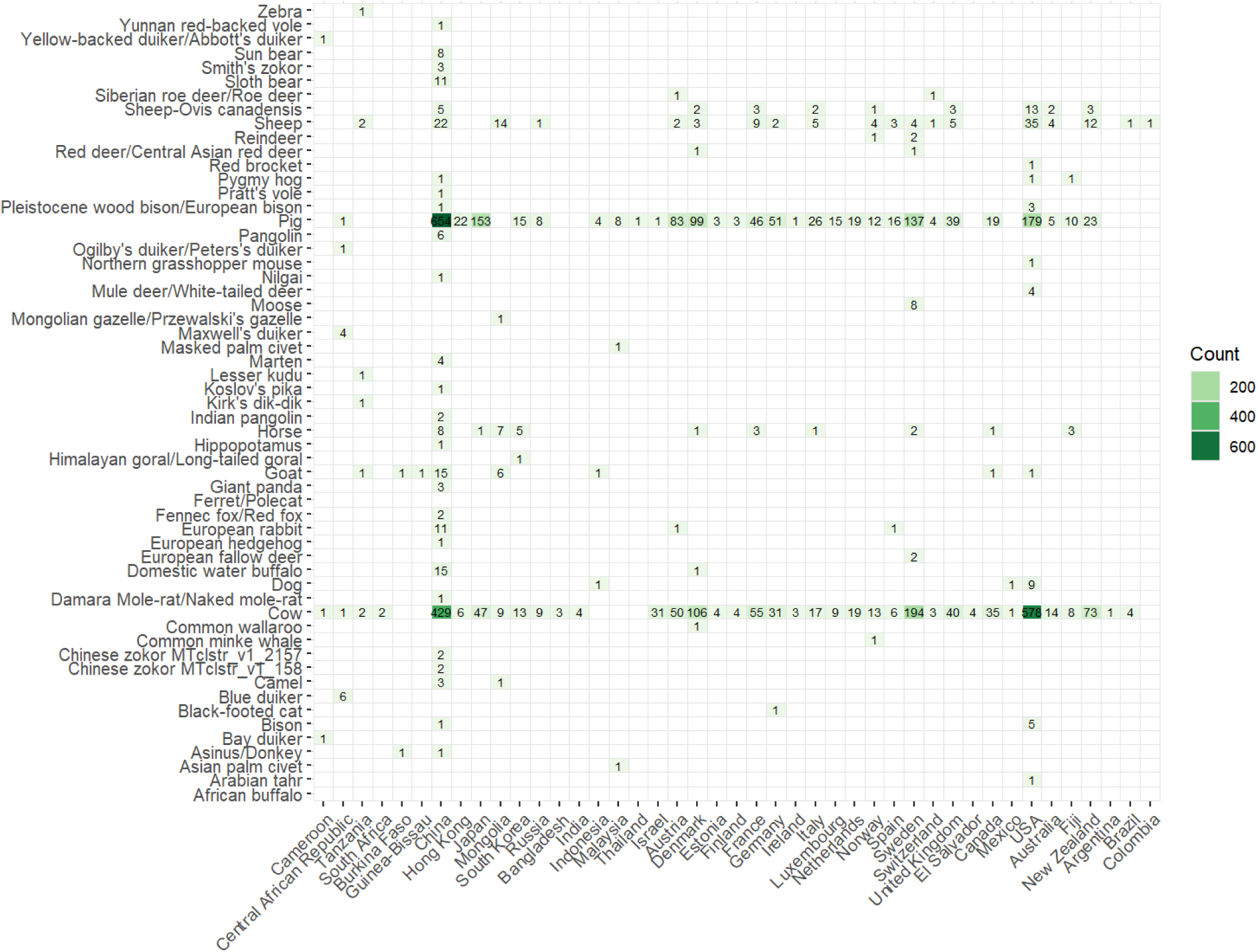
All detectable mammal species in the human gut WMS samples.

**Figure S9.**
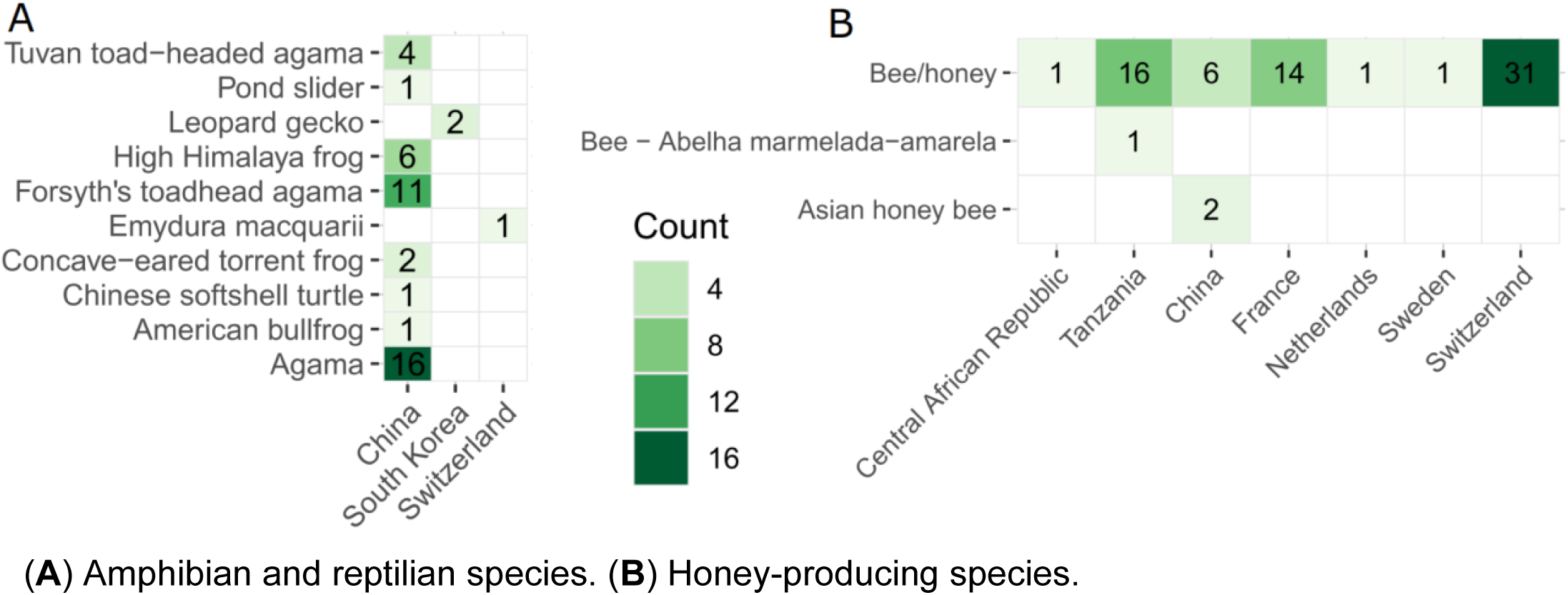
Rare food species in human gut WMS samples. (**A**) Amphibian and reptilian species. (**B**) Honey-producing species.

**Figure S10.**
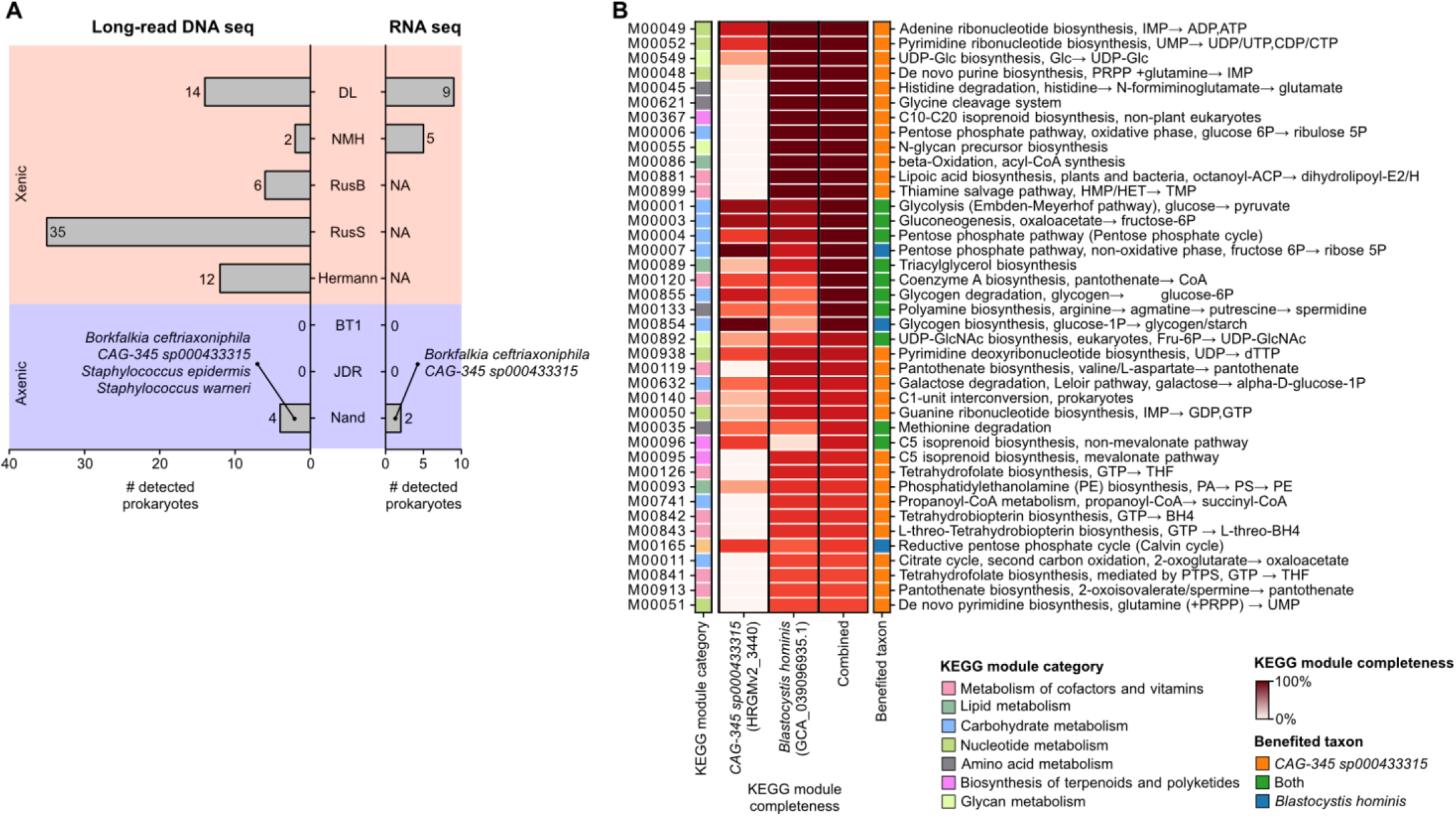
Detection of prokaryotic species in xenic and axenic *Blastocystis* cultures. (**A**) Numbers of prokaryotic species detected in DNA-seq and RNA-seq data from *Blastocystis* cultures. The red background indicates xenic *Blastocystis* cultures, whereas the blue background indicates axenic *Blastocystis* cultures. RNA-seq data were not available for the RusB, RusS, and Hermann strains. In the axenic NAHD II strain, *Ca.* Borkfalkia ceftriaxoniphila and CAG-345 sp000433315 were detected in both DNA-seq and RNA-seq data. (**B**) Complementarity of KEGG modules encoded by the genomes of *Blastocystis hominis* (NAND II) and CAG-345 sp000433315. The combined column indicates KEGG module completeness after pooling enzymes encoded by both genomes. Shown are the 40 complementary KEGG modules with the highest combined completeness, in which modules incomplete in one genome can be complemented by enzymes encoded in the other.

**Figure S11.**
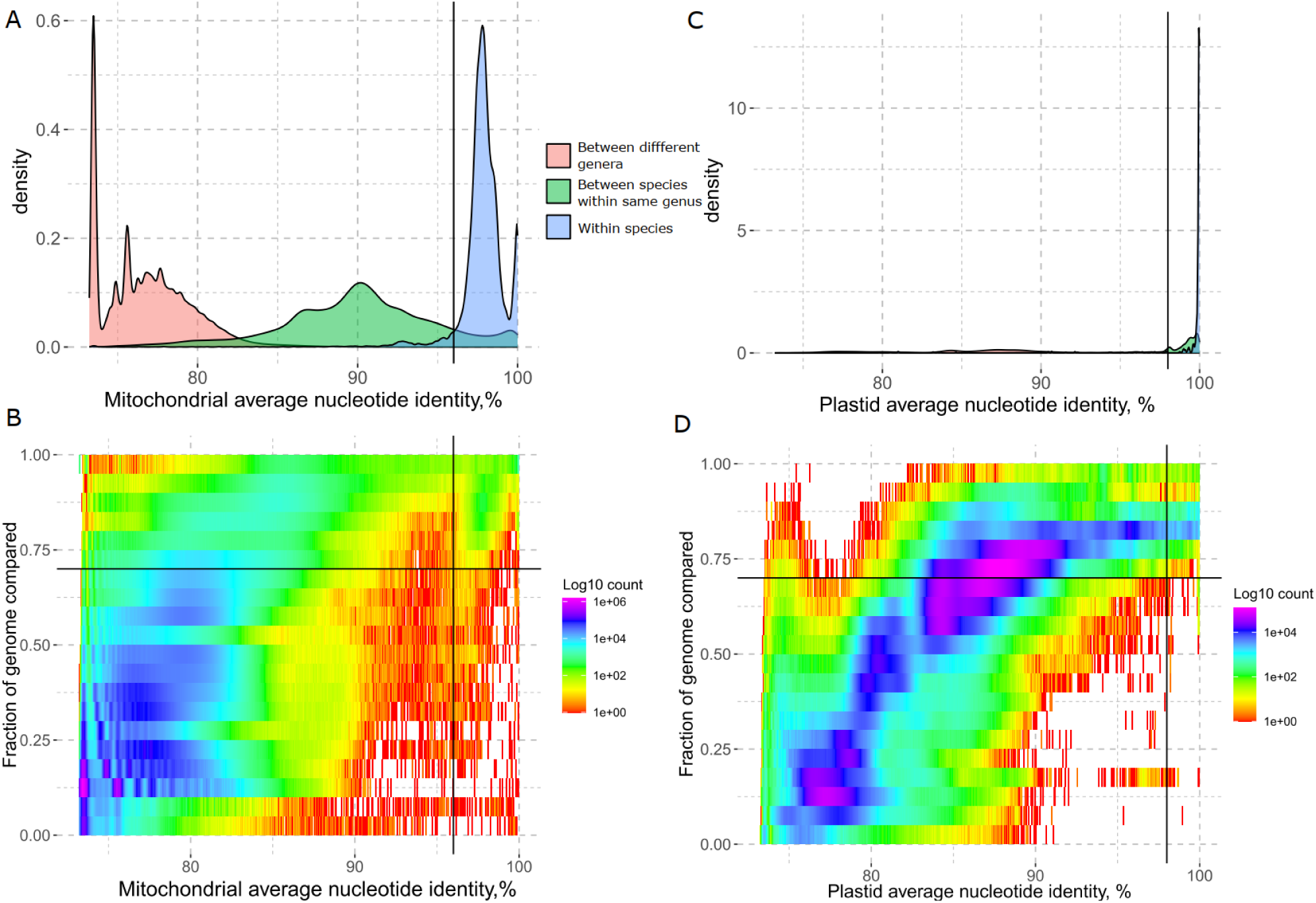
Determination of the threshold for dereplication of plastid and mitochondrial genomes. (**A**) Density of pairwise average nucleotide identity comparisons grouped by species, genus, family taxonomic levels in mitochondrial genomes. (**B**) Heatmap of density of the fraction of genome and average nucleotide identity for mitochondrial genomes. (**C**) Density of pairwise average nucleotide identity comparisons grouped by species, genus, family taxonomic levels in plastid genomes. (**D**) Heatmap of density of the fraction of genome and average nucleotide identity for plastid genomes.

**Figure S12.**
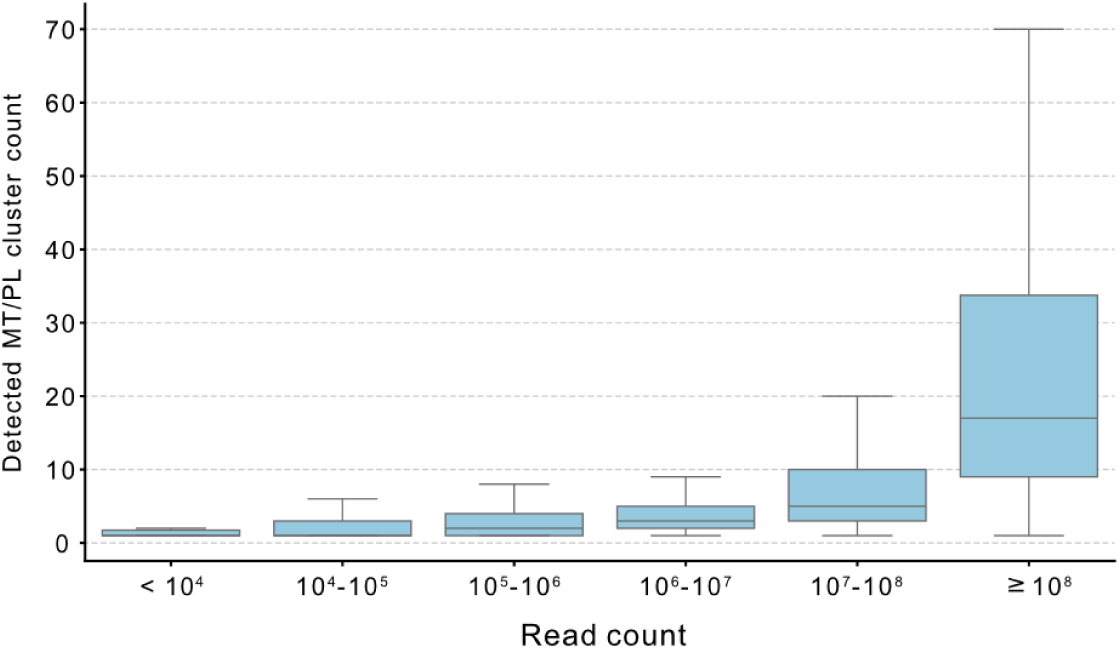
Effect of sequencing depth on eukaryote detection. Number of detected mitochondrial/plastid (MT/PL) species clusters across metagenomic samples grouped by sequencing depth. Detection increased with read count; samples with fewer than 1×10⁷ reads were excluded from downstream analyses.

## References

1. Lukeš, J., Stensvold, C.R., Jirků-Pomajbíková, K., and Parfrey, L.W. (2015). Are Human Intestinal Eukaryotes Beneficial or Commensals? PLOS Pathog. 11, e1005039. 10.1371/journal.ppat.1005039.

2. Bjorbækmo, M.F.M., Evenstad, A., Røsæg, L.L., Krabberød, A.K., and Logares, R. (2020). The planktonic protist interactome: where do we stand after a century of research? ISME J. 14, 544–559. 10.1038/s41396-019-0542-5.

3. Zoccarato, L., Sher, D., Miki, T., Segrè, D., and Grossart, H.-P. (2022). A comparative whole-genome approach identifies bacterial traits for marine microbial interactions. Commun. Biol. 5, 1–13. 10.1038/s42003-022-03184-4.

4. del Campo, J., Bass, D., and Keeling, P.J. (2020). The eukaryome: Diversity and role of microeukaryotic organisms associated with animal hosts. Funct. Ecol. 34, 2045–2054. 10.1111/1365-2435.13490.

5. Ianiro, G., Iorio, A., Porcari, S., Masucci, L., Sanguinetti, M., Perno, C.F., Gasbarrini, A., Putignani, L., and Cammarota, G. (2022). How the gut parasitome affects human health. Ther. Adv. Gastroenterol. 15, 17562848221091524. 10.1177/17562848221091524.

6. Bush, A., Sollmann, R., Wilting, A., Bohmann, K., Cole, B., Balzter, H., Martius, C., Zlinszky, A., Calvignac-Spencer, S., Cobbold, C.A., et al. (2017). Connecting Earth observation to high-throughput biodiversity data. Nat. Ecol. Evol. 1, 0176. 10.1038/s41559-017-0176.

7. Sparrow, B.D., Edwards, W., Munroe, S.E.M., Wardle, G.M., Guerin, G.R., Bastin, J.-F., Morris, B., Christensen, R., Phinn, S., and Lowe, A.J. (2020). Effective ecosystem monitoring requires a multi-scaled approach. Biol. Rev. 95, 1706–1719. 10.1111/brv.12636.

8. Jetz, W., Cavender-Bares, J., Pavlick, R., Schimel, D., Davis, F.W., Asner, G.P., Guralnick, R., Kattge, J., Latimer, A.M., Moorcroft, P., et al. (2016). Monitoring plant functional diversity from space. Nat. Plants 2, 16024. 10.1038/nplants.2016.24.

9. Rees, H.C., Maddison, B.C., Middleditch, D.J., Patmore, J.R.M., and Gough, K.C. (2014). REVIEW: The detection of aquatic animal species using environmental DNA – a review of eDNA as a survey tool in ecology. J. Appl. Ecol. 51, 1450–1459. 10.1111/1365-2664.12306.

10. Çevik, T., and Çevik, N. (2025). Environmental DNA (eDNA): A review of ecosystem biodiversity detection and applications. Biodivers. Conserv. 34, 2999–3035. 10.1007/s10531-025-03112-y.

11. Petrone, B.L., Aqeel, A., Jiang, S., Durand, H.K., Dallow, E.P., McCann, J.R., Dressman, H.K., Hu, Z., Tenekjian, C.B., Yancy, W.S., et al. (2023). Diversity of plant DNA in stool is linked to dietary quality, age, and household income. Proc. Natl. Acad. Sci. 120, e2304441120. 10.1073/pnas.2304441120.

12. Cabodevilla, X., Gómez-Moliner, B.J., Abad, N., and Madeira, M.J. (2023). Simultaneous analysis of the intestinal parasites and diet through eDNA metabarcoding. Integr. Zool. 18, 399–413. 10.1111/1749-4877.12634.

13. Bass, D., Stentiford, G.D., Littlewood, D.T.J., and Hartikainen, H. (2015). Diverse Applications of Environmental DNA Methods in Parasitology. Trends Parasitol. 31, 499–513. 10.1016/j.pt.2015.06.013.

14. Stensvold, C.R. (2024). Metabarcoding in gut protozoology. Trends Parasitol. 40, 1173–1182. 10.1016/j.pt.2024.10.015.

15. Martin, K.J., and Rygiewicz, P.T. (2005). Fungal-specific PCR primers developed for analysis of the ITS region of environmental DNA extracts. BMC Microbiol. 5, 28. 10.1186/1471-2180-5-28.

16. de Sousa, L.L., Silva, S.M., and Xavier, R. (2019). DNA metabarcoding in diet studies: Unveiling ecological aspects in aquatic and terrestrial ecosystems. Environ. DNA 1, 199–214. 10.1002/edn3.27.

17. de Vargas, C., Audic, S., Henry, N., Decelle, J., Mahé, F., Logares, R., Lara, E., Berney, C., Le Bescot, N., Probert, I., et al. (2015). Eukaryotic plankton diversity in the sunlit ocean. Science 348, 1261605. 10.1126/science.1261605.

18. Hord, N.G. (2008). Eukaryotic-Microbiota Crosstalk: Potential Mechanisms for Health Benefits of Prebiotics and Probiotics. Annu. Rev. Nutr. 28, 215–231. 10.1146/annurev.nutr.28.061807.155402.

19. Holzapfel, W.H., Haberer, P., Snel, J., Schillinger, U., and Huis in’t Veld, J.H.J. (1998). Overview of gut flora and probiotics. Int. J. Food Microbiol. 41, 85–101. 10.1016/S0168-1605(98)00044-0.

20. Vargas-Albores, F., Garibay-Valdez, E., Medina-Félix, D., and Martínez-Porchas, M. (2023). The micro-eukaryotic community: An underrated component of the mammalian gut microbiota? Front. Microbiol. 14. 10.3389/fmicb.2023.1123513.

21. Laforest-Lapointe, I., and Arrieta, M.-C. (2018). Microbial Eukaryotes: a Missing Link in Gut Microbiome Studies. mSystems 3, 10.1128/msystems.00201-17. 10.1128/msystems.00201-17.

22. Gerrick, E.R., and Howitt, M.R. (2025). The Lost Kingdom: commensal protists in the gut microbiota. Trends Microbiol. 33, 603–618. 10.1016/j.tim.2025.01.008.

23. Heredia, M.Y., and Knoll, L.J. (2026). Contributions of intestinal protists on the human gut landscape through the lens of Entamoeba spp. mSphere 11, e00011–26. 10.1128/msphere.00011-26.

24. Lind, A. (2025). Evolution and ecology of commensal gut protists: recent advances. Curr. Opin. Genet. Dev. 94, 102382. 10.1016/j.gde.2025.102382.

25. Saraiva, J.P., Bartholomäus, A., Toscan, R.B., Baldrian, P., and Nunes da Rocha, U. (2023). Recovery of 197 eukaryotic bins reveals major challenges for eukaryote genome reconstruction from terrestrial metagenomes. Mol. Ecol. Resour. 23, 1066–1076. 10.1111/1755-0998.13776.

26. Steinegger, M., and Salzberg, S.L. (2020). Terminating contamination: large-scale search identifies more than 2,000,000 contaminated entries in GenBank. Genome Biol. 21, 115. 10.1186/s13059-020-02023-1.

27. Pronk, L.J.U., and Medema, M.H. (2022). Whokaryote: distinguishing eukaryotic and prokaryotic contigs in metagenomes based on gene structure. Microb. Genomics 8, 000823. 10.1099/mgen.0.000823.

28. Karlicki, M., Antonowicz, S., and Karnkowska, A. (2022). Tiara: deep learning-based classification system for eukaryotic sequences. Bioinformatics 38, 344–350. 10.1093/bioinformatics/btab672.

29. Hou, S., Tang, T., Cheng, S., Liu, Y., Xia, T., Chen, T., Fuhrman, J.A., and Sun, F. (2024). DeepMicroClass sorts metagenomic contigs into prokaryotes, eukaryotes and viruses. NAR Genomics Bioinforma. 6, lqae044. 10.1093/nargab/lqae044.

30. West, P.T., Probst, A.J., Grigoriev, I.V., Thomas, B.C., and Banfield, J.F. (2018). Genome-reconstruction for eukaryotes from complex natural microbial communities. Genome Res. 28, 569–580. 10.1101/gr.228429.117.

31. plastiC: A pipeline for recovery and … | Wellcome Open Research https://wellcomeopenresearch.org/articles/8-475.

32. Tong, Y., Marcelino, V.R., Turnbull, R., and Verbruggen, H. (2026). ChloroScan: Recovering Plastid Genome Bins From Metagenomic Data. Mol. Ecol. Resour. 26, e70143. 10.1111/1755-0998.70143.

33. MitoZ: a toolkit for animal mitochondrial genome assembly, annotation and visualization | Nucleic Acids Research | Oxford Academic https://academic.oup.com/nar/article/47/11/e63/5377471?guestAccessKey=.

34. Marcelino, V.R., Clausen, P.T.L.C., Buchmann, J.P., Wille, M., Iredell, J.R., Meyer, W., Lund, O., Sorrell, T.C., and Holmes, E.C. (2020). CCMetagen: comprehensive and accurate identification of eukaryotes and prokaryotes in metagenomic data. Genome Biol. 21, 103. 10.1186/s13059-020-02014-2.

35. Krinos, A.I., Mars Brisbin, M., Hu, S.K., Cohen, N.R., Rynearson, T.A., Follows, M.J., Schulz, F., and Alexander, H. (2024). Missing microbial eukaryotes and misleading meta-omic conclusions. Nat. Commun. 15, 9873. 10.1038/s41467-024-52212-w.

36. Zhao, D., Salas-Leiva, D.E., Williams, S.K., Dunn, K.A., Shao, J.D., and Roger, A.J. (2025). Eukfinder: a pipeline to retrieve microbial eukaryote genome sequences from metagenomic data. mBio 16, e00699–25. 10.1128/mbio.00699-25.

37. Blanco-Míguez, A., Beghini, F., Cumbo, F., McIver, L.J., Thompson, K.N., Zolfo, M., Manghi, P., Dubois, L., Huang, K.D., Thomas, A.M., et al. (2023). Extending and improving metagenomic taxonomic profiling with uncharacterized species using MetaPhlAn 4. Nat. Biotechnol. 41, 1633–1644. 10.1038/s41587-023-01688-w.

38. Lind, A.L., and Pollard, K.S. (2021). Accurate and sensitive detection of microbial eukaryotes from whole metagenome shotgun sequencing. Microbiome 9, 58. 10.1186/s40168-021-01015-y.

39. Shih, J.B., Zhao, C., Pollard, K.S., and Lind, A.L. (2026). Quantitative detection of gut microbial eukaryotes with EukDetect2 reveals global distribution of commensal protists and association with distinct microbial community structure. Preprint at bioRxiv, 10.64898/2026.06.24.734308 10.64898/2026.06.24.734308.

40. Commichaux, S., Javkar, K., Muralidharan, H.S., Ramachandran, P., Ottesen, A., Rand, H., and Pop, M. (2022). taxaTarget: Fast, sensitive, and precise classification of microeukaryotes in metagenomic data. Preprint at Research Square, 10.21203/rs.3.rs-1186624/v3 10.21203/rs.3.rs-1186624/v3.

41. Bazant, W., Blevins, A.S., Crouch, K., and Beiting, D.P. (2023). Improved eukaryotic detection compatible with large-scale automated analysis of metagenomes. Microbiome 11, 72. 10.1186/s40168-023-01505-1.

42. Pita, S., Myers, P.N., Johansen, J., Russel, J., Nielsen, M.C., Eklund, A.C., and Nielsen, H.B. (2024). CHAMP delivers accurate taxonomic profiles of the prokaryotes, eukaryotes, and bacteriophages in the human microbiome. Front. Microbiol. 15. 10.3389/fmicb.2024.1425489.

43. Levy Karin, E., Mirdita, M., and Söding, J. (2020). MetaEuk—sensitive, high-throughput gene discovery, and annotation for large-scale eukaryotic metagenomics. Microbiome 8, 48. 10.1186/s40168-020-00808-x.

44. Ruscheweyh, H.-J., Milanese, A., Paoli, L., Karcher, N., Clayssen, Q., Keller, M.I., Wirbel, J., Bork, P., Mende, D.R., Zeller, G., et al. (2022). Cultivation-independent genomes greatly expand taxonomic-profiling capabilities of mOTUs across various environments. Microbiome 10, 212. 10.1186/s40168-022-01410-z.

45. Wood, D.E., Lu, J., and Langmead, B. (2019). Improved metagenomic analysis with Kraken 2. Genome Biol. 20, 257. 10.1186/s13059-019-1891-0.

46. Ngwese, M.M., Adegbite, B.R., Zinsou, J.F., Fitzstevens, J.L., Schmidt, V.T., Moure, P.A.N., Maloum, M.N., Tyakht, A.V., Huus, K.E., Youngblut, N.D., et al. (2025). Infection with gut parasites correlates with gut microbiome diversity across human populations in Africa. Gut Microbes 17, 2587966. 10.1080/19490976.2025.2587966.

47. Woodcroft, B.J., Aroney, S.T.N., Zhao, R., Cunningham, M., Mitchell, J.A.M., Nurdiansyah, R., Blackall, L., and Tyson, G.W. (2025). Comprehensive taxonomic identification of microbial species in metagenomic data using SingleM and Sandpiper. Nat. Biotechnol., 1–6. 10.1038/s41587-025-02738-1.

48. Diener, C., Holscher, H.D., Filek, K., Corbin, K.D., Moissl-Eichinger, C., and Gibbons, S.M. (2025). Metagenomic estimation of dietary intake from human stool. Nat. Metab. 7, 617–630. 10.1038/s42255-025-01220-1.

49. Krinos, A.I., Hu, S.K., Cohen, N.R., and Alexander, H. (2021). EUKulele: Taxonomic annotation of the unsung eukaryotic microbes. J. Open Source Softw. 6, 2817. 10.21105/joss.02817.

50. Beghini, F., Pasolli, E., Truong, T.D., Putignani, L., Cacciò, S.M., and Segata, N. (2017). Large-scale comparative metagenomics of Blastocystis, a common member of the human gut microbiome. ISME J. 11, 2848–2863. 10.1038/ismej.2017.139.

51. Piperni, E., Nguyen, L.H., Manghi, P., Kim, H., Pasolli, E., Andreu-Sánchez, S., Arrè, A., Bermingham, K.M., Blanco-Míguez, A., Manara, S., et al. (2024). Intestinal Blastocystis is linked to healthier diets and more favorable cardiometabolic outcomes in 56,989 individuals from 32 countries. Cell 187, 4554–4570.e18. 10.1016/j.cell.2024.06.018.

52. Maixner, F., Turaev, D., Cazenave-Gassiot, A., Janko, M., Krause-Kyora, B., Hoopmann, M.R., Kusebauch, U., Sartain, M., Guerriero, G., O’Sullivan, N., et al. (2018). The Iceman’s Last Meal Consisted of Fat, Wild Meat, and Cereals. Curr. Biol. 28, 2348–2355.e9. 10.1016/j.cub.2018.05.067.

53. Maixner, F., Sarhan, M.S., Huang, K.D., Tett, A., Schoenafinger, A., Zingale, S., Blanco-Míguez, A., Manghi, P., Cemper-Kiesslich, J., Rosendahl, W., et al. (2021). Hallstatt miners consumed blue cheese and beer during the Iron Age and retained a non-Westernized gut microbiome until the Baroque period. Curr. Biol. 10.1016/j.cub.2021.09.031.

54. Alexander, H., Hu, S.K., Krinos, A.I., Pachiadaki, M., Tully, B.J., Neely, C.J., and Reiter, T. (2023). Eukaryotic genomes from a global metagenomic data set illuminate trophic modes and biogeography of ocean plankton. mBio 14, e01676–23. 10.1128/mbio.01676-23.

55. Shrestha, B., Romero, M.F., Villada, J.C., Consortium, P.M., Blaby-Haas, C.E., and Schulz, F. (2025). Global Metagenomics Reveals Plastid Diversity and Unexplored Algal Lineages. Preprint at bioRxiv, 10.1101/2025.03.28.644651 10.1101/2025.03.28.644651.

56. Romero, M.F., Krinos, A.I., Maurer-Alcala, X., Burns, J.A., Stepanauskas, R., Woyke, T., and Schulz, F. (2026). Global Metagenomics Reveals Hidden Protist Diversity. Preprint at bioRxiv, 10.1101/2025.03.26.645542 10.1101/2025.03.26.645542.

57. Weyrich, L.S., Duchene, S., Soubrier, J., Arriola, L., Llamas, B., Breen, J., Morris, A.G., Alt, K.W., Caramelli, D., Dresely, V., et al. (2017). Neanderthal behaviour, diet, and disease inferred from ancient DNA in dental calculus. Nature 544, 357–361. 10.1038/nature21674.

58. Valdés-Mas, R., Leshem, A., Zheng, D., Cohen, Y., Kern, L., Zmora, N., He, Y., Katina, C., Eliyahu-Miller, S., Yosef-Hevroni, T., et al. (2025). Metagenome-informed metaproteomics of the human gut microbiome, host, and dietary exposome uncovers signatures of health and inflammatory bowel disease. Cell 188, 1062–1083.e36. 10.1016/j.cell.2024.12.016.

59. Shaw, J., and Yu, Y.W. (2025). Rapid species-level metagenome profiling and containment estimation with sylph. Nat. Biotechnol. 43, 1348–1359. 10.1038/s41587-024-02412-y.

60. Schmidt, T.S.B., Fullam, A., Ferretti, P., Orakov, A., Maistrenko, O.M., Ruscheweyh, H.-J., Letunic, I., Duan, Y., Van Rossum, T., Sunagawa, S., et al. (2024). SPIRE: a Searchable, Planetary-scale mIcrobiome REsource. Nucleic Acids Res. 52, D777–D783. 10.1093/nar/gkad943.

61. Kuhn, M., Schmidt, T.S.B., Ferretti, P., Głazek, A., Robbani, S.M., Akanni, W., Fullam, A., Schudoma, C., Cetin, E., Hassan, M., et al. (2026). Metalog: curated and harmonised contextual data for global metagenomics samples. Nucleic Acids Res. 54, D826–D834. 10.1093/nar/gkaf1118.

62. Bary, A. de (1879). (De Gruyter) 10.1515/9783111471839.

63. Karnkowska, A., Vacek, V., Zubáčová, Z., Treitli, S.C., Petrželková, R., Eme, L., Novák, L., Žárský, V., Barlow, L.D., Herman, E.K., et al. (2016). A Eukaryote without a Mitochondrial Organelle. Curr. Biol. CB 26, 1274–1284. 10.1016/j.cub.2016.03.053.

64. León-Avila, G., and Tovar, J. (2004). Mitosomes of Entamoeba histolytica are abundant mitochondrion-related remnant organelles that lack a detectable organellar genome. Microbiology 150, 1245–1250. 10.1099/mic.0.26923-0.

65. Guillou, L., Bachar, D., Audic, S., Bass, D., Berney, C., Bittner, L., Boutte, C., Burgaud, G., De Vargas, C., Decelle, J., et al. (2012). The Protist Ribosomal Reference database (PR2): a catalog of unicellular eukaryote Small Sub-Unit rRNA sequences with curated taxonomy. Nucleic Acids Res. 41, D597–D604. 10.1093/nar/gks1160.

66. Rosa, B.A., Supali, T., Gankpala, L., Djuardi, Y., Sartono, E., Zhou, Y., Fischer, K., Martin, J., Tyagi, R., Bolay, F.K., et al. (2018). Differential human gut microbiome assemblages during soil-transmitted helminth infections in Indonesia and Liberia. Microbiome 6, 33. 10.1186/s40168-018-0416-5.

67. Lokmer, A., Cian, A., Froment, A., Gantois, N., Viscogliosi, E., Chabé, M., and Ségurel, L. (2019). Use of shotgun metagenomics for the identification of protozoa in the gut microbiota of healthy individuals from worldwide populations with various industrialization levels. PLOS ONE 14, e0211139. 10.1371/journal.pone.0211139.

68. Tee, M.Z., Er, Y.X., Easton, A.V., Yap, N.J., Lee, I.L., Devlin, J., Chen, Z., Ng, K.S., Subramanian, P., Angelova, A., et al. (2022). Gut microbiome of helminth-infected indigenous Malaysians is context dependent. Microbiome 10, 214. 10.1186/s40168-022-01385-x.

69. Gu, W., Broekhoven, A.G.C., Van Blerk, S., Keller, M.I., Kuhn, M., Arif, S., Insonere, J.-L.-M., Roulet, A., Schierwagen, R., Brol, M.J., et al. (2026). Mapping multi-site and longitudinal microbiome transiting from decompensated cirrhosis to acute-on-chronic liver failure. (In preparation).

70. Wang, J., and Davis, R.E. (2020). Ascaris. Curr. Biol. 30, R423–R425. 10.1016/j.cub.2020.02.064.

71. Bethony, J., Brooker, S., Albonico, M., Geiger, S.M., Loukas, A., Diemert, D., and Hotez, P.J. (2006). Soil-transmitted helminth infections: ascariasis, trichuriasis, and hookworm. The Lancet 367, 1521– 1532. 10.1016/S0140-6736(06)68653-4.

72. Doyle, S.R., Sankaranarayanan, G., Allan, F., Berger, D., Jimenez Castro, P.D., Collins, J.B., Crellen, T., Duque-Correa, M.A., Ellis, P., Jaleta, T.G., et al. (2019). Evaluation of DNA Extraction Methods on Individual Helminth Egg and Larval Stages for Whole-Genome Sequencing. Front. Genet. 10, 826. 10.3389/fgene.2019.00826.

73. Nagata, N., Nishijima, S., Miyoshi-Akiyama, T., Kojima, Y., Kimura, M., Aoki, R., Ohsugi, M., Ueki, K., Miki, K., Iwata, E., et al. (2022). Population-level Metagenomics Uncovers Distinct Effects of Multiple Medications on the Human Gut Microbiome. Gastroenterology 163, 1038–1052. 10.1053/j.gastro.2022.06.070.

74. Nisiotou, A.A., Spiropoulos, A.E., and Nychas, G.-J.E. (2007). Yeast Community Structures and Dynamics in Healthy and Botrytis-Affected Grape Must Fermentations. Appl. Environ. Microbiol. 73, 6705–6713. 10.1128/AEM.01279-07.

75. Bendif, E.M., Nevado, B., Wong, E.L.Y., Hagino, K., Probert, I., Young, J.R., Rickaby, R.E.M., and Filatov, D.A. (2019). Repeated species radiations in the recent evolution of the key marine phytoplankton lineage Gephyrocapsa. Nat. Commun. 10, 4234. 10.1038/s41467-019-12169-7.

76. Zingone, A., and Oksfeldt Enevoldsen, H. (2000). The diversity of harmful algal blooms: a challenge for science and management. Ocean Coast. Manag. 43, 725–748. 10.1016/S0964-5691(00)00056-9.

77. Chapman, R.L. (2013). Algae: the world’s most important “plants”—an introduction. Mitig. Adapt. Strateg. Glob. Change 18, 5–12. 10.1007/s11027-010-9255-9.

78. Breyer, E., Stix, C., Kilker, S., Roller, B.R.K., Panagou, F., Doebke, C., Amano, C., Saavedra, D.E.M., Coll-García, G., Steger-Mähnert, B., et al. (2025). The contribution of pelagic fungi to ocean biomass. Cell 188, 3992–4002.e13. 10.1016/j.cell.2025.05.004.

79. Rahimlou, S., Amend, A.S., and James, T.Y. (2025). Malassezia in environmental studies is derived from human inputs. mBio 16, e01142–25. 10.1128/mbio.01142-25.

80. Füssy, Z., Lampe, R.H., Arrigo, K.R., Barry, K., Brisbin, M.M., Brussaard, C.P.D., Decelle, J., de Vargas, C., DiTullio, G.R., Elbourne, L.D.H., et al. (2025). Genome-resolved biogeography of Phaeocystales, cosmopolitan bloom-forming algae. Nat. Commun. 16, 8559. 10.1038/s41467-025-63565-1.

81. Smith, W.O., and Trimborn, S. (2024). Phaeocystis: A Global Enigma. Annu. Rev. Mar. Sci. 16, 417– 441. 10.1146/annurev-marine-022223-025031.

82. Van Nuland, M.E., Averill, C., Stewart, J.D., Prylutskyi, O., Corrales, A., van Galen, L.G., Manley, B.F., Qin, C., Lauber, T., Mikryukov, V., et al. (2025). Global hotspots of mycorrhizal fungal richness are poorly protected. Nature 645, 414–422. 10.1038/s41586-025-09277-4.

83. Tedersoo, L., Drenkhan, R., Abarenkov, K., Anslan, S., Bahram, M., Bitenieks, K., Buegger, F., Gohar, D., Hagh-Doust, N., Klavina, D., et al. (2024). The influence of tree genus, phylogeny, and richness on the specificity, rarity, and diversity of ectomycorrhizal fungi. Environ. Microbiol. Rep. 16, e13253. 10.1111/1758-2229.13253.

84. Vánky, K. (2005). The smut fungi (Ustilaginomycetes) of Chrysopogon (Poaceae). Fungal Diversity, 177–187.

85. Abrego, N., Furneaux, B., Hardwick, B., Somervuo, P., Palorinne, I., Aguilar-Trigueros, C.A., Andrew, N.R., Babiy, U.V., Bao, T., Bazzano, G., et al. (2024). Airborne DNA reveals predictable spatial and seasonal dynamics of fungi. Nature 631, 835–842. 10.1038/s41586-024-07658-9.

86. Bonner, M., Fresno, M., Gironès, N., Guillén, N., and Santi-Rocca, J. (2018). Reassessing the Role of Entamoeba gingivalis in Periodontitis. Front. Cell. Infect. Microbiol. 8. 10.3389/fcimb.2018.00379.

87. Macias-Paz, I.U., Pérez-Hernández, S., Tavera-Tapia, A., Luna-Arias, J.P., Guerra-Cárdenas, J.E., and Reyna-Beltrán, E. (2023). *Candida albicans* the main opportunistic pathogenic fungus in humans. Rev. Argent. Microbiol. 55, 189–198. 10.1016/j.ram.2022.08.003.

88. Kainz, K., Bauer, M.A., Madeo, F., and Carmona-Gutierrez, D. Fungal infections in humans: the silent crisis. Microb. Cell.

89. Palmeira-de-Oliveira, A., Silva, B.M., Palmeira-de-Oliveira, R., Martinez-de-Oliveira, J., and Salgueiro, L. (2013). Are Plant Extracts a Potential Therapeutic Approach for Genital Infections? Curr. Med. Chem. 20, 2914–2928. 10.2174/09298673113209990007.

90. Camelliae sinensis non fermentatum folium - herbal medicinal product | European Medicines Agency (EMA) (2021). https://www.ema.europa.eu/en/medicines/herbal/camelliae-sinensis-non-fermentatum-folium.

91. Huang, R., Li, F., Zhou, Y., Zeng, Z., He, X., Fang, L., Pan, F., Chen, Y., Lin, J., Li, J., et al. (2020). Metagenome-wide association study of the alterations in the intestinal microbiome composition of ankylosing spondylitis patients and the effect of traditional and herbal treatment. J. Med. Microbiol. 69, 797–805. 10.1099/jmm.0.001107.

92. Fitzcharles, M.-A., Rampakakis, E., Sampalis, J., Shir, Y., Cohen, M., Starr, M., and Häuser, W. (2020). Medical Cannabis Use by Rheumatology Patients Following Recreational Legalization: A Prospective Observational Study of 1000 Patients in Canada. ACR Open Rheumatol. 2, 286–293. 10.1002/acr2.11138.

93. Efficacy and safety of cannabidiol followed by an open label add-on of tetrahydrocannabinol for the treatment of chronic pain in patients with rheumatoid arthritis or ankylosing spondylitis: protocol for a multicentre, randomised, placebo-controlled study | BMJ Open https://bmjopen.bmj.com/content/9/6/e028197.abstract.

94. Carbone, K., and Gervasi, F. (2022). An Updated Review of the Genus Humulus: A Valuable Source of Bioactive Compounds for Health and Disease Prevention. Plants 11, 3434. 10.3390/plants11243434.

95. Györke, A., Pop, L., and Cozma, V. (2013). Prevalence and distribution of Eimeria species in broiler chicken farms of different capacities. Parasite 20, 50. 10.1051/parasite/2013052.

96. Chrenkova, V., Vadkertiova, R., Vlachova, K., Babjuk, M., Lischke, R., Bebrova, E., and Hubacek, P. (2022). *Candida sojae*: First report of a human infection. J. Med. Mycol. 32, 101309. 10.1016/j.mycmed.2022.101309.

97. Kano, R. (2020). Emergence of Fungal-Like Organisms: Prototheca. Mycopathologia 185, 747–754. 10.1007/s11046-019-00365-4.

98. Herrera, G., Barragán, N., Luna, N., Martínez, D., De Martino, F., Medina, J., Niño, S., Páez, L., Ramírez, A., Vega, L., et al. (2020). An interactive database of Leishmania species distribution in the Americas. Sci. Data 7, 110. 10.1038/s41597-020-0451-5.

99. Kawai, T., Sekizuka, T., Yahata, Y., Kuroda, M., Kumeda, Y., Iijima, Y., Kamata, Y., Sugita-Konishi, Y., and Ohnishi, T. (2012). Identification of Kudoa septempunctata as the Causative Agent of Novel Food Poisoning Outbreaks in Japan by Consumption of Paralichthys olivaceus in Raw Fish. Clin. Infect. Dis. 54, 1046–1052. 10.1093/cid/cir1040.

100. Tracking diseases from the sewer - Joint Research Centre (2026). https://joint-research-centre.ec.europa.eu/jrc-news-and-updates/tracking-diseases-sewer-2025-01-29_en.

101. Xiao, K., and Zhang, L. (2023). Wastewater pathogen surveillance based on One Health approach. Lancet Microbe 4, e297. 10.1016/S2666-5247(23)00039-3.

102. Levy, J.I., Andersen, K.G., Knight, R., and Karthikeyan, S. (2023). Wastewater surveillance for public health. Science 379, 26–27. 10.1126/science.ade2503.

103. Tiwari, A., Radu, E., Kreuzinger, N., Ahmed, W., and Pitkänen, T. (2024). Key considerations for pathogen surveillance in wastewater. Sci. Total Environ. 945, 173862. 10.1016/j.scitotenv.2024.173862.

104. Kilaru, P., Hill, D., Anderson, K., Collins, M.B., Green, H., Kmush, B.L., and Larsen, D.A. (2023). Wastewater Surveillance for Infectious Disease: A Systematic Review. Am. J. Epidemiol. 192, 305–322. 10.1093/aje/kwac175.

105. Singh, S., Ahmed, A.I., Almansoori, S., Alameri, S., Adlan, A., Odivilas, G., Chattaway, M.A., Salem, S.B., Brudecki, G., and Elamin, W. (2024). A narrative review of wastewater surveillance: pathogens of concern, applications, detection methods, and challenges. Front. Public Health 12. 10.3389/fpubh.2024.1445961.

106. Grassly, N.C., Shaw, A.G., and Owusu, M. (2025). Global wastewater surveillance for pathogens with pandemic potential: opportunities and challenges. Lancet Microbe 6. 10.1016/j.lanmic.2024.07.002.

107. Santos, M.C. dos, Silva, A.C.C., Teixeira, C. dos R., Prazeres, F.P.M., Santos, R.F. dos, Rolo, C. de A., Santos, E. de S., Fonseca, M.S. da, Valente, C.O., Hodel, K.V.S., et al. (2024). Wastewater surveillance for viral pathogens: A tool for public health. Heliyon 10. 10.1016/j.heliyon.2024.e33873.

108. Capone, D., Chiluvane, M., Cumbane, V., Dalton, J., Holcomb, D., Kowalsky, E., Lai, A., Mataveia, E., Monteiro, V., Rao, G., et al. (2026). Environmental pathogen surveillance in cities without universal piped wastewater infrastructure. PLOS Glob. Public Health 6, e0004994. 10.1371/journal.pgph.0004994.

109. Carlson, C.J., Brookson, C.B., Becker, D.J., Cummings, C.A., Gibb, R., Halliday, F.W., Heckley, A.M., Huang, Z.Y.X., Lavelle, T., Robertson, H., et al. (2025). Pathogens and planetary change. Nat. Rev. Biodivers. 1, 32–49. 10.1038/s44358-024-00005-w.

110. Gardy, J.L., and Loman, N.J. (2018). Towards a genomics-informed, real-time, global pathogen surveillance system. Nat. Rev. Genet. 19, 9–20. 10.1038/nrg.2017.88.

111. World Bank Country and Lending Groups – World Bank Data Help Desk https://datahelpdesk.worldbank.org/knowledgebase/articles/906519-world-bank-country-and-lending-groups.

112. Time to stop referring to the “developing world” World Bank Blogs. https://blogs.worldbank.org/en/opendata/time-stop-referring-developing-world.

113. Bourgoin, G., Callait-Cardinal, M.-P., Bouhsira, E., Polack, B., Bourdeau, P., Roussel Ariza, C., Carassou, L., Lienard, E., and Drake, J. (2022). Prevalence of major digestive and respiratory helminths in dogs and cats in France: results of a multicenter study. Parasit. Vectors 15, 314. 10.1186/s13071-022-05368-7.

114. Charlier, J., Rinaldi, L., Musella, V., Ploeger, H.W., Chartier, C., Vineer, H.R., Hinney, B., von Samson-Himmelstjerna, G., Băcescu, B., Mickiewicz, M., et al. (2020). Initial assessment of the economic burden of major parasitic helminth infections to the ruminant livestock industry in Europe. Prev. Vet. Med. 182, 105103. 10.1016/j.prevetmed.2020.105103.

115. Fu, J., Zhang, J., He, R., Dong, Q., Mao, H., Shen, W., Wu, W., Chen, X., Ma, W., Zhai, Q., et al. (2026). A global metagenomic atlas of aging identifies a microbiota phase transition associated with disease risk. Npj Biofilms Microbiomes 12, 97. 10.1038/s41522-026-00970-4.

116. Vaiserman, A.M., Koliada, A.K., and Marotta, F. (2017). Gut microbiota: A player in aging and a target for anti-aging intervention. Ageing Res. Rev. 35, 36–45. 10.1016/j.arr.2017.01.001.

117. López-Otín, C., Blasco, M.A., Partridge, L., Serrano, M., and Kroemer, G. (2023). Hallmarks of aging: An expanding universe. Cell 186, 243–278. 10.1016/j.cell.2022.11.001.

118. Ghosh, T.S., Shanahan, F., and O’Toole, P.W. (2022). The gut microbiome as a modulator of healthy ageing. Nat. Rev. Gastroenterol. Hepatol. 19, 565–584. 10.1038/s41575-022-00605-x.

119. Martino, C., Dilmore, A.H., Burcham, Z.M., Metcalf, J.L., Jeste, D., and Knight, R. (2022). Microbiota succession throughout life from the cradle to the grave. Nat. Rev. Microbiol. 20, 707–720. 10.1038/s41579-022-00768-z.

120. Ni, Z., Xie, J., Zhang, Z., Zhang, K., Ding, J., Chang, M., Su, Y., Zhang, Y., Yao, M., Wang, C., et al. Microbiome and aging: Trajectories of microbiome age across human ecosystems and their systemic effects. iMeta n/a, e70148. 10.1002/imt2.70148.

121. Ferretti, P., Wirbel, J., Maistrenko, O.M., Rossum, T.V., Alves, R., Fullam, A., Akanni, W., Schudoma, C., Schwarz, A., Thielemann, R., et al. (2023). C. difficile may be overdiagnosed in adults and is a prevalent commensal in infants. eLife 12. 10.7554/eLife.90111.1.

122. Zisova, L.G., Chokoeva, A.A., Amaliev, G.I., Petleshkova, P.V., Miteva-Katrandzhieva, T.М., Krasteva, M.B., Uchikova, E.H., Kouzmanov, A.H., and Ivanova, Z.V. (2016). Vulvovaginal Candidiasis in Pregnant Women and its Importance for Candida Colonization of Newborns. Folia Med. (Plovdiv) 58, 108–114. 10.1515/folmed-2016-0018.

123. Schnorr, S.L., Candela, M., Rampelli, S., Centanni, M., Consolandi, C., Basaglia, G., Turroni, S., Biagi, E., Peano, C., Severgnini, M., et al. (2014). Gut microbiome of the Hadza hunter-gatherers. Nat. Commun. 5, 3654. 10.1038/ncomms4654.

124. Kim, C.Y., Podlesny, D., Schiller, J., Khedkar, S., Fullam, A., Orakov, A., Schudoma, C., Robbani, S.M., Grekova, A., Kuhn, M., et al. (2026). Planetary microbiome structure and generalist-driven gene flow across disparate habitats. Cell 189, 2073–2091.e21. 10.1016/j.cell.2025.12.051.

125. Calderaro, A., Buttrini, M., Montecchini, S., Rossi, S., Farina, B., Arcangeletti, M.C., De Conto, F., and Chezzi, C. (2022). Prevalence of Intestinal Parasitoses in a Non-Endemic Setting during a 10-Year Period (2011-2020): A Focus on Dientamoeba fragilis. Microorganisms 10, 426. 10.3390/microorganisms10020426.

126. Hurych, J., Oscarsson, E., Håkanson, Å., Jirků-Pomajbíková, K., Jirků, M., Aronson, C.A., Cinek, O., Agardh, D., and HEDIMED Investigator Group (2023). Effects of Lactiplantibacillus plantarum and Lacticaseibacillus paracasei supplementation on the single-cell fecal parasitome in children with celiac disease autoimmunity: a randomized, double-blind placebo-controlled clinical trial. Parasit. Vectors 16, 411. 10.1186/s13071-023-06027-1.

127. Shi, W., Huang, X., Schooling, C.M., and Zhao, J.V. (2023). Red meat consumption, cardiovascular diseases, and diabetes: a systematic review and meta-analysis. Eur. Heart J. 44, 2626–2635. 10.1093/eurheartj/ehad336.

128. Hu, B., He, X., Sun, H., Hu, Y., Li, F., Sun, Y., Sun, J., and Feng, L. (2024). Red and processed meat intake and risk of cardiovascular disease: A two-sample Mendelian randomization study. Clin. Nutr. ESPEN 60, 289–297. 10.1016/j.clnesp.2024.02.014.

129. Wang, M., Gao, C., Lessing, D.J., and Chu, W. (2025). Saccharomyces cerevisiae SC-2201 Attenuates AOM/DSS-Induced Colorectal Cancer by Modulating the Gut Microbiome and Blocking Proinflammatory Mediators. Probiotics Antimicrob. Proteins 17, 1523–1535. 10.1007/s12602-024-10228-0.

130. Li, J.Q., Li, J.L., Xie, Y.H., Wang, Y., Shen, X.N., Qian, Y., Han, J.X., Chen, Y.X., and Fang, J.-Y. (2020). Saccharomyces cerevisiae may serve as a probiotic in colorectal cancer by promoting cancer cell apoptosis. J. Dig. Dis. 21, 571–582. 10.1111/1751-2980.12930.

131. Culpepper, T., Senthil, K., Vlcek, J., Hazelton, A., Heavey, M.K., Sellers, R.S., Nguyen, J., and Arthur, J.C. (2025). Engineered Probiotic Saccharomyces boulardii Reduces Colitis-Associated Colorectal Cancer Burden in Mice. Dig. Dis. Sci. 70, 2348–2367. 10.1007/s10620-025-09008-9.

132. Shamekhi, S., Abdolalizadeh, J., Ostadrahimi, A., Mohammadi, S.A., Barzegari, A., Lotfi, H., Bonabi, E., and Zarghami, N. (2020). Apoptotic Effect of Saccharomyces cerevisiae on Human Colon Cancer SW480 Cells by Regulation of Akt/NF-ĸB Signaling Pathway. Probiotics Antimicrob. Proteins 12, 311– 319. 10.1007/s12602-019-09528-7.

133. Peng, X., Li, G., and Liu, Z. (2016). Zero-Inflated Beta Regression for Differential Abundance Analysis with Metagenomics Data. J. Comput. Biol. 23, 102–110. 10.1089/cmb.2015.0157.

134. Ma, J., Kim, N., Cha, J.H., Kim, W., Kim, C.Y., Lee, Y., Kim, H.S., Han, Y.D., Yong, D., Han, E., et al. (2026). A human gut metagenome-assembled genome catalogue spanning 41 countries supports genome-scale metabolic models. Nat. Microbiol. 11, 317–334. 10.1038/s41564-025-02206-1.

135. Watson, A.R., Füssel, J., Veseli, I., DeLongchamp, J.Z., Silva, M., Trigodet, F., Lolans, K., Shaiber, A., Fogarty, E., Runde, J.M., et al. (2023). Metabolic independence drives gut microbial colonization and resilience in health and disease. Genome Biol. 24, 78. 10.1186/s13059-023-02924-x.

136. Kanehisa, M., and Goto, S. (2000). KEGG: Kyoto Encyclopedia of Genes and Genomes. Nucleic Acids Res. 28, 27–30. 10.1093/nar/28.1.27.

137. Podlesny, D., Kim, C.Y., Robbani, S.M., Schudoma, C., Fullam, A., Reimer, L.C., Koblitz, J., Schober, I., Iyappan, A., Van Rossum, T., et al. (2026). metaTraits: a large-scale integration of microbial phenotypic trait information. Nucleic Acids Res. 54, D835–D841. 10.1093/nar/gkaf1241.

138. Lind, A.L., McDonald, N.A., Gerrick, E.R., Bhatt, A.S., and Pollard, K.S. (2025). Contiguous and complete assemblies of Blastocystis gut microbiome-associated protists reveal evolutionary diversification to host ecology. Genome Res. 35, 1377–1390. 10.1101/gr.279080.124.

139. Hildebrand, F., Moitinho-Silva, L., Blasche, S., Jahn, M.T., Gossmann, T.I., Huerta-Cepas, J., Hercog, R., Luetge, M., Bahram, M., Pryszlak, A., et al. (2019). Antibiotics-induced monodominance of a novel gut bacterial order. 10.1136/gutjnl-2018-317715.

140. Hildebrand, F., Pallen, M.J., and Bork, P. (2020). Towards standardisation of naming novel prokaryotic taxa in the age of high-throughput microbiology. Gut 69, 1358–1359. 10.1136/gutjnl-2019-319045.

141. Carradec, Q., Pelletier, E., Da Silva, C., Alberti, A., Seeleuthner, Y., Blanc-Mathieu, R., Lima-Mendez, G., Rocha, F., Tirichine, L., Labadie, K., et al. (2018). A global ocean atlas of eukaryotic genes. Nat. Commun. 9, 373. 10.1038/s41467-017-02342-1.

142. Mosca, A., Leclerc, M., and Hugot, J.P. (2016). Gut Microbiota Diversity and Human Diseases: Should We Reintroduce Key Predators in Our Ecosystem? Front. Microbiol. 7. 10.3389/fmicb.2016.00455.

143. Sonnenburg, E.D., and Sonnenburg, J.L. (2019). The ancestral and industrialized gut microbiota and implications for human health. Nat. Rev. Microbiol. 17, 383–390. 10.1038/s41579-019-0191-8.

144. Navarro, S., Ferreira, I., and Loukas, A. (2013). The hookworm pharmacopoeia for inflammatory diseases. Int. J. Parasitol. 43, 225–231. 10.1016/j.ijpara.2012.11.005.

145. Smallwood, T.B., Giacomin, P.R., Loukas, A., Mulvenna, J.P., Clark, R.J., and Miles, J.J. (2017). Helminth Immunomodulation in Autoimmune Disease. Front. Immunol. 8. 10.3389/fimmu.2017.00453.

146. Ianiro, G., Pecere, S., Giorgio, V., Gasbarrini, A., and Cammarota, G. (2016). Digestive Enzyme Supplementation in Gastrointestinal Diseases. Curr. Drug Metab. 17, 187–193. 10.2174/138920021702160114150137.

147. Pokorska, J., Kułaj, D., Dusza, M., Żychlińska-Buczek, J., and Makulska, J. (2016). New Rapid Method of DNA Isolation from Milk Somatic Cells. Anim. Biotechnol. 27, 113–117. 10.1080/10495398.2015.1116446.

148. Sims, C.E., and Allbritton, N.L. (2007). Analysis of single mammalian cells on-chip. Lab. Chip 7, 423–440. 10.1039/b615235j.

149. Gogarten, J.F., Calvignac-Spencer, S., Nunn, C.L., Ulrich, M., Saiepour, N., Nielsen, H.V., Deschner, T., Fichtel, C., Kappeler, P.M., Knauf, S., et al. (2020). Metabarcoding of eukaryotic parasite communities describes diverse parasite assemblages spanning the primate phylogeny. Mol. Ecol. Resour. 20, 204–215. 10.1111/1755-0998.13101.

150. Berlinches de Gea, A., Walochnik, J., Boenigk, J., Dumack, K., Henriquez, F., Rückert, S., Simon, M., and Geisen, S. (2025). Protists as determinants of the One Health framework. ISME J. 19, wraf179. 10.1093/ismejo/wraf179.

151. Castañeda, S., Tomiak, J., Andersen, L.O., Acosta, C.P., Vasquez-A, L.R., Stensvold, C.R., and Ramírez, J.D. (2025). Impact of Blastocystis carriage and colonization intensity on gut microbiota composition in a non-westernized rural population from Colombia. PLoS Negl. Trop. Dis. 19, e0013111. 10.1371/journal.pntd.0013111.

152. Nieves-Ramírez, M.E., Partida-Rodríguez, O., Laforest-Lapointe, I., Reynolds, L.A., Brown, E.M., Valdez-Salazar, A., Morán-Silva, P., Rojas-Velázquez, L., Morien, E., Parfrey, L.W., et al. (2018). Asymptomatic Intestinal Colonization with Protist Blastocystis Is Strongly Associated with Distinct Microbiome Ecological Patterns. mSystems 3, 10.1128/msystems.00007-18. 10.1128/msystems.00007-18.

153. Tito, R.Y., Chaffron, S., Caenepeel, C., Lima-Mendez, G., Wang, J., Vieira-Silva, S., Falony, G., Hildebrand, F., Darzi, Y., Rymenans, L., et al. (2019). Population-level analysis of Blastocystis subtype prevalence and variation in the human gut microbiota. 10.1136/gutjnl-2018-316106.

154. Husnik, F., Tashyreva, D., Boscaro, V., George, E.E., Lukeš, J., and Keeling, P.J. (2021). Bacterial and archaeal symbioses with protists. Curr. Biol. 31, R862–R877. 10.1016/j.cub.2021.05.049.

155. Gast, R.J., Sanders, R.W., and Caron, D.A. (2009). Ecological strategies of protists and their symbiotic relationships with prokaryotic microbes. Trends Microbiol. 17, 563–569. 10.1016/j.tim.2009.09.001.

156. Machado, D., Maistrenko, O.M., Andrejev, S., Kim, Y., Bork, P., Patil, K.R., and Patil, K.R. (2021). Polarization of microbial communities between competitive and cooperative metabolism. Nat. Ecol. Evol., 1–9. 10.1038/s41559-020-01353-4.

157. Maistrenko, O.M., Mende, D.R., Luetge, M., Hildebrand, F., Schmidt, T.S.B., Li, S.S., Rodrigues, J.F.M., von Mering, C., Pedro Coelho, L., Huerta-Cepas, J., et al. (2020). Disentangling the impact of environmental and phylogenetic constraints on prokaryotic within-species diversity. ISME J. 14, 1247– 1259. 10.1038/s41396-020-0600-z.

158. Stenzel, D.J., and Boreham, P.F.L. (1994). Bacteria-like endosymbionts in *Blastocystis* sp. Int. J. Parasitol. 24, 147–149. 10.1016/0020-7519(94)90070-1.

159. Tan, T.C., and Suresh, K.G. (2006). Amoeboid form of Blastocystis hominis—a detailed ultrastructural insight. Parasitol. Res. 99, 737–742. 10.1007/s00436-006-0214-z.

160. Zierdt, C.H., and Tan, H. (1976). Endosymbiosis in *Blastocystis hominis*. Exp. Parasitol. 39, 422– 430. 10.1016/0014-4894(76)90046-1.

161. Shaw, D., Gentekaki, E., and Tsaousis, A.D. The Microbiome Within a Microbe: Rethinking Blastocystis Biology. 10.1111/jeu.70056.

162. Nguyen, T.B.-A., Bonkowski, M., Dumack, K., Chen, Q.-L., He, J.-Z., and Hu, H.-W. (2023). Protistan predation selects for antibiotic resistance in soil bacterial communities. ISME J. 17, 2182–2189. 10.1038/s41396-023-01524-8.

163. Sharon, I., Quijada, N.M., Pasolli, E., Fabbrini, M., Vitali, F., Agamennone, V., Dötsch, A., Selberherr, E., Grau, J.H., Meixner, M., et al. (2022). The Core Human Microbiome: Does It Exist and How Can We Find It? A Critical Review of the Concept. Nutrients 14, 2872. 10.3390/nu14142872.

164. Antunes, P., Novais, C., and Peixe, L. (2020). Food-to-Humans Bacterial Transmission. Microbiol. Spectr. 8, 10.1128/microbiolspec.mtbp-0019–2016. 10.1128/microbiolspec.mtbp-0019-2016.

165. Ratnasingham, S., and Hebert, P.D.N. (2007). bold: The Barcode of Life Data System (http://www.barcodinglife.org). Mol. Ecol. Notes 7, 355–364. 10.1111/j.1471-8286.2007.01678.x.

166. Soppa, J. (2017). Polyploidy and community structure. Nat. Microbiol. 2, 16261. 10.1038/nmicrobiol.2016.261.

167. Van de Peer, Y., Mizrachi, E., and Marchal, K. (2017). The evolutionary significance of polyploidy. Nat. Rev. Genet. 18, 411–424. 10.1038/nrg.2017.26.

168. Prasoodanan PK, V., Maistrenko, O.M., Fullam, A., Mende, D.R., Kartal, E., Coelho, L.P., Spang, A., Bork, P., and Schmidt, T.S.B. (2026). Unbinned contigs expand known diversity in the global microbiome. Nat. Microbiol., 1–13. 10.1038/s41564-026-02314-6.

169. Li, H., and Durbin, R. (2009). Fast and accurate short read alignment with Burrows–Wheeler transform. Bioinformatics 25, 1754–1760. 10.1093/bioinformatics/btp324.

170. Arranz, V., Pearman, W.S., Aguirre, J.D., and Liggins, L. (2020). MARES, a replicable pipeline and curated reference database for marine eukaryote metabarcoding. Sci. Data 7, 209. 10.1038/s41597-020-0549-9.

171. Jochum, L., and Stecher, B. (2020). Label or Concept – What Is a Pathobiont? Trends Microbiol. 10.1016/j.tim.2020.04.011.

172. WHO fungal priority pathogens list to guide research, development and public health action https://www.who.int/publications/i/item/9789240060241.

173. CDC (2025). A-Z Index of Parasitic Diseases. Parasites. https://www.cdc.gov/parasites/listing/index.html.

174. Prevention and control of intestinal parasitic infections: WHO Technical Report Series N°749 https://www.who.int/publications/i/item/WHO-TRS-749.

175. Tomofuji, Y., Sonehara, K., Kishikawa, T., Maeda, Y., Ogawa, K., Kawabata, S., Nii, T., Okuno, T., Oguro-Igashira, E., Kinoshita, M., et al. (2023). Reconstruction of the personal information from human genome reads in gut metagenome sequencing data. Nat. Microbiol., 1–16. 10.1038/s41564-023-01381-3.

176. Parks, D.H., Chuvochina, M., Rinke, C., Mussig, A.J., Chaumeil, P.-A., and Hugenholtz, P. (2022). GTDB: an ongoing census of bacterial and archaeal diversity through a phylogenetically consistent, rank normalized and complete genome-based taxonomy. Nucleic Acids Res. 50, D785–D794. 10.1093/nar/gkab776.

177. Chaumeil, P.-A., Mussig, A.J., Hugenholtz, P., and Parks, D.H. (2022). GTDB-Tk v2: memory friendly classification with the genome taxonomy database. Bioinformatics 38, 5315–5316. 10.1093/bioinformatics/btac672.

178. Li, H. (2018). Minimap2: pairwise alignment for nucleotide sequences. Bioinformatics 34, 3094– 3100. 10.1093/bioinformatics/bty191.

179. Langmead, B., and Salzberg, S.L. (2012). Fast gapped-read alignment with Bowtie 2. Nat. Methods 9, 357–359. 10.1038/nmeth.1923.

180. Aramaki, T., Blanc-Mathieu, R., Endo, H., Ohkubo, K., Kanehisa, M., Goto, S., and Ogata, H. (2020). KofamKOALA: KEGG Ortholog assignment based on profile HMM and adaptive score threshold. Bioinformatics 36, 2251–2252. 10.1093/bioinformatics/btz859.

181. Palù, M., Basile, A., Zampieri, G., Treu, L., Rossi, A., Morlino, M.S., and Campanaro, S. (2022). KEMET – A python tool for KEGG Module evaluation and microbial genome annotation expansion. Comput. Struct. Biotechnol. J. 20, 1481–1486. 10.1016/j.csbj.2022.03.015.

182. Pečar, J., Rupert, L., and Wahlers, M. (2020). EMBL Heidelberg HPC Cluster. 10.5281/zenodo.12785830.

183. Keesey, T.M. PhyloPic. https://www.phylopic.org.

