## Supplementary material for "Identification of eukaryotes as a bycatch in global microbiome data": Figure S1

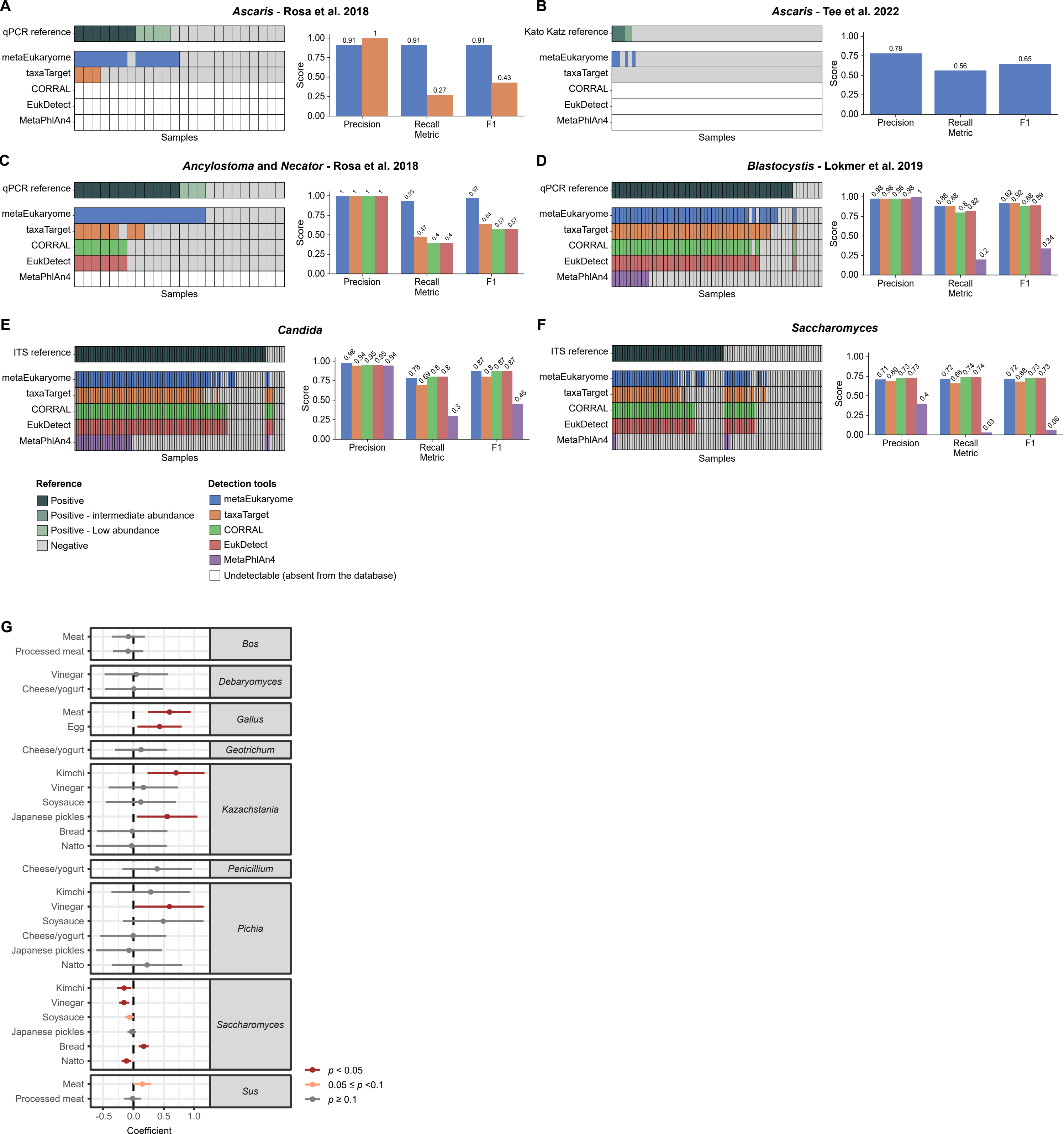

A
Ascaris - Rosa et al. 2018
qPCR reference
metaEukaryome
taxaTarget
CORRAL
EukDetect
MetaPhlAn4
Samples
Score
Precision
Recall
F1
Metric
0.91
1
0.91
0.27
0.91
0.43
B
Ascaris - Tee et al. 2022
Kato Katz reference
0.78
0.56
0.65
C
Ancylostoma and Necator - Rosa et al. 2018
0.93
0.47
0.4
0.4
0.97
0.64
0.57
0.57
D
Blastocystis - Lokmer et al. 2019
0.98
0.88
0.8
0.82
0.2
0.92
0.89
0.34
E
Candida
ITS reference
0.94
0.95
0.78
0.69
0.3
0.87
0.45
F
Saccharomyces
0.71
0.69
0.73
0.4
0.72
0.66
0.74
0.03
0.68
0.06
Reference
Positive
Positive - intermediate abundance
Positive - Low abundance
Negative
Detection tools
metaEukaryome
taxaTarget
CORRAL
EukDetect
MetaPhlAn4
Undetectable (absent from the database)
G
Meat
Processed meat
Bos
Vinegar
Cheese/yogurt
Debaryomyces
Egg
Gallus
Geotrichum
Kimchi
Soysauce
Japanese pickles
Bread
Natto
Kazachstania
Penicillium
Pichia
Saccharomyces
Sus
Coefficient
p < 0.05
0.05 ≤ p <0.1
p ≥ 0.1
