## Supplementary material for "Identification of eukaryotes as a bycatch in global microbiome data": Figure S2

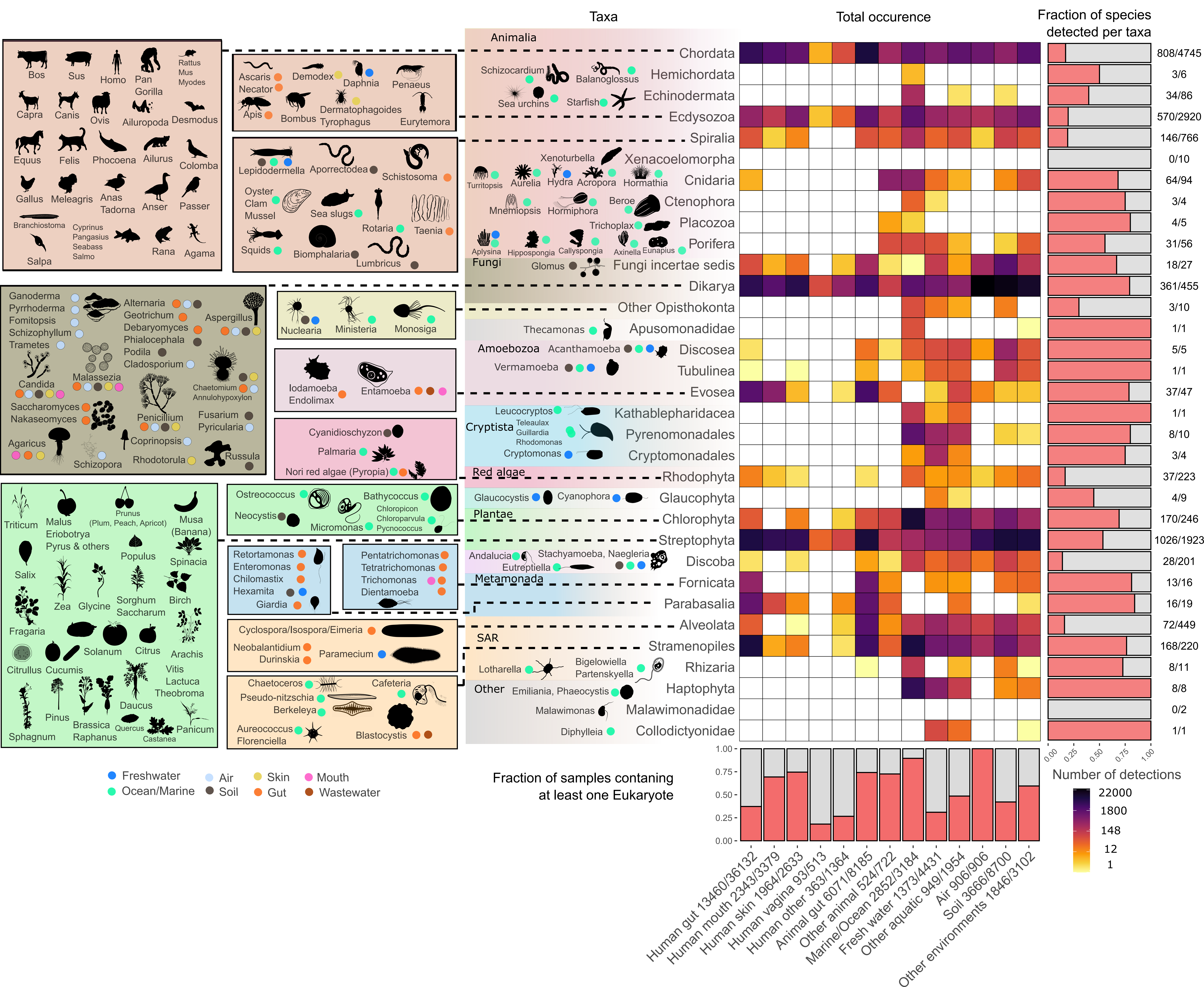

Taxa
Total occurence
Fraction of species detected per taxa
Bos
Sus
Homo
Pan
Gorilla
Rattus
Mus
Myodes
Capra
Canis
Ovis
Ailuropoda
Desmodus
Equus
Felis
Phocoena
Ailurus
Colomba
Gallus
Meleagris
Anas
Tadorna
Anser
Passer
Branchiostoma
Cyprinus
Pangasius
Seabass
Salmo
Rana
Agama
Salpa
Ascaris
Necator
Demodex
Daphnia
Penaeus
Apis
Bombus
Dermatophagoides
Tyrophagus
Eurytemora
Lepidodermella
Aporrectodea
Schistosoma
Oyster
Clam
Mussel
Sea slugs
Rotaria
Taenia
Squids
Biomphalaria
Lumbricus
Animalia
Chordata
Schizocardium
Balanoglossus
Hemichordata
Sea urchins
Starfish
Echinodermata
Ecdysozoa
Spiralia
Xenoturbella
Xenacoelomorpha
Turritopsis
Aurelia
Hydra
Acropora
Hormathia
Cnidaria
Mnemiopsis
Hormiphora
Beroe
Ctenophora
Trichoplax
Placozoa
Aplysina
Hippospongia
Callyspongia
Axinella
Eunapius
Porifera
Fungi
Glomus
Fungi incertae sedis
Dikarya
Ganoderma
Pyrrhoderma
Fomitopsis
Schizophyllum
Trametes
Alternaria
Geotrichum
Debaryomyces
Phialocephala
Podila
Cladosporium
Aspergillus
Candida
Malassezia
Chaetomium
Annulohypoxylon
Saccharomyces
Nakaseomyces
Penicillium
Fusarium
Pyricularia
Agaricus
Coprinopsis
Schizopora
Rhodotorula
Russula
Nuclearia
Ministeria
Monosiga
Other Opisthokonta
Thecamonas
Apusomonadidae
Amoebozoa
Acanthamoeba
Discosea
Vermamoeba
Tubulinea
Iodamoeba
Endolimax
Entamoeba
Evosea
Leucocryptos
Kathablepharidacea
Cryptista
Teleaulax
Guillardia
Rhodomonas
Pyrenomonadales
Cryptomonas
Cryptomonadales
Cyanidioschyzon
Palmaria
Nori red algae (Pyropia)
Red algae
Rhodophyta
Glaucocystis
Cyanophora
Glaucophyta
Plantae
Chlorophyta
Streptophyta
Ostreococcus
Bathycoccus
Neocystis
Chloropicon
Chloroparvula
Micromonas
Pycnococcus
Triticum
Malus
Eriobotrya
Pyrus & others
Prunus
(Plum, Peach, Apricot)
Musa
(Banana)
Populus
Spinacia
Salix
Zea
Glycine
Sorghum
Saccharum
Birch
Fragaria
Solanum
Citrus
Arachis
Citrullus
Cucumis
Vitis
Lactuca
Theobroma
Pinus
Daucus
Brassica
Raphanus
Quercus
Castanea
Panicum
Sphagnum
Andalucia
Eutreptiella
Stachyameba, Naegleria
Discoba
Metamonada
Fornicata
Parabasalia
Retortamonas
Enteromonas
Chilomastix
Hexamita
Giardia
Pentatrichomonas
Tetratrichomonas
Trichomonas
Dientamoeba
Alveolata
SAR
Stramenopiles
Cyclospora/Isospora/Eimeria
Neobalantidium
Durinskia
Paramecium
Lotharella
Bigelowiella
Partenskyella
Rhizaria
Other
Emiliania, Phaeocystis
Haptophyta
Malawimonas
Malawimonadidae
Diphylleia
Collodictyonidae
Chaetoceros
Pseudo-nitzschia
Berkeleya
Cafeteria
Aureococcus
Florenciella
Blastocystis
808/4745
3/6
34/86
570/2920
146/766
0/10
64/94
3/4
4/5
31/56
18/27
361/455
3/10
1/1
5/5
1/1
37/47
1/1
8/10
3/4
37/223
4/9
170/246
1026/1923
28/201
13/16
16/19
72/449
168/220
8/11
8/8
0/2
1/1
0.00
0.25
0.50
0.75
1.00
Freshwater
Ocean/Marine
Air
Soil
Skin
Gut
Mouth
Wastewater
Fraction of samples contaning at least one Eukaryote
Number of detections
22000
1800
148
12
1
Human gut 13460/36132
Human mouth 2343/3379
Human skin 1964/2633
Human vagina 93/513
Human other 363/1364
Animal gut 6071/8185
Other animal 524/722
Marine/Ocean 2852/3184
Fresh water 1373/4431
Other aquatic 949/1954
Air 906/906
Soil 3666/8700
Other environments 1846/3102
