## Supplementary figures and images for "Identification of eukaryotes as a bycatch in global microbiome data"

### Figure S3

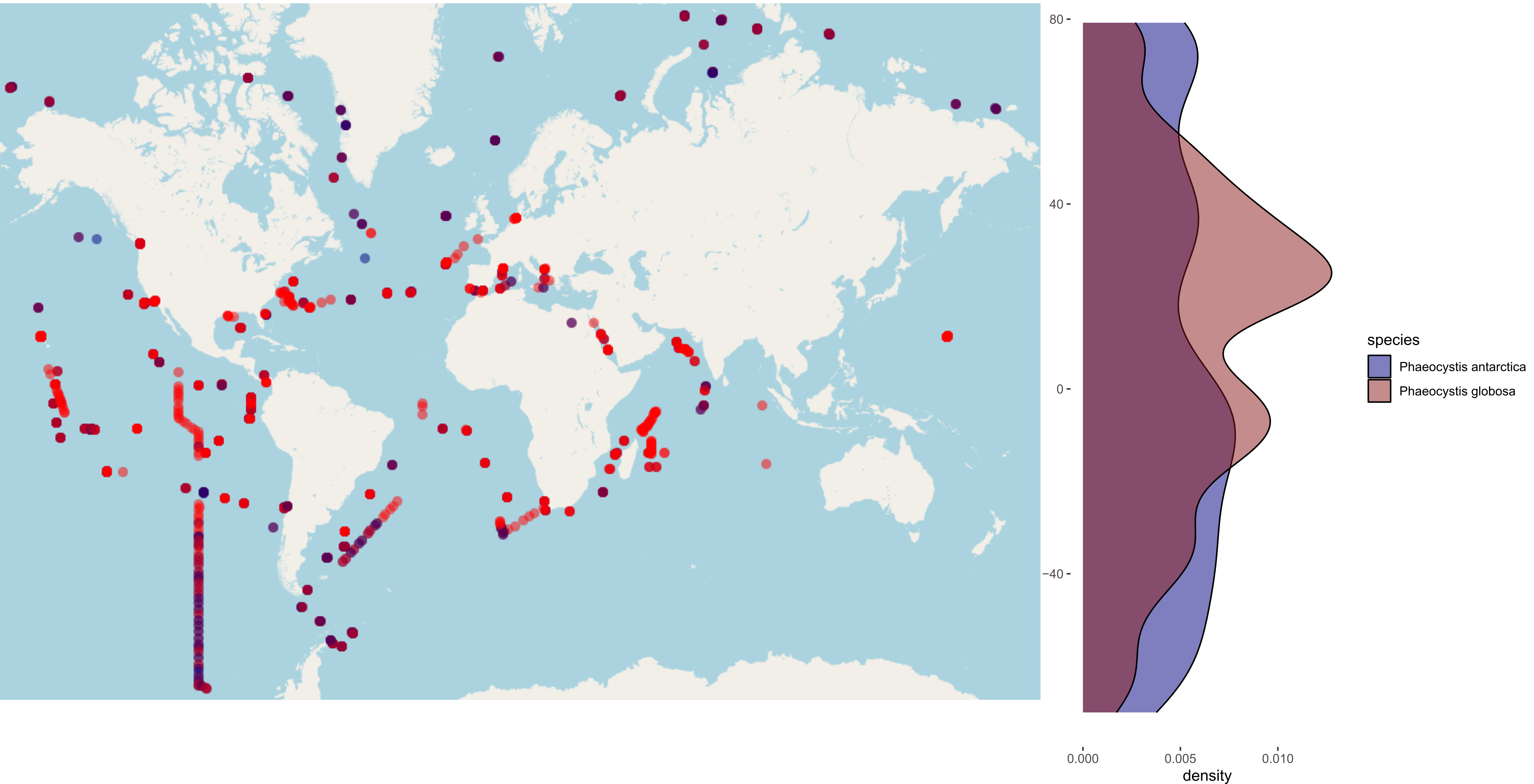

80
40
0
−40
0.000
0.005
0.010
density
species
Phaeocystis antarctica
Phaeocystis globosa

### Figure S12

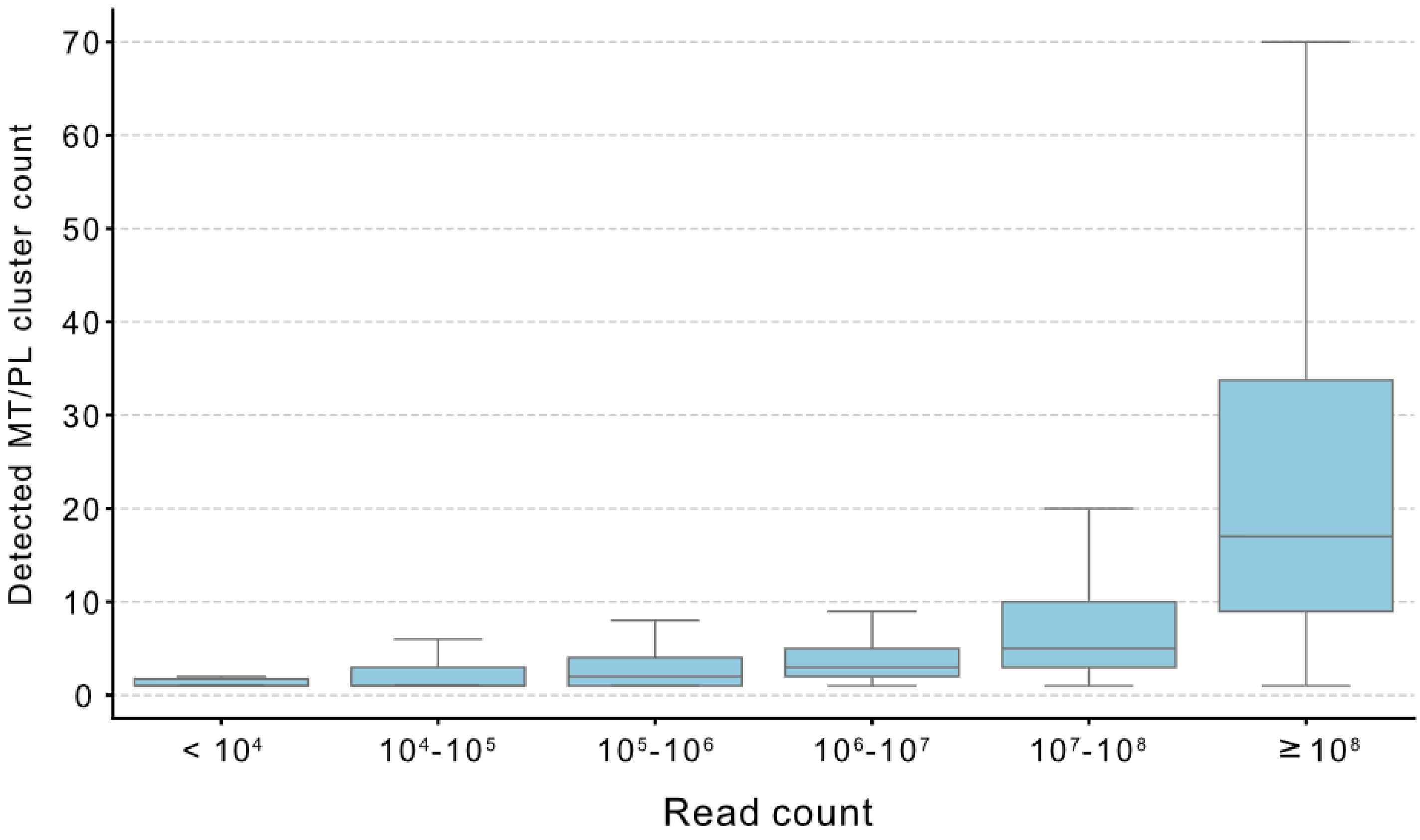

Detected MT/PL cluster count
70
60
50
40
30
20
10
0
< 10⁴
10⁴-10⁵
10⁵-10⁶
10⁶-10⁷
10⁷-10⁸
≥ 10⁸
Read count
