## Supplementary material for "Identification of eukaryotes as a bycatch in global microbiome data": Figure S4

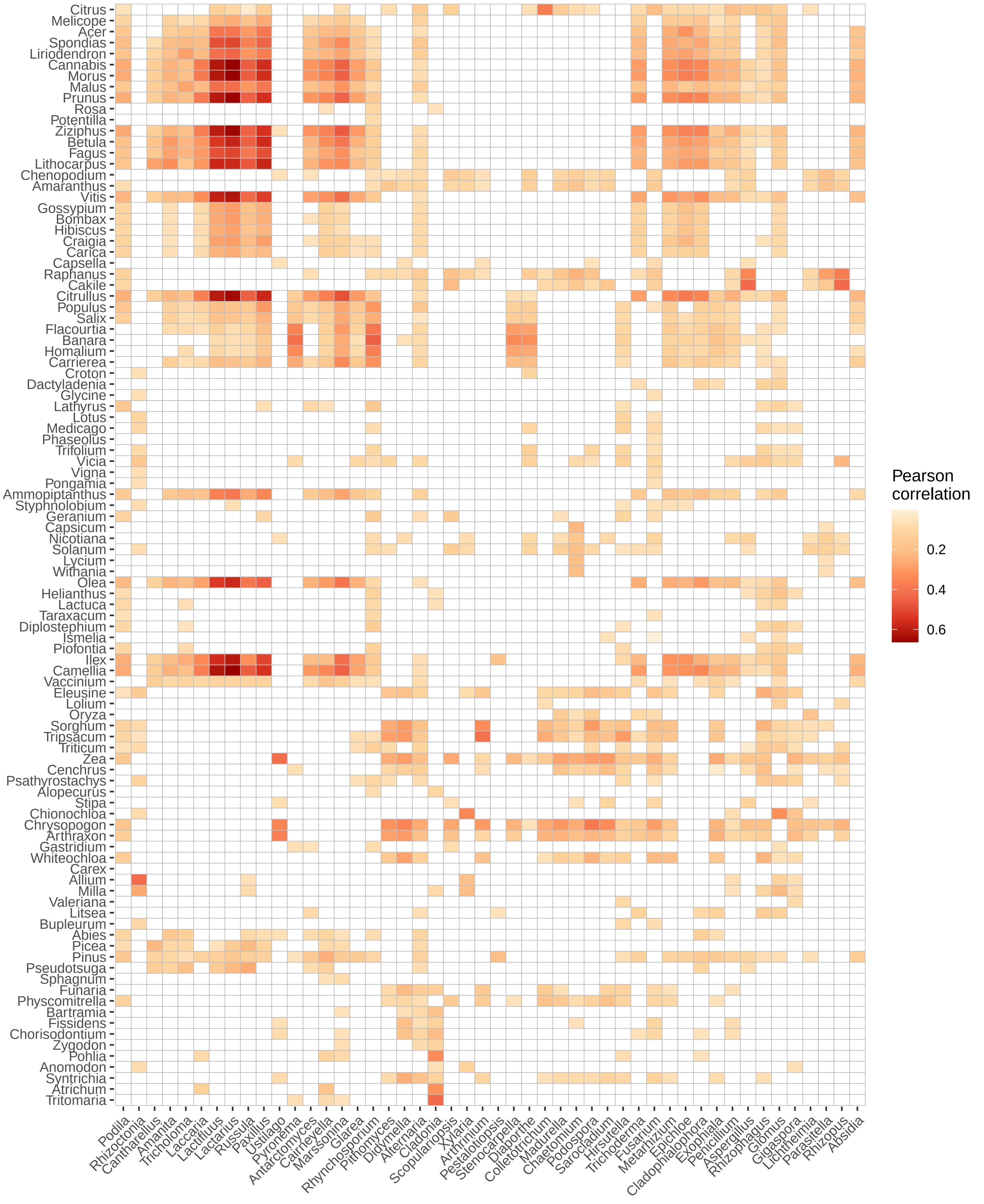

Citrus
Melicope
Acer
Spondias
Liriodendron
Cannabis
Morus
Malus
Prunus
Rosa
Potentilla
Ziziphus
Betula
Fagus
Lithocarpus
Chenopodium
Amaranthus
Vitis
Gossypium
Bombax
Hibiscus
Craigia
Carica
Capsella
Raphanus
Cakile
Citrullus
Populus
Salix
Flacourtia
Banara
Homalium
Carrierea
Croton
Dactyladenia
Glycine
Lathyrus
Lotus
Medicago
Phaseolus
Trifolium
Vicia
Vigna
Pongamia
Ammopiptanthus
Styphnolobium
Geranium
Capsicum
Nicotiana
Solanum
Lycium
Withania
Olea
Helianthus
Lactuca
Taraxacum
Diplostephium
Ismelia
Piofontia
Ilex
Camellia
Vaccinium
Eleusine
Lolium
Oryza
Sorghum
Tripsacum
Triticum
Zea
Cenchrus
Psathyrostachys
Alopecurus
Stipa
Chionochloa
Chrysopogon
Arthraxon
Gastridium
Whiteochloa
Carex
Allium
Milla
Valeriana
Litsea
Bupleurum
Abies
Picea
Pinus
Pseudotsuga
Sphagnum
Funaria
Physcomitrella
Bartramia
Fissidens
Chorisodontium
Zygodon
Pohlia
Anomodon
Syntrichia
Atrichum
Tritomaria
Podila
Rhizoctonia
Cantharellus
Amanita
Tricholoma
Laccaria
Lactifluus
Lactarius
Russula
Paxillus
Ustilago
Pyronema
Antarctomyces
Cairneyella
Marssonina
Glarea
Rhynchosporium
Pithomyces
Didymella
Alternaria
Cladonia
Scopulariopsis
Xylaria
Arthrinium
Pestalotiopsis
Stenocarpella
Diaporthe
Colletotrichum
Madurella
Chaetomium
Podospora
Sarocladium
Hirsutella
Trichoderma
Fusarium
Metarhizium
Epichloe
Cladophialophora
Exophiala
Penicillium
Aspergillus
Rhizophagus
Glomus
Gigaspora
Lichtheimia
Parasitella
Rhizopus
Absidia
Pearson correlation
0.2
0.4
0.6
