## Supplementary material for "Identification of eukaryotes as a bycatch in global microbiome data": Figure S5

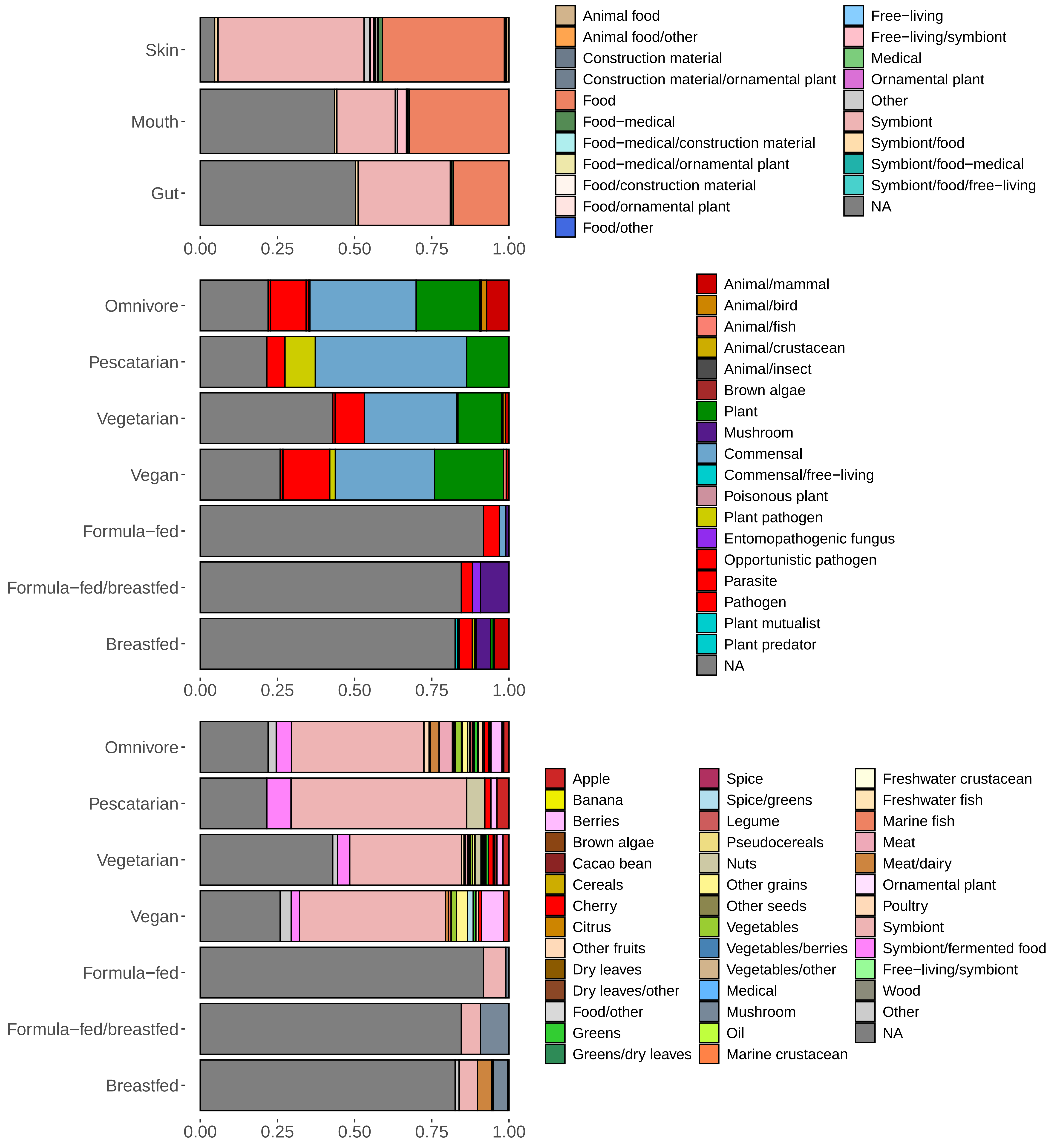
Skin
Mouth
Gut
0.00
0.25
0.50
0.75
1.00
Animal food
Animal food/other
Construction material
Construction material/ornamental plant
Food
Food−medical
Food−medical/construction material
Food−medical/ornamental plant
Food/construction material
Food/ornamental plant
Food/other
Free−living
Free−living/symbiont
Medical
Ornamental plant
Other
Symbiont
Symbiont/food
Symbiont/food−medical
Symbiont/food/free−living
NA
Omnivore
Pescatarian
Vegetarian
Vegan
Formula−fed
Formula−fed/breastfed
Breastfed
0.00
0.25
0.50
0.75
1.00
Animal/mammal
Animal/bird
Animal/fish
Animal/crustacean
Animal/insect
Brown algae
Plant
Mushroom
Commensal
Commensal/free−living
Poisonous plant
Plant pathogen
Entomopathogenic fungus
Opportunistic pathogen
Parasite
Pathogen
Plant mutualist
Plant predator
NA
Omnivore
Pescatarian
Vegetarian
Vegan
Formula−fed
Formula−fed/breastfed
Breastfed
0.00
0.25
0.50
0.75
1.00
Apple
Banana
Berries
Brown algae
Cacao bean
Cereals
Cherry
Citrus
Other fruits
Dry leaves
Dry leaves/other
Food/other
Greens
Greens/dry leaves
Spice
Spice/greens
Legume
Pseudocereals
Nuts
Other grains
Other seeds
Vegetables
Vegetables/berries
Vegetables/other
Medical
Mushroom
Oil
Marine crustacean
Freshwater crustacean
Freshwater fish
Marine fish
Meat
Meat/dairy
Ornamental plant
Poultry
Symbiont
Symbiont/fermented food
Free−living/symbiont
Wood
Other
NA
