## Supplementary material for "Identification of eukaryotes as a bycatch in global microbiome data": Figure S6

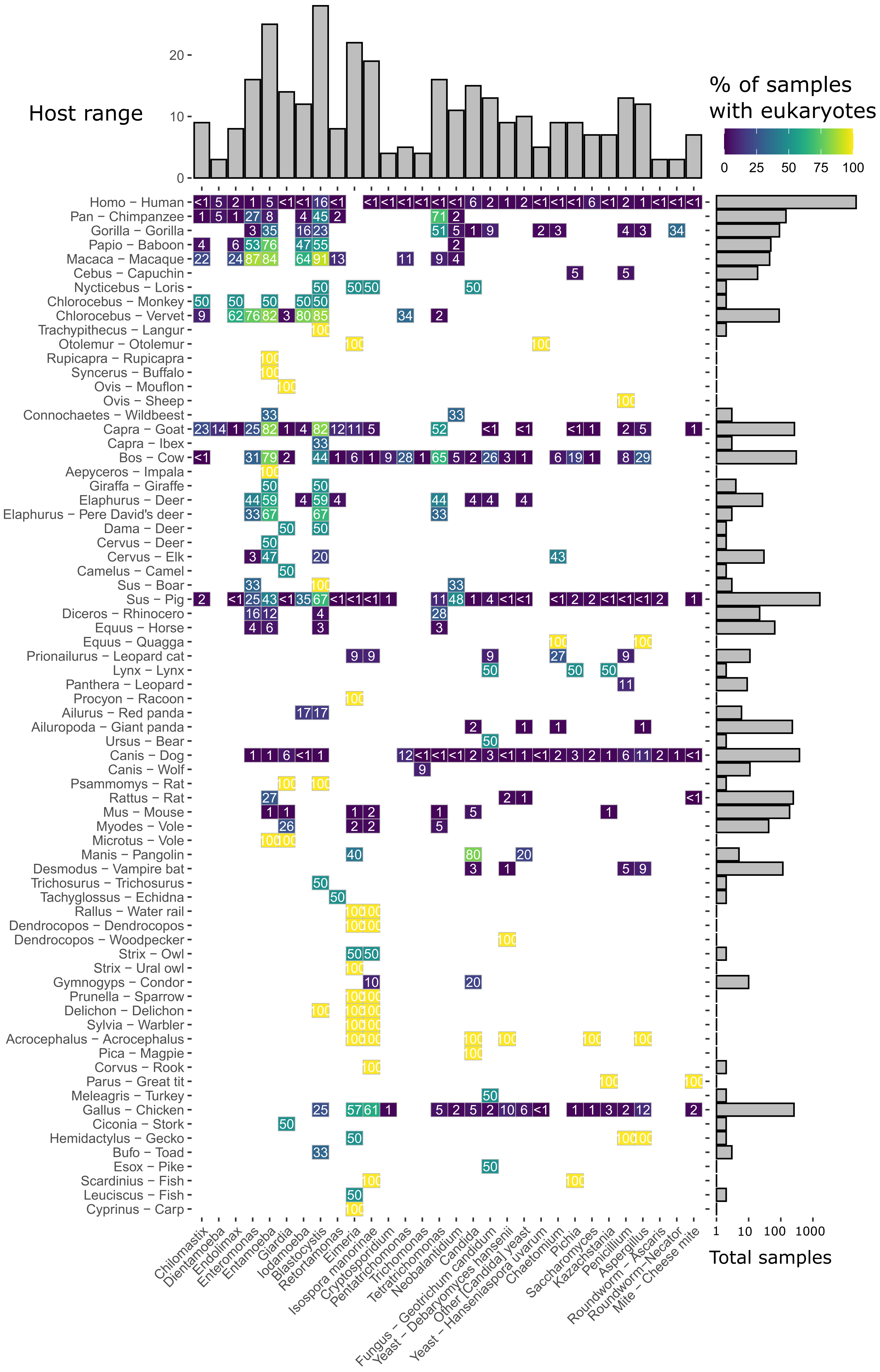

Host range
% of samples
with eukaryotes
0
25
50
75
100
0
10
20
Total samples
1
10
100
1000
Homo − Human
Pan − Chimpanzee
Gorilla − Gorilla
Papio − Baboon
Macaca − Macaque
Cebus − Capuchin
Nycticebus − Loris
Chlorocebus − Monkey
Chlorocebus − Vervet
Trachypithecus − Langur
Otolemur − Otolemur
Rupicapra − Rupicapra
Syncerus − Buffalo
Ovis − Mouflon
Ovis − Sheep
Connochaetes − Wildbeest
Capra − Goat
Capra − Ibex
Bos − Cow
Aepyceros − Impala
Giraffa − Giraffe
Elaphurus − Deer
Elaphurus − Pere David's deer
Dama − Deer
Cervus − Deer
Cervus − Elk
Camelus − Camel
Sus − Boar
Sus − Pig
Diceros − Rhinocero
Equus − Horse
Equus − Quagga
Prionailurus − Leopard cat
Lynx − Lynx
Panthera − Leopard
Procyon − Racoon
Ailurus − Red panda
Ailuropoda − Giant panda
Ursus − Bear
Canis − Dog
Canis − Wolf
Psammomys − Rat
Rattus − Rat
Mus − Mouse
Myodes − Vole
Microtus − Vole
Manis − Pangolin
Desmodus − Vampire bat
Trichosurus − Trichosurus
Tachyglossus − Echidna
Rallus − Water rail
Dendrocopos − Dendrocopos
Dendrocopos − Woodpecker
Strix − Owl
Strix − Ural owl
Gymnogyps − Condor
Prunella − Sparrow
Delichon − Delichon
Sylvia − Warbler
Acrocephalus − Acrocephalus
Pica − Magpie
Corvus − Rook
Parus − Great tit
Meleagris − Turkey
Gallus − Chicken
Ciconia − Stork
Hemidactylus − Gecko
Bufo − Toad
Esox − Pike
Scardinius − Fish
Leuciscus − Fish
Cyprinus − Carp
Chilomastix
Dientamoeba
Endolimax
Enteromonas
Entamoeba
Giardia
Iodamoeba
Blastocystis
Retortamonas
Eimeria
Isospora manorinae
Cryptosporidium
Pentatrichomonas
Trichomonas
Tetratrichomonas
Neobalantidium
Candida
Fungus − Geotrichum candidum
Yeast − Debaryomyces hansenii
Other [Candida] yeast
Yeast − Hanseniaspora uvarum
Chaetomium
Pichia
Saccharomyces
Kazachstania
Penicillium
Aspergillus
Roundworm − Ascaris
Roundworm−Necator
Mite − Cheese mite
