## Supplementary material for "Identification of eukaryotes as a bycatch in global microbiome data": Figure S7

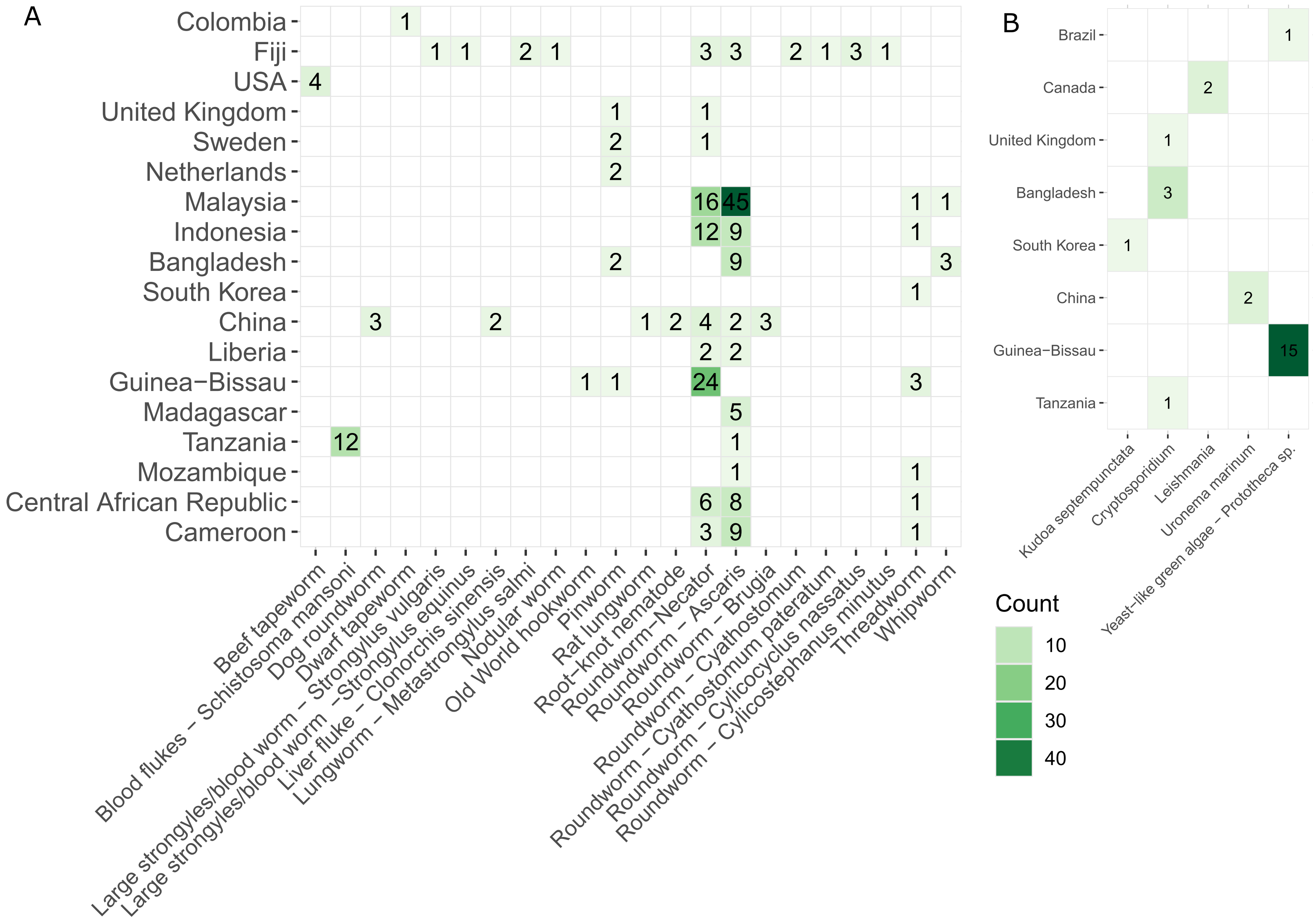
A
Colombia
Fiji
USA
United Kingdom
Sweden
Netherlands
Malaysia
Indonesia
Bangladesh
South Korea
China
Liberia
Guinea−Bissau
Madagascar
Tanzania
Mozambique
Central African Republic
Cameroon
Beef tapeworm
Blood flukes – Schistosoma mansoni
Dog roundworm
Dwarf tapeworm
Large strongyles/blood worm – Strongylus vulgaris
Large strongyles/blood worm –Strongylus equinus
Liver fluke – Clonorchis sinensis
Lungworm – Metastrongylus salmi
Nodular worm
Old World hookworm
Pinworm
Rat lungworm
Root−knot nematode
Roundworm−Necator
Roundworm – Ascaris
Roundworm – Brugia
Roundworm – Cyathostomum
Roundworm – Cyathostomum pateratum
Roundworm – Cylicocyclus nassatus
Roundworm – Cylicostephanus minutus
Threadworm
Whipworm
B
Brazil
Canada
United Kingdom
Bangladesh
South Korea
China
Guinea−Bissau
Tanzania
Kudoa septempunctata
Cryptosporidium
Leishmania
Uronema marinum
Yeast−like green algae – Prototheca sp.
Count
10
20
30
40
