## Supplementary material for "Identification of eukaryotes as a bycatch in global microbiome data": Figure S8

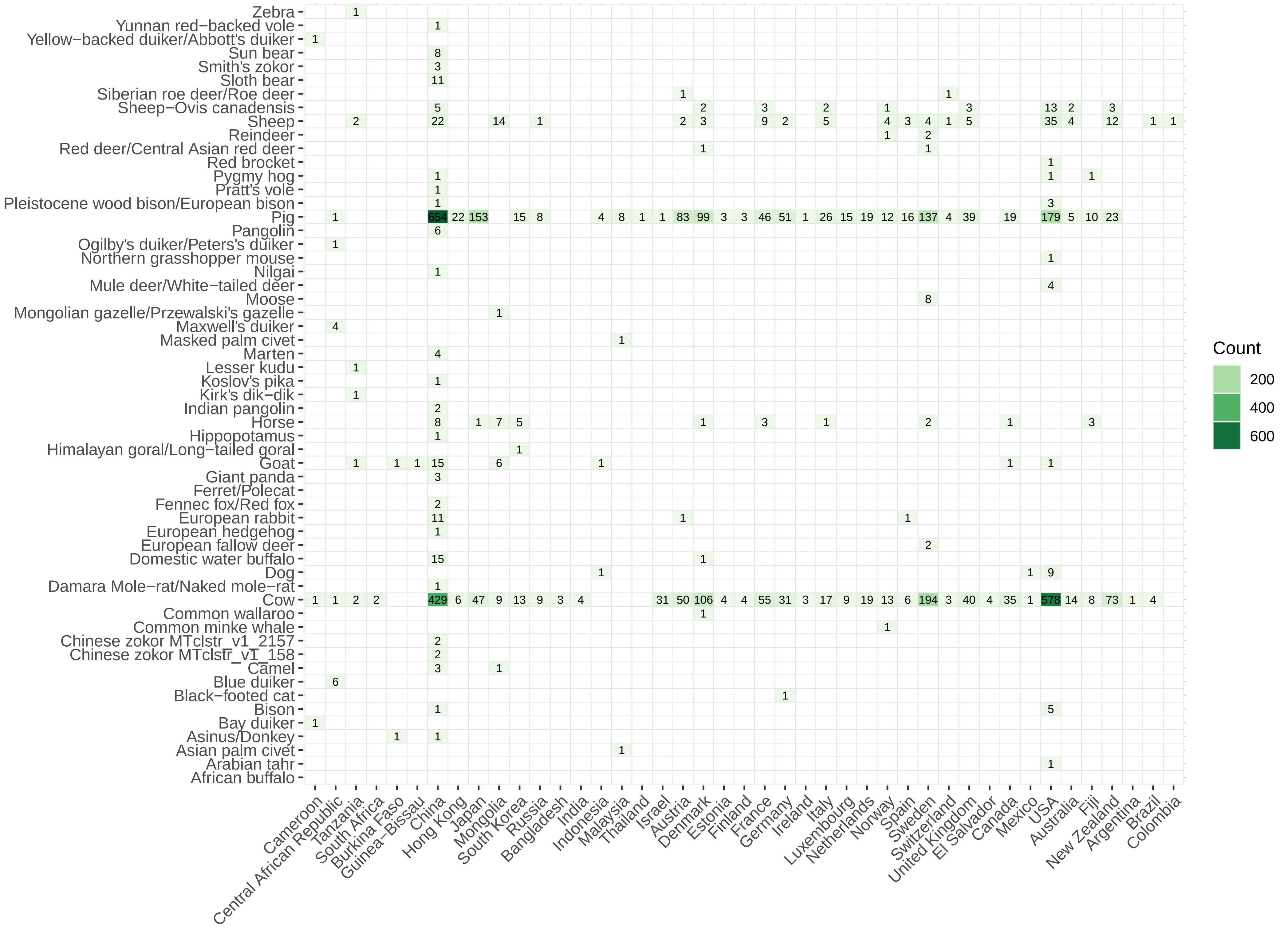
Zebra
Yunnan red−backed vole
Yellow−backed duiker/Abbott's duiker
Sun bear
Smith's zokor
Sloth bear
Siberian roe deer/Roe deer
Sheep−Ovis canadensis
Sheep
Reindeer
Red deer/Central Asian red deer
Red brocket
Pygmy hog
Pratt's vole
Pleistocene wood bison/European bison
Pig
Pangolin
Ogilby's duiker/Peters's duiker
Northern grasshopper mouse
Nilgai
Mule deer/White−tailed deer
Moose
Mongolian gazelle/Przewalski's gazelle
Maxwell's duiker
Masked palm civet
Marten
Lesser kudu
Koslov's pika
Kirk's dik−dik
Indian pangolin
Horse
Hippopotamus
Himalayan goral/Long−tailed goral
Goat
Giant panda
Ferret/Polecat
Fennec fox/Red fox
European rabbit
European hedgehog
European fallow deer
Domestic water buffalo
Dog
Damara Mole−rat/Naked mole−rat
Cow
Common wallaroo
Common minke whale
Chinese zokor MTclstr_v1_2157
Chinese zokor MTclstr_v1_158
Camel
Blue duiker
Black−footed cat
Bison
Bay duiker
Asinus/Donkey
Asian palm civet
Arabian tahr
African buffalo
Cameroon
Central African Republic
Tanzania
South Africa
Burkina Faso
Guinea−Bissau
China
Hong Kong
Japan
Mongolia
South Korea
Russia
Bangladesh
India
Indonesia
Malaysia
Thailand
Israel
Austria
Denmark
Estonia
Finland
France
Germany
Ireland
Italy
Luxembourg
Netherlands
Norway
Spain
Sweden
Switzerland
United Kingdom
El Salvador
Canada
Mexico
USA
Australia
Fiji
New Zealand
Argentina
Brazil
Colombia
Count
200
400
600
