## Supplementary material for "Identification of eukaryotes as a bycatch in global microbiome data": Figure S9

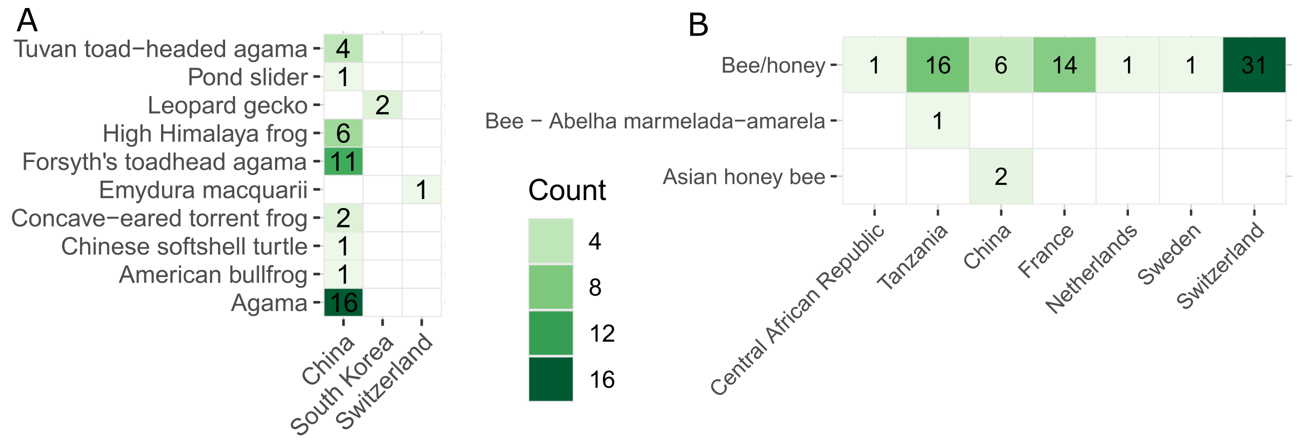
A
Tuvan toad−headed agama
Pond slider
Leopard gecko
High Himalaya frog
Forsyth's toadhead agama
Emydura macquarii
Concave−eared torrent frog
Chinese softshell turtle
American bullfrog
Agama
4
1
2
6
11
1
2
1
1
16
China
South Korea
Switzerland
Count
4
8
12
16
B
Bee/honey
Bee − Abelha marmelada−amarela
Asian honey bee
1
16
6
14
1
1
31
1
2
Central African Republic
Tanzania
China
France
Netherlands
Sweden
Switzerland
