## Supplementary material for "Identification of eukaryotes as a bycatch in global microbiome data": Figure S10

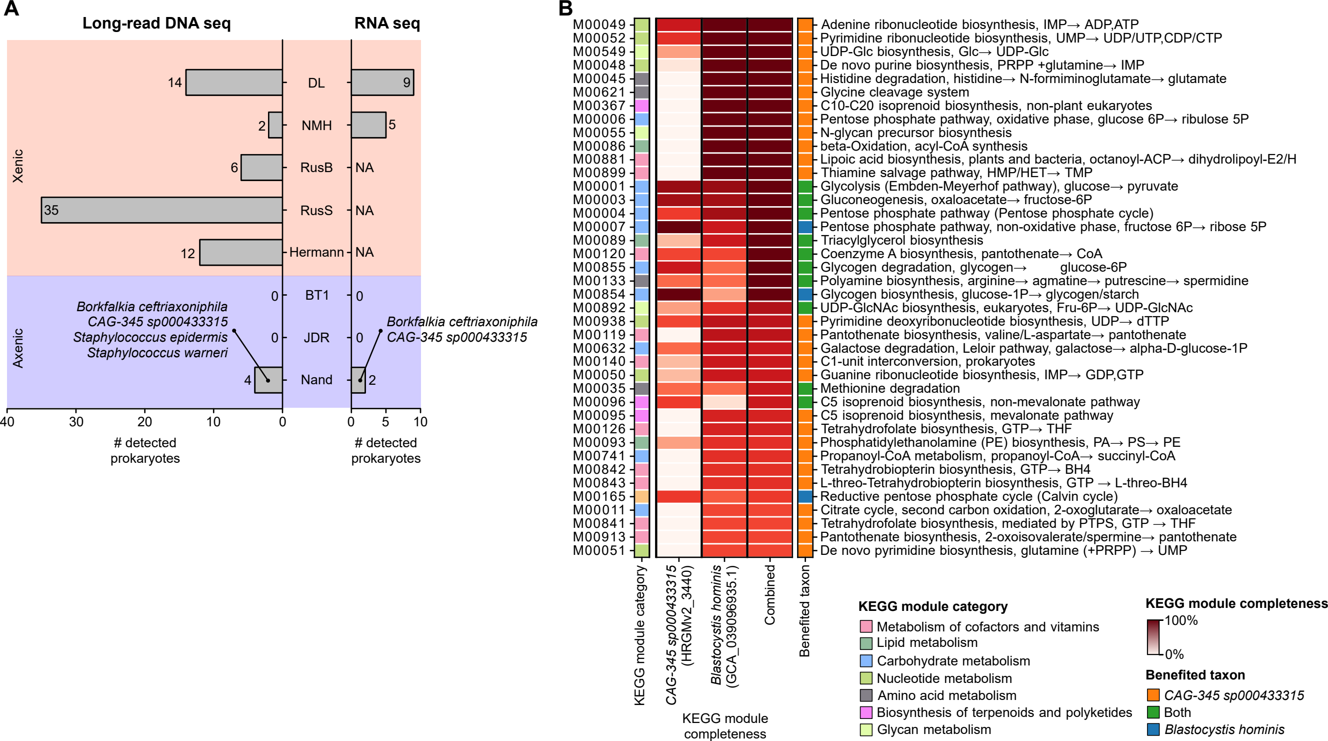

A
Long-read DNA seq
RNA seq
Xenic
Axenic
14
DL
9
2
NMH
5
6
RusB
NA
35
RusS
NA
12
Hermann
NA
0
BT1
0
0
JDR
0
4
Nand
2
Borkfalkia ceftriaxoniphila
CAG-345 sp000433315
Staphylococcus epidermis
Staphylococcus warneri
Borkfalkia ceftriaxoniphila
CAG-345 sp000433315
40 30 20 10 0
0 5 10
### detected prokaryotes
### detected prokaryotes
B
M00049 Adenine ribonucleotide biosynthesis, IMP→ ADP,ATP
M00052 Pyrimidine ribonucleotide biosynthesis, UMP→ UDP/UTP,CDP/CTP
M00549 UDP-Glc biosynthesis, Glc→ UDP-Glc
M00048 De novo purine biosynthesis, PRPP +glutamine→ IMP
M00045 Histidine degradation, histidine→ N-formiminoglutamate→ glutamate
M00621 Glycine cleavage system
M00367 C10-C20 isoprenoid biosynthesis, non-plant eukaryotes
M00006 Pentose phosphate pathway, oxidative phase, glucose 6P→ ribulose 5P
M00055 N-glycan precursor biosynthesis
M00086 beta-Oxidation, acyl-CoA synthesis
M00881 Lipoic acid biosynthesis, plants and bacteria, octanoyl-ACP→ dihydrolipoyl-E2/H
M00899 Thiamine salvage pathway, HMP/HET→ TMP
M00001 Glycolysis (Embden-Meyerhof pathway), glucose→ pyruvate
M00003 Gluconeogenesis, oxaloacetate→ fructose-6P
M00004 Pentose phosphate pathway (Pentose phosphate cycle)
M00007 Pentose phosphate pathway, non-oxidative phase, fructose 6P→ ribose 5P
M00089 Triacylglycerol biosynthesis
M00120 Coenzyme A biosynthesis, pantothenate→ CoA
M00855 Glycogen degradation, glycogen→ glucose-6P
M00133 Polyamine biosynthesis, arginine→ agmatine→ putrescine→ spermidine
M00854 Glycogen biosynthesis, glucose-1P→ glycogen/starch
M00892 UDP-GlcNAc biosynthesis, eukaryotes, Fru-6P→ UDP-GlcNAc
M00938 Pyrimidine deoxyribonucleotide biosynthesis, UDP→ dTTP
M00119 Pantothenate biosynthesis, valine/L-aspartate→ pantothenate
M00632 Galactose degradation, Leloir pathway, galactose→ alpha-D-glucose-1P
M00140 C1-unit interconversion, prokaryotes
M00050 Guanine ribonucleotide biosynthesis, IMP→ GDP,GTP
M00035 Methionine degradation
M00096 C5 isoprenoid biosynthesis, non-mevalonate pathway
M00095 C5 isoprenoid biosynthesis, mevalonate pathway
M00126 Tetrahydrofolate biosynthesis, GTP→ THF
M00093 Phosphatidylethanolamine (PE) biosynthesis, PA→ PS→ PE
M00741 Propanoyl-CoA metabolism, propanoyl-CoA→ succinyl-CoA
M00842 Tetrahydrobiopterin biosynthesis, GTP→ BH4
M00843 L-threo-Tetrahydrobiopterin biosynthesis, GTP → L-threo-BH4
M00165 Reductive pentose phosphate cycle (Calvin cycle)
M00011 Citrate cycle, second carbon oxidation, 2-oxoglutarate→ oxaloacetate
M00841 Tetrahydrofolate biosynthesis, mediated by PTPS, GTP → THF
M00913 Pantothenate biosynthesis, 2-oxoisovalerate/spermine→ pantothenate
M00051 De novo pyrimidine biosynthesis, glutamine (+PRPP) → UMP
KEGG module category
CAG-345 sp000433315 (HRGMv2_3440)
Blastocystis hominis (GCA_039096935.1)
Combined
Benefited taxon
KEGG module completeness
KEGG module category
Metabolism of cofactors and vitamins
Lipid metabolism
Carbohydrate metabolism
Nucleotide metabolism
Amino acid metabolism
Biosynthesis of terpenoids and polyketides
Glycan metabolism
KEGG module completeness
100%
0%
Benefited taxon
CAG-345 sp000433315
Both
Blastocystis hominis
