## Supplementary material for "Identification of eukaryotes as a bycatch in global microbiome data": Figure S11

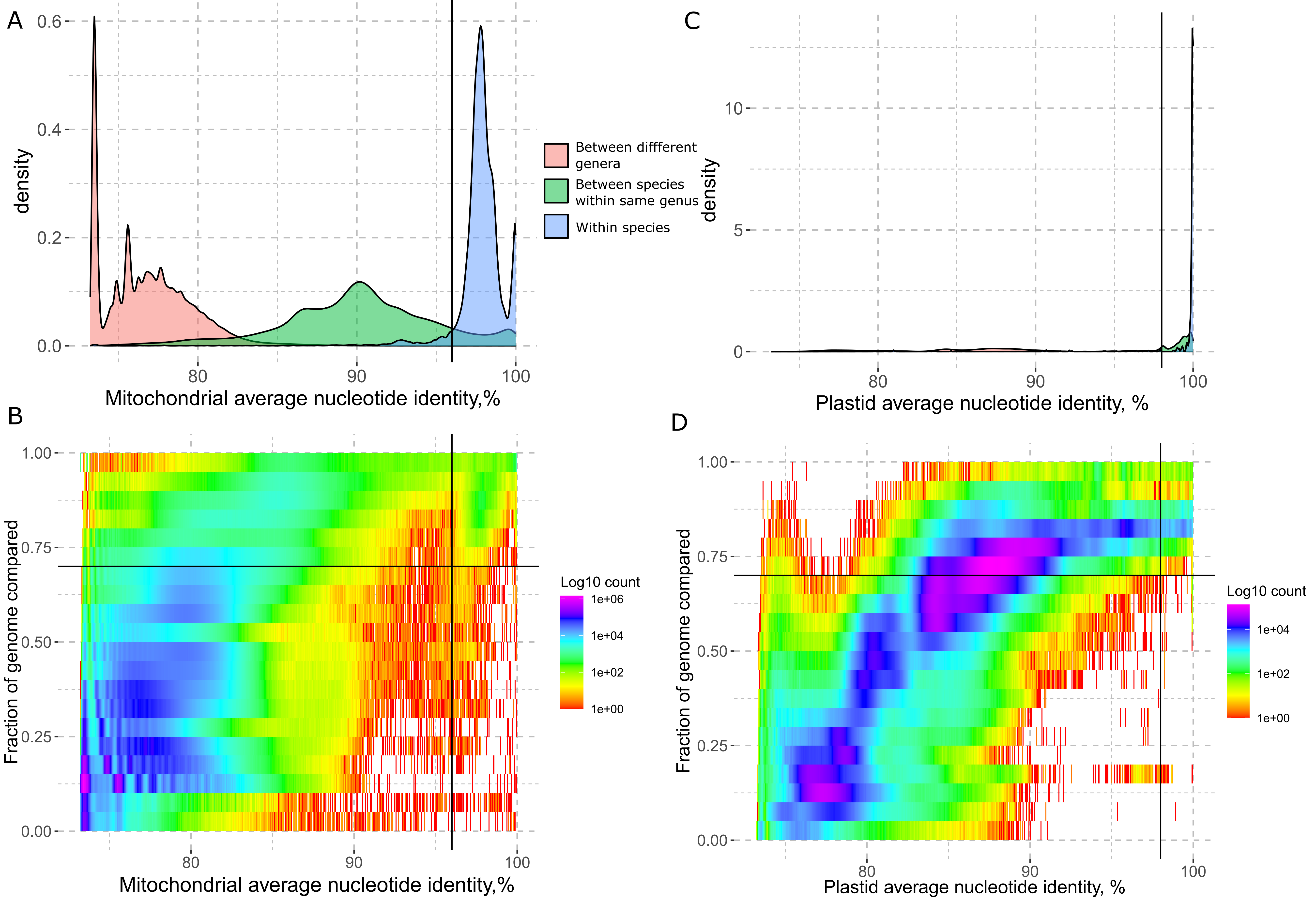

A
0.6
0.4
0.2
0.0
density
80
90
100
Mitochondrial average nucleotide identity,%
Between diffferent genera
Between species within same genus
Within species
C
10
5
0
density
80
90
100
Plastid average nucleotide identity, %
B
1.00
0.75
0.50
0.25
0.00
Fraction of genome compared
80
90
100
Mitochondrial average nucleotide identity,%
Log10 count
1e+06
1e+04
1e+02
1e+00
D
1.00
0.75
0.50
0.25
0.00
Fraction of genome compared
80
90
100
Plastid average nucleotide identity, %
Log10 count
1e+04
1e+02
1e+00
